# Generating Structural Ensembles of Disordered Proteins with Diffusion Models

**DOI:** 10.64898/2026.10.02.756250

**Authors:** Olivier Trottier, Jack H. Gwozdecky, Sarah Rauscher

**Affiliations:** Department of Chemical and Physical Sciences, University of Toronto Mississauga, Mississauga, ON, Canada; Department of Physics, University of Toronto, Toronto, ON, Canada; Acceleration Consortium - University of Toronto, ON, Canada; Department of Chemistry, University of Toronto, Toronto, ON, Canada

## Abstract

Generative models of protein structures have accelerated scientific discovery. However, the leading structure-prediction models are trained primarily on crystal structures, limiting their ability to sample the conformations of intrinsically disordered proteins (IDPs). All-atom molecular dynamics (MD) simulations can probe the diversity of IDP conformational ensembles, but these simulations are slow and costly. Leveraging the speed of generative models and the accuracy of MD, we develop Zephyr, an SE(3) equivariant diffusion model trained on simulated IDP ensembles. The model generates the structure of a single protein sequence in two phases: a coarse-grained structure is first sampled and heavy atoms are subsequently denoised onto this structure. We evaluate our method on two fronts: ensemble accuracy and sampling efficiency. Across five test systems, the model reproduces key geometric and energetic metrics of the MD ensembles, while sampling at least 63 times faster.

---

Intrinsically disordered proteins (IDPs) and disordered regions (IDRs) lack a single fixed structure, and they are prevalent in the human proteome: only *∼* 37% of proteins are fully folded, *∼* 58% include both folded domains and IDRs, and *∼* 5% are entirely disordered.^1^ IDPs and IDRs perform essential roles, including regulating the cell cycle, transcription and translation, and acting as hubs in signaling networks,^2^ and they are implicated in diseases including Alzheimer’s, Parkinson’s and cancer.^3^ Their defining feature is also their central difficulty: rather than a single structure, IDPs and IDRs populate ensembles of interconverting conformations. This heterogeneity makes them challenging to study experimentally, as most techniques report ensemble-averaged observables that cannot uniquely define the underlying conformational ensemble.^4^ All-atom molecular dynamics (MD) simulations can resolve individual conformations, but exhaustive sampling of disordered ensembles remains computationally challenging, motivating the development of faster sampling and ensemble generation methods.^5–14^

Recently, machine learning (ML) models have made remarkable progress in learning the sequence-structure relationship of proteins, ^15–20^ but capturing the diversity of conformational ensembles remains a challenge. ^11,21^ ML approaches have also had success in reproducing MD ensembles and generally employ one of two strategies: force learning or ensemble learning. In the first approach, models are trained to emulate MD simulators. ^22–24^ In the second approach, models learn the structural distribution directly from data.^5,25,26^ Although the learning difficulty of the second approach can be higher, its sampling efficiency is not limited by energy barriers.

In this work, we follow the second approach and develop Zephyr. Specifically, we train lightweight (*<* 1M parameters) diffusion models with an equivariant graph neural network architecture to reconstruct the conformational ensemble of disordered proteins after training on MD-generated structures. To simplify the learning task, models are trained on the structural ensemble of a single sequence. To assess accuracy, ensembles of five systems that span various levels of disorder, force fields and sequence lengths are compared on structural and energy-based metrics. Finally, to assess efficiency, the sampling speed of the model is compared against MD.

Zephyr is inspired by the success of generative models for protein structures^16,19^ and, in the specific case of IDPs, it builds upon previous works ^5,26–31^ with the following novelties and methodological choices: 1) Zephyr generates all-atom structures of a single sequence after training directly on the target sequence’s MD ensemble; 2) the quadratic complexity scaling that plagues all-atom graph neural networks is avoided by separating the structure generation into two steps that focus on different spatial resolutions and are solved by independent models; 3) techniques suggested by Karras et al.^32^ to improve the training and sampling of diffusion models for image generation are adapted to structure generation; 4) the conditioner framework of Chroma^16^ is used to steer denoising trajectories towards physically relevant structures. Overall, our results demonstrate that lightweight single-sequence models can learn the diverse conformational ensembles of disordered proteins, laying the foundation for building more general models of IDPs and IDRs.

## Results

### Overview

Our proposed method for sampling the conformational ensemble of a given protein of interest consists of three phases: 1) dataset building, 2) model training and 3) ensemble generation (Figure 1 and Supplementary Notes A to C). In the first phase, a set of all-atom MD trajectories of a given protein is generated with a given force field. In the second phase, two SE(3) equivariant diffusion models are trained to denoise structures gathered from MD trajectories. One model generates coarse-grained structures, while the second model generates positions of the heavy atoms (all atoms excluding hydrogen) conditioned on a given coarse structure. In the third phase, heavy-atom positions of the protein are generated with the trained models: the coarse-structure model generates positions of the coarse-grained beads, which are used as input by the fine-structure model to sample the heavy-atom positions. In addition, the heavy-atom model sampling process is steered by physical conditioners to prevent atom clashes and unphysical bond geometries. After sampling, hydrogen atoms are added using GROMACS’ `pdb2gmx` to produce an all-atom structure. Finally, samples are pruned based on their force-field potential energies in comparison to the training dataset energies.

**Figure 1:**
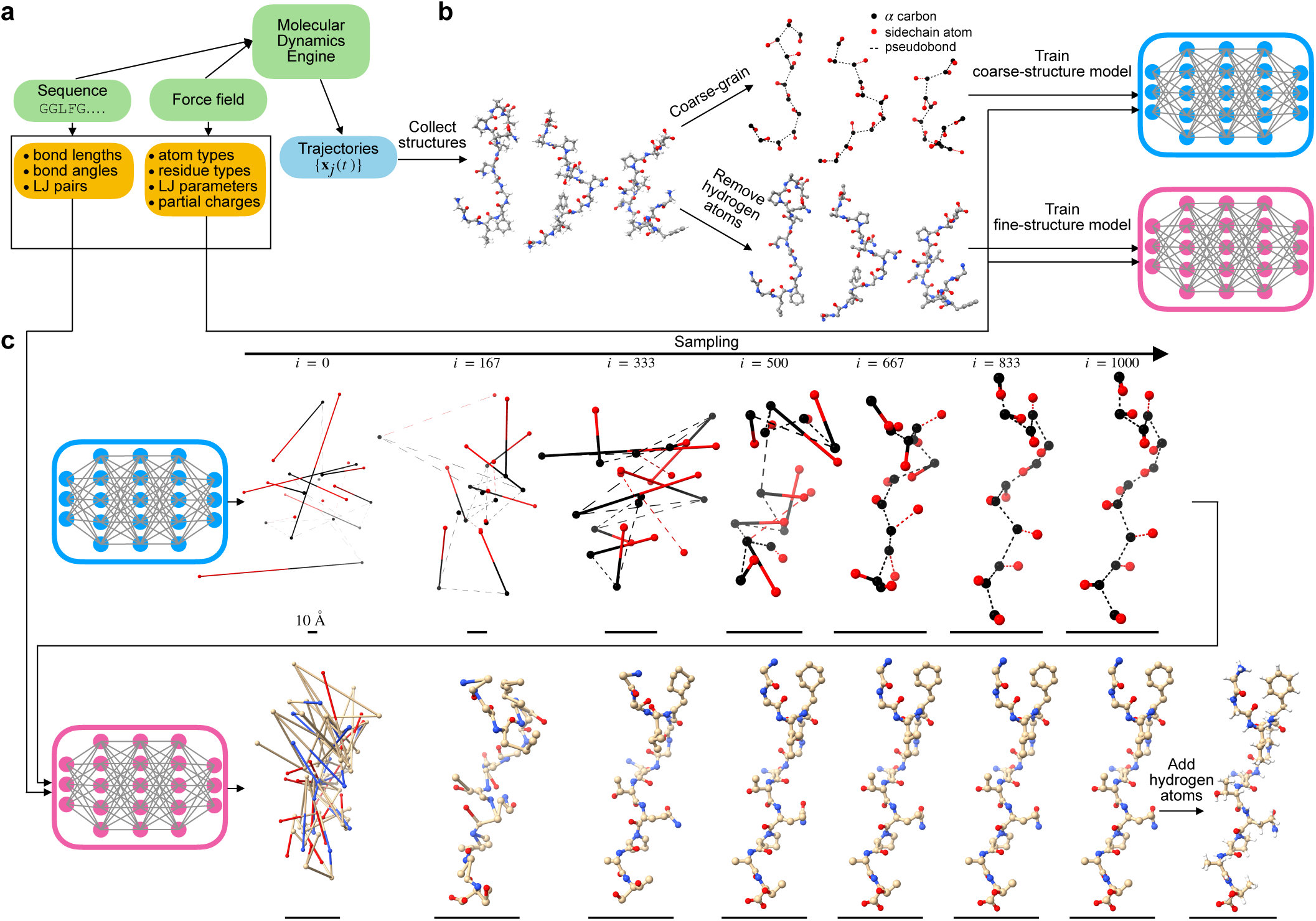
Summary of sampling with Zephyr. (**a**) Phase 1: Build MD Datasets. MD trajectories are obtained for a given protein sequence, force field and MD engine (green boxes). Physical and chemical properties are then derived from these choices (yellow boxes) and used by both models during training and sampling. (**b**) Phase 2: Train Models. Two diffusion graph neural networks are trained on structures from MD trajectories. The first model (blue) is trained on coarse-grained structures and aims to capture large-scale features of the structural ensemble. The second model (pink) is trained on heavy-atom structures and aims to learn small-scale features using local message-passing layers whose neighborhoods are defined by a given coarse-grained structure. (**c**) Phase 3: Generate Ensemble. The coarse-structure model is first used to generate the positions of the coarse-grained beads, which are then given to the fine-structure model as input to sample the heavy-atom positions. Snapshots of denoising trajectories for models trained on the Nup98-12 system are shown, where *i* labels the denoising step. Finally, hydrogen atoms are added using GROMACS’ `pdb2gmx`. Black scale bars correspond to 10 Å. 3D protein models are made with ChimeraX. ^33^ See Supplementary Movies 1–10 for examples of denoising trajectories for all five systems studied in this work.

### Model Architecture

Diffusion models are trained to reverse a diffusion process that gradually adds noise to a given object.^34,35^ Our work focuses on denoising a set of 3D atomic coordinates **x** representing either a coarse-grained or heavy-atom protein structure. During training, a denoiser *D_θ_*(**x***, t*) is trained to remove noise from noisy atom coordinates for a range of noise amplitudes. Once trained, sampling is achieved by applying the model autoregressively on pure noisy coordinates until all noise is removed.

The denoiser *D_θ_*(**x***, t*) is modeled using a graph neural network (GNN) with SE(3) equivariant message-passing layers. Each layer consists of an equivariant graph convolution (EGC) followed by a cross-product module (CPM) (Figure 2a, Supplementary Note C).

**Figure 2:**
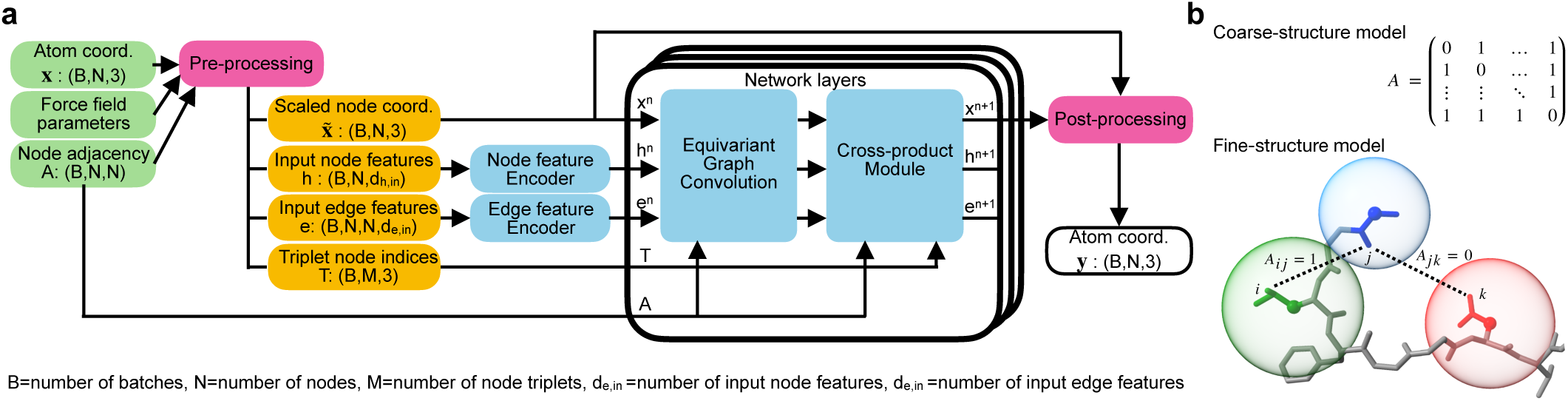
Architecture of the graph neural network. (**a**) The model’s input is a molecular graph whose node features correspond to noisy atom coordinates **x**. Graph edges are encoded in the adjacency matrix *A* and indicate atoms that exchange features in message-passing layers. Additional features are defined in the preprocessing step, which depends on the structure type (coarse-grained or heavy-atom). After passing through encoders, node and edge features are updated iteratively with a sequence of layers. The final layer’s output is post-processed with noise-schedule-dependent functions to yield the denoised atom coordinates **y**. All blue boxes represent trainable modules while magenta boxes are fixed operations. See Supplementary Note C for details. The equivariant graph convolution updates node coordinates based on a weighted sum of the coordinates of neighboring nodes using SE(3) invariant weights. Node and edge features are updated with conventional message-passing layers. The cross-product module further updates node positions by aggregating local cross-products of the node coordinates (SE(3) equivariant). The triplets of node indices that form the cross-products are determined a priori. The shape of tensors is indicated in parentheses. (**b**) Definition of graph topologies. Coarse-grained bead graphs are fully connected, i.e. the adjacency matrix *A* is full. For heavy-atom graphs, two nodes are adjacent when their respective coarse-grained beads are adjacent. Beads have a fixed radius, and their intersection, within a buffered distance, implies adjacency (Supplementary Note A). In the depicted example, all blue elements are in contact with all green elements, while red elements are not adjacent to either blue or green elements. The 3D protein model was made with ChimeraX.^33^

The denoiser’s inputs are the following: (1) *{***x***_i_}*, a set of noisy atom positions defining a given protein structure; (2) *S*, the amino acid sequence; (3) *t*, the diffusion time, which informs the denoiser about the input’s noise level; (4) *A_ij_ ∈ {*0, 1*}*, the adjacency matrix, where *A_ij_* = 1 indicates that atoms *i* and *j* exchange messages; (5) **p**_ff_, force field parameters; and (6) *T*, the set of triplet node indices *T* = *{*(*i, j, k*) : *i, j, k ∈ I}* used by the CPM. Input node and edge features are derived from these inputs (Supplementary Note C.1), and are subsequently passed to trainable encoders (Supplementary Note C.2).

The EGC layer updates node features, positions and edge features using residual connections. The node feature and edge feature updates are performed using message-passing layers with attention-like weights, while the position updates are calculated using a weighted sum of edge vectors (Supplementary Note C.3).

The CPM updates node positions by aggregating cross-products of neighboring node positions defined by a predetermined index set of node triplets *T* (Supplementary Note C.4). Since the CPM is not invariant to reflections (**x** *→ −***x**), the model is sensitive to chiral features, such as dihedral angles, allowing it to correct errors in these features during denoising (Supplementary Note E.7).

Finally, the output of the last layer is post-processed to implement an SE(3) equivariant rescaling that accelerates training by controlling the output’s variance (Supplementary Note C.5). A summary of the model’s hyperparameters is given in Supplementary Table 3.

### Coarse-Graining

Our approach consists of factoring the conformational ensemble into two parts and training models independently to learn the distribution of each factor:

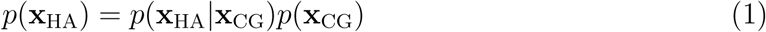

The coarse-structure model is tasked with learning the distribution of the coarse-grained beads’ positions (*p*(**x**_CG_)), while the fine-structure model’s goal is to accurately position heavy atoms given the coarse-grained bead positions (*p*(**x**_HA_|**x**_CG_)).

The coarse structure represents each residue using two beads: one centered at the backbone C*α* atom and one centered at a side-chain atom, each with a predetermined radius (Figure 2b). The positions and radii of these beads are determined by analyzing pairwise atomic distances in a curated set of PDB structures (Supplementary Note A). The fine structure represents all heavy atoms.

The main purpose of separating the structure generation into two steps is to reduce the computational complexity of the heavy-atom model. Ideally, all nodes should exchange messages to account for all possible pairwise interactions. However, this operation would scale quadratically with the number of atoms, preventing applications to large systems. Our approach avoids this by restricting message passing to pairs of heavy atoms whose respective coarse-grained beads are in contact (Supplementary Note A). Consequently, the computational complexity of the fine-structure model scales sub-quadratically with the system size since the positions of heavy atoms are influenced only by neighboring atoms, as defined by the associated coarse-grained bead positions (Supplementary Note A.2). Therefore, the fundamental assumption of our approach is *locality*, i.e. that, given the coarse-grained structure, the sub-angstrom precision of atom positions is determined by local interactions.

### Assessment of Ensemble Accuracy

Unlike generative models trained on large protein databases, Zephyr trains models on a *single* protein sequence. To assess the robustness of our approach across different proteins, we consider five test systems that span a range of disorder and sequence lengths and were simulated with different force fields (Supplementary Note D). Specifically, models are trained on a short (Nup98-12) and a long (Nup98-24) model sequence of an FG-nucleoporin,^36^ the 10-residue protein CLN025,^37^ a 24-residue IDP containing RS motifs (RS peptide) ^38^ and the sequence `AAQAA` repeated three times (`(AAQAA)`_3_).^39^

To evaluate how accurately our approach reproduces the MD ensemble, each model ensemble is compared against its training and test sets (Supplementary Table 7). The model– test differences quantify both model errors and the finite sampling of the MD ensembles, while the train–test differences quantify the quality and extent of conformational sampling in the MD simulations. Model–test differences that exceed the train–test differences are, therefore, interpreted as model errors.

Ensembles are evaluated by comparing several 1D and 2D distributions of a set of metrics that capture both coarse- and fine-grained structural features. In addition to using structural metrics (e.g. radius of gyration, hydrogen bonds), we use metrics that are derived from the geometric quantities used in all-atom biomolecular force fields. Furthermore, ensembles are assessed using selected terms of the force field potential energy function that only involve protein atoms (Supplementary Note E.2). Since these energy terms depend on higher powers of an associated geometric quantity (e.g. bond energies scale with the squared deviation of the bond length from its reference value), they assess higher moments of the geometric quantity distribution. As such, these energies are general measures of dispersion that quantify a given ensemble’s variability, where a higher energy value implies that the associated geometric quantity is further from the reference as determined by the force field. The fine structure is assessed with the distributions of bond lengths, bond angles and dihedral angles defined by the force field of the given system (Figure 3). During training, we also used these metrics to detect model failures and suggest improvements (see Methods – Training). As all bond length and bond angle distributions are unimodal, the Z-score of each defined feature is used to standardize the values, where the reference mean and standard deviation are estimated using the training dataset (Supplementary Note E.1). To compare two ensembles, Z-scores of all features of the same type (bond length or bond angle), across all structures in a given ensemble, are combined into a single distribution. For all systems, the model accurately reproduces bond lengths based on comparison of bond length Z-score and bond energy distributions with the MD reference ensembles. While the bond angle Z-score distributions are largely consistent between model and MD ensembles, the right-shifted bond angle energy distributions of some model ensembles (Nup98-12, Nup98-24) indicate oversampling of less probable bond angles compared to the corresponding MD ensembles. The dihedral angle energy distributions for the model ensembles are also slightly right-shifted relative to those from MD, except for the Nup98-12 and CLN025 systems. This indicates that the model generally succeeds at learning the dihedral energy surface of the underlying force field but sometimes oversamples improbable angles relative to MD.

**Figure 3:**
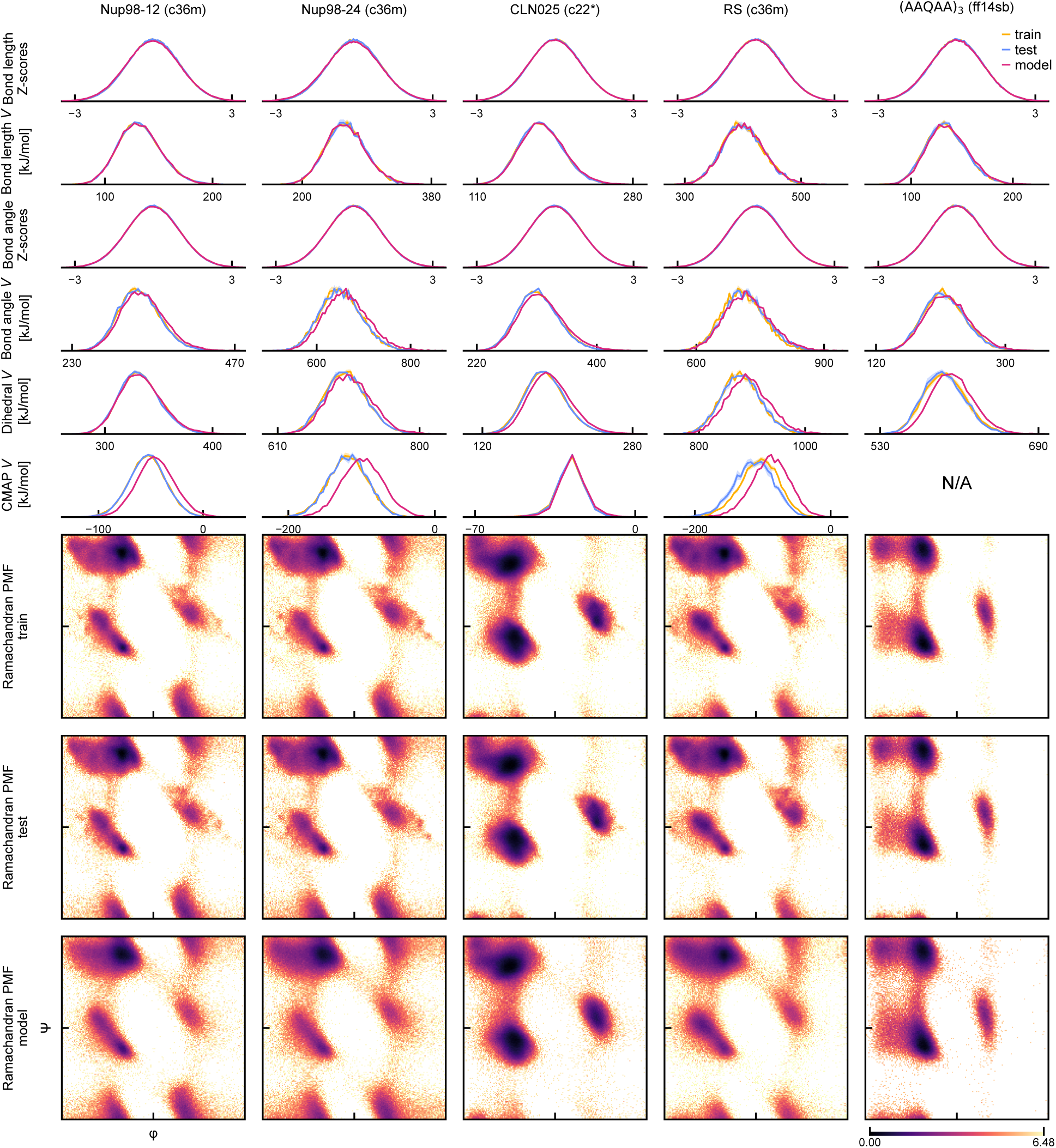
Fine Structure Assessment. Probability densities of several metrics (rows), including Z-scores and potential energies, are shown for all studied systems (columns). Supplementary Note E details how these metrics were computed. The distributions for the model, train and test sets are shown in each case. The train and test distributions are averaged over independent MD trajectories, with the shaded area indicating the standard error of the mean. Note that Amber14sb has no CMAP correction. The last three rows show the Ramachandran potential of mean force (PMF), *E*_Rama_(*ϕ, ψ*) = *−* log *P* (*ϕ, ψ*), shifted such that the global minimum is zero for the model, train and test sets. Backbone dihedral angles *ϕ, ψ* span the range [*−*180*^◦^,* 180*^◦^*].

Backbone dihedral angles, which report on the protein’s secondary structure, are also included in the fine structure assessment. For all five systems, model ensembles exhibit broader and smoother sampling of Ramachandran basins relative to MD. For systems simulated with CHARMM36m,^40^ this is also evidenced by model CMAP energy distributions that are right-shifted compared to the associated MD distributions. The same right-shift is not seen for the CLN025 system, which was simulated with CHARMM22*,^41^ as this force field only has a CMAP correction for glycine and proline residues.

Coarse structure is assessed using the radius of gyration (*R_g_*), hydrogen-bond contact maps, and Lennard-Jones and Coulomb energy distributions (Figure 4). The overall features of the hydrogen-bond contact maps are preserved in the model ensembles across all studied systems. However, the frequency of hydrogen bonds is lower in the model ensembles compared to the MD ensembles. These differences are observed across all systems, with CLN025 showing the smallest difference in hydrogen bond frequency. The *R_g_* distributions of the model ensembles are consistent with those of MD, except for the RS peptide. For this system, the model ensemble’s distribution is more similar to the training set than to the test set. This discrepancy likely reflects under-sampling, as indicated by the difference between the training and test distributions for the RS peptide. Regarding the Lennard-Jones energy, the Nup98-12 system shows the best agreement between the model and MD distributions, but all systems exhibit discrepancies. The Coulomb energy distributions are generally well-sampled by the model, with the Nup98-24 and `(AAQAA)`_3_ systems showing small discrepancies.

**Figure 4:**
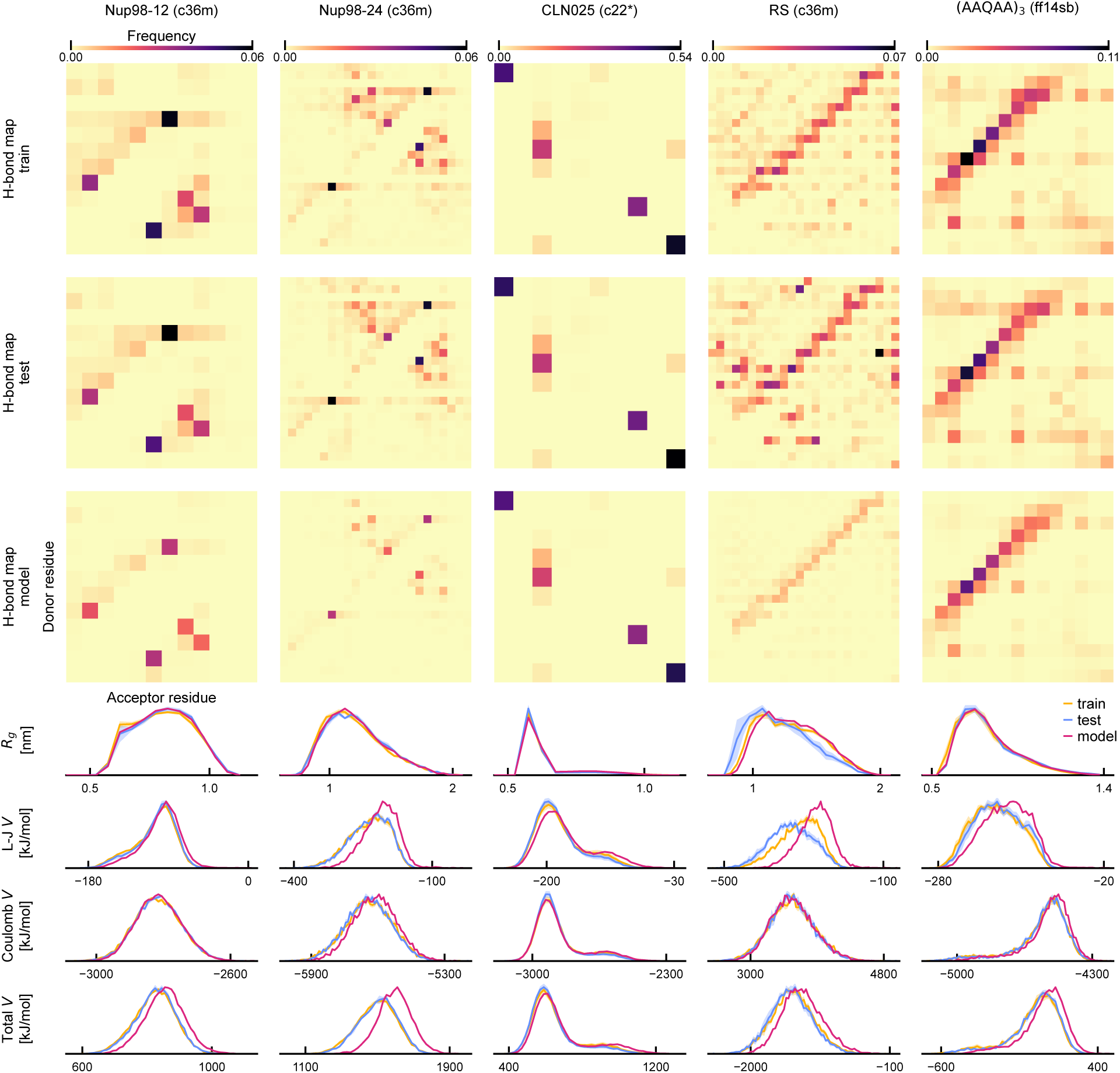
Coarse Structure Assessment. Hydrogen-bond contact maps are shown for the model, train and test sets of all systems (upper three rows). The color of each square in these maps indicates the probability of a hydrogen bond between a pair of residues. In the lower four rows, one-dimensional distributions compare model, training and test set distributions as indicated in the legend. Distributions for the training and test sets are averaged over independent MD trajectories, and the standard error of the mean is shown (shaded area).

Among all systems, the model ensemble for CLN025 exhibits the best agreement across all observables with the MD ensemble, consistent with CLN025 having the largest training dataset. To further test Zephyr’s ability to accurately predict the ensemble of this system, we quantified the sampling of the CLN025 folded and unfolded states (Supplementary Note F.5). These states are sampled in the same proportion as MD.

To assess the accuracy and variability of all geometric features, the distribution of the total potential energy defined by the force field and involving only protein atoms is compared between model and MD ensembles (Figure 4, last row). In general, model ensembles exhibit greater variability than MD ensembles, with the CLN025 model distribution most closely resembling that of MD.

Overall, the model reproduces the fine structure more accurately than the coarse structure (comparing Figure 3 and Figure 4). Two reasons may explain why the coarse structure is more difficult to learn. First, the model may be under-trained, resulting in structures with higher potential energies due to model errors. Second, the additional equilibration introduced by the *ψ* parameter during sampling may not reweight the conformational distribution with the same weight as the Boltzmann distribution associated with the force-field potential energy. However, additional equilibration during coarse structure sampling does not resolve these discrepancies (Supplementary Fig. 6).

In addition to comparing the coarse and fine structure, 2D energy surfaces of UMAP^42^- embedded properties are compared to assess the sampling of the conformational landscape (Supplementary Note E.5). In general, model surfaces retain the global structure of MD surfaces, but exhibit local discrepancies that are also present between the train and test MD datasets.

We next assess the effect of the force field on the ensemble learning difficulty by comparing two models trained on structures of Nup98-12 generated with CHARMM36m^40^ and Amber14sb,^43^ respectively (Supplementary Note E.8). We hypothesized that the CMAP of the CHARMM36m force field increases the difficulty of learning the Ramachandran map because it results in more local free energy minima. Contrary to our hypothesis, the force field has no significant effect on the learning difficulty of the fine structure (Supplementary Fig. 11a). However, the coarse structure of the Nup98-12 ensemble generated with Amber14sb is more difficult (Supplementary Fig. 11b). Thus, the ability of the model to learn the ensemble depends on force field.

Finally, foreseeing applications that require strict reproduction of the features of the MD ensembles, the value of a short MD equilibration after generation is assessed (Supplementary Note E.9). An assessment of the fine (Supplementary Fig. 12) and coarse (Supplementary Fig. 13) structure demonstrates that equilibration generally reduces discrepancies between the model and MD ensembles, as expected, but at the cost of additional computation.

### Assessment of Sampling Efficiency

The efficiency of Zephyr in sampling a diverse set of conformations is assessed in several ways. First, the compute time needed to generate a fixed number of samples is benchmarked on a single workstation (8 CPUs, 1 GPU) (Figure 5a, Supplementary Note F.1). MD samples were recorded every 1 ns to mirror the dataset construction method (Supplementary Note D). The total sampling time of our method accounts for dataset preparation, input feature calculation and training of both models. Once training is complete, the model generates samples 63 to 165 times faster than MD sampling at 1 ns intervals (Supplementary Table 9). Note that the acceleration factor of the model compared to MD depends on the downsampling interval of the MD trajectories.

Second, the efficiency of the model in reproducing the MD distributions is assessed as a function of the number of samples *N* (Figure 5b). The similarity of a given ensemble to the MD ensemble is quantified with the Jensen-Shannon (JS) distance relative to the associated test ensemble (Supplementary Note F.2). The decay rate of the JS distance curve quantifies the efficiency of converging to the test ensemble. For each metric and system, the JS distance curves of model and training ensembles are compared. For bond lengths and bond angles, the model’s efficiency is similar to MD. For sampling Ramachandran maps and *R_g_*, the model is as efficient as MD for ensembles with *<* 2500 samples, but exhibits differences for larger ensembles.

Third, the discovery rate of unique backbone dihedral states is assessed and compared with MD (Figure 5c). The backbone dihedral state of a given structure is defined in Supplementary Note F.4. Across all systems, the model discovers unique dihedral states at a rate that is similar to or faster than MD.

Fourth, the training, sampling and evaluation methodology is repeated using a minimal amount of training data (train split *≤* 10%) (Supplementary Note F.3). The fine structure assessment demonstrates that the model reproduces the MD distributions, while the coarse structure assessment demonstrates increased variability in the model distributions as seen in the main (full dataset) assessments. Overall, this analysis shows that the model is able to generate conformational ensembles with a smaller amount of data than we initially anticipated, suggesting that a higher efficiency could be achieved by varying the amount of training data.

Finally, the ability of Zephyr to discover unseen conformational states is assessed using the CLN025 protein (Supplementary Note F.5). When trained on unfolded structures, a varying number of folded structures are sampled depending on the similarity of the training dataset to the folded state. When trained on folded structures, the model succeeds at generating unfolded structures only in one of the two tests that we conducted (Supplementary Fig. 18c). The best examples of model structures that demonstrate generalization are shown in Supplementary Fig. 18d. These results demonstrate that the model has some ability to sample beyond its training dataset.

## Discussion

Obtaining conformational ensembles of IDPs in atomistic detail is computationally intensive, with MD simulations being one of the most widely used methods for this purpose. Here, we introduce Zephyr, a sampling method that uses a generative model to construct the conformational ensemble of disordered proteins after training on MD-generated structures. The generation task is decomposed into two spatial resolution levels: coarse-grained beads and heavy atoms. Independent models with *<* 1M parameters are trained for each of these resolutions. Combined with a method to add hydrogen atoms, the resulting approach generates all-atom structures with a computational cost that scales sub-quadratically with the number of atoms.

Across the five model systems tested, the model reproduces many features of the reference MD ensembles as assessed by structural and energetic metrics. Agreement is strongest for local geometric features, including bond lengths, bond angles and dihedral angles, suggesting the model accurately recovers many features of the fine structure. However, some non-local features, such as hydrogen bonds, exhibit discrepancies compared to the MD ensembles, suggesting further improvements to the coarse-structure model are needed. Analysis of the effect of energy minimization and MD equilibration also supports this idea since it provides slight improvements in the accuracy of the fine structure, but more significant improvements in the coarse structure. Furthermore, our results demonstrate that the ability of models to learn fine and coarse features is force-field dependent.

In the initial conceptualization of this work, we set out to train a sequence-conditioned model to sample IDP conformational ensembles using MD training datasets. However, given the highly variable conformations of IDPs, we realized that training a generative model with MD accuracy would not be feasible with the amount of compute time and data at our disposal. We therefore focused on learning a single-sequence ensemble, as it is a prerequisite for multi-sequence learning. This allowed us to quickly iterate the method, develop efficient training and sampling strategies that are transferable to the multi-sequence task and discover failures of the model that would warrant further investigation. Furthermore, our tests with minimally trained models show that the amount of training data required to accurately sample the conformational ensemble is smaller than initially anticipated.

Although the specificity of trained models to one protein sequence is a limitation in the context of sampling the space of disordered proteins, we believe these models are already applicable to understanding disease-related IDPs. Indeed, for some diseases (e.g. Alzheimer’s), the conformational ensemble of a *single* IDP (e.g. amyloid-*β*) is of interest in drug discovery.^44,45^ In such cases, the generalizability of general-purpose protein language models is unnecessary. Training de novo models on a single-sequence dataset therefore allows one to trade off generalizability for a more accurate ensemble, at the cost of additional compute time. Such approaches would be desirable when general-purpose models predict low-confidence structures ^46^ and full all-atom MD simulation studies are too expensive.

In conclusion, we have demonstrated the feasibility of training lightweight generative models to sample single-sequence conformational ensembles using training datasets generated with MD simulations. Combining the accuracy of MD with the speed of generative models, Zephyr promises to accelerate computational studies of IDPs and facilitate the identification of conformational states relevant to drug development.

## Methods

### Diffusion Model Framework

Our approach uses the stochastic differential equation (SDE) formalism of the diffusion process to diffuse the 3D coordinates of atoms in a protein structure:

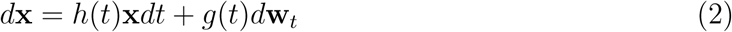

where **x** *∈* R*^N×^*^3^, *N* is the number of atoms, *d***w***_t_* is a standard Wiener process and *h*(*t*)*, g*(*t*) are predetermined functions of time *t ∈* [*t*_min_*, t*_max_]. The related probability density function *p*(**x***, t*) of obtaining **x** at time *t* admits a closed-form formula:

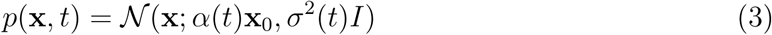

where *N* (**x**; *µ, σ*^2^) denotes a normal distribution with mean *µ* and variance *σ*^2^ and **x**_0_ denotes the initial (noiseless) coordinates. (*α*(*t*)*, σ*(*t*)) define what is commonly called the noise schedule, which determines the parametric functions *g*(*t*)*, h*(*t*) (Supplementary Note B.1).

Reversing the process described by eq. (2) yields another diffusion process given by the following SDE:^47,48^

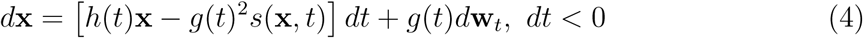

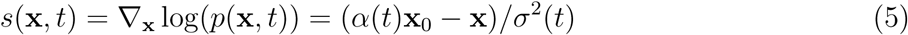

where *s*(**x***, t*) is the score function of *p*(**x***, t*). Therefore, starting from a set of noisy coordinates **x**(*t*_max_), integrating eq. (4) generates a denoising trajectory that converges to the initial noiseless coordinates **x**_0_ (Figure 1c). Generating this trajectory and saving its final point is the essence of the sampling process for diffusion models.

However, since *p*(**x***, t*) inherently depends on **x**_0_, the true reverse SDE cannot be used to generate data from pure noise. In practice, the reverse process is modeled using an estimate of **x**_0_:

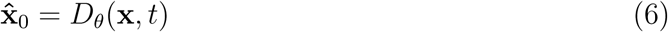

where *D_θ_*(**x***, t*) is a machine learning model parameterized by *θ*. *D_θ_*(**x***, t*) is called a denoiser as it learns to predict the initial noiseless data **x**_0_ given noisy data **x** at various times *t*. Given **x̂**_0_, sampling is achieved using the score function estimate *ŝ_θ_*(**x***, t*) calculated with eq. (5).

Finally, following Karras et al., we use a linear-in-time variance-exploding noise schedule since denoising gradients for this schedule point towards modes of the data distribution at all times *t*.^32^ The associated parametric functions of this noise schedule are:

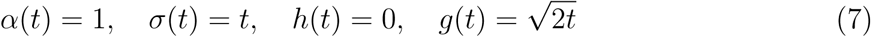

This schedule is used for both coarse-structure and fine-structure models, while its range is adjusted for each type of model to focus on relevant length scales for each type of structure (Supplementary Note B.2).

### Training

The denoiser *D_θ_*(**x***, t*) is trained by minimizing the squared error of the denoised coordinates:

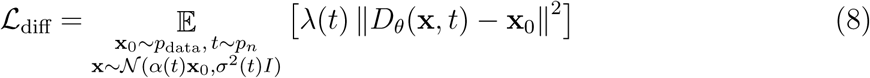

where *p*_data_ denotes the training data distribution, *p_n_* denotes the denoising time distribution, and *λ*(*t*) is a time-dependent weight used to equalize the denoiser error across all times (Supplementary Note B.2).

In addition to monitoring the diffusion loss, the model’s accuracy was tracked every 50 epochs of training by analyzing an ensemble of 100 structures generated with 100 denoising steps. Specifically, the mean and standard deviation of several metric distributions of the 100-structure sample were compared with the respective statistics calculated with the train and validation dataset splits (Supplementary Note B.3). Tracking statistics of the structural and energy metric distributions using a 100-structure ensemble allowed us to monitor their convergence, which is not evidently manifested in the diffusion loss curve.

In addition to tracking metric distributions during training, a detailed assessment of the fine structure and coarse structure was performed between training runs. These assessments were used to understand model failures and suggest improvements during model development. Specifically, in addition to the metrics that are presented above, the distributions of each bond length, bond angle and dihedral angle in the model ensemble were compared with those of the MD ensemble. For the sake of brevity, we do not present these individual comparisons but instead rely on metrics that agglomerate these geometric features.

Finally, in addition to tracking the validation loss, model overfitting was assessed with the distribution of the minimum train root mean square deviation (RMSD) RMSD_train, min_, the smallest heavy-atom RMSD of a given structure found by comparing it to all training dataset structures (Supplementary Fig. 9 and Supplementary Note E.6). For all systems, the model distributions of RMSD_train, min_ are similar to the test set distributions, indicating that model structures are as similar to the training set as the test set structures. In other words, the model does not memorize the structures of the training dataset.

### Sampling

Once models are trained, the reverse-diffusion SDE (eq. (4)) is used to generate a protein structure (coarse or fine) using an initial set of positions sampled from an isotropic normal distribution *N* (**x**; 0*, σ*^2^(*t*_max_)*I*). When sampling coarse structures, the denoising gradient depends entirely on the model output **x̂**_0_ through the score function estimate *ŝ*(**x***, t*). However, when sampling heavy-atom structures, additional restraints on the sampled geometries are applied to guide the denoising trajectories towards physically relevant landscapes. These restraints are implemented by substituting *p̂*(**x***, t*) with conditional marginal distributions *p̂*(**x***, t|C*), where *C* represents a condition on the geometry of the structures. As a consequence of Bayes’ theorem, the score of *p̂*(**x***, t|C*) is conveniently expressed as the sum of the unconditioned score function and a conditioning gradient:

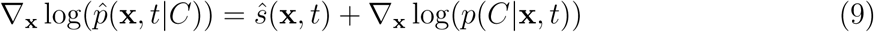

Four kinds of geometric features are restrained when sampling heavy-atom structures: 1) coarse-grained bead size, 2) bond length, 3) bond angle and 4) Lennard-Jones pair distance. These restraints are implemented by limiting the range of specific pairwise distances within a 5-*σ* level that is evaluated with statistics of the training dataset (Supplementary Note B.5). Following the sampling strategy used in the Chroma model,^16^ the diffusion SDE is modified to increase the equilibration of the marginal distributions *p̂*(**x***, t*). The additional equilibration has the effect of rescaling two terms in the original SDE:

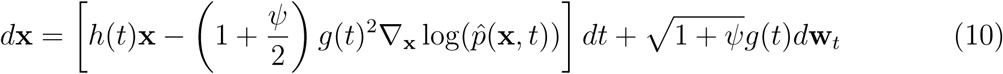

where *ψ ≥* 0 controls the amount of equilibration with *ψ* = 0 corresponding to the original SDE. A value of *ψ* = 10.0 is used to generate coarse structures, while *ψ* = 2.0 is used to sample fine structures (Supplementary Note B.4).

Finally, after generating a set of samples, structures that possess non-physical potential energies are removed. Specifically, the energy Z-scores of each structure are calculated with statistics of the training dataset. Then, structures that have an absolute Z score of 5 or more for any of the assessed energies are removed (Supplementary Note B.6). In all ensembles that were generated, this pruning process removes *<* 1% of the samples (Supplementary Table 2).

### Datasets

Our method is assessed on five systems that span various levels of disorder, force fields and sequence lengths (10 to 24 residues): 1) a 12-residue protein representing a model sequence of an FG-nucleoporin (Nup98-12),^36^ 2) a 24-residue protein whose sequence is that of Nup98-12 repeated twice (Nup98-24), 3) a 15-residue protein that consists of the sequence `AAQAA` repeated three times (`(AAQAA)`_3_),^39^ 4) a 24-residue protein that has several `RS` motifs^49^ and 5) CLN025, a 10-residue protein.^50^ The first four systems are chosen for their high degree of disorder, while the CLN025 system is used to test the model’s ability to generalize beyond its training data. Datasets of the first four systems were constructed from MD trajectories generated with GROMACS, and the CLN025 structures were obtained from the trajectory data of Lindorff-Larsen et al.,^50^ provided by D.E. Shaw Research (Supplementary Note D). Across all systems, three different force fields are probed: CHARMM22*,^41^ CHARMM36m^40^ and Amber14sb.^43^

## Supporting information

Supplementary Movie 1: nup98_12 coarse structure denoising trajectory

Supplementary Movie 2: nup98_12 fine structure denoising trajectory

Supplementary Movie 3: nup98_24 coarse structure denoising trajectory

Supplementary Movie 4: nup98_24 fine structure denoising trajectory

Supplementary Movie 5: CLN025 coarse structure denoising trajectory

Supplementary Movie 6: CLN025 fine structure denoising trajectory

Supplementary Movie 7: RS coarse structure denoising trajectory

Supplementary Movie 8: RS fine structure denoising trajectory

Supplementary Movie 9: AAQAA3 coarse structure denoising trajectory

Supplementary Movie 10: AAQAA3 fine structure denoising trajectory

## Data Availability

MD trajectories and model datasets generated in this work are available on Zenodo (https://doi.org/10.5281/zenodo.20492056).

## Code Availability

The codebase for Zephyr is open source and available on GitHub (https://github.com/ rauscher-lab/zephyr).

## Acknowledgments

S.R. acknowledges support from the Natural Science and Engineering Research Council of Canada (Discovery Grant no. RGPIN-201806408), Calcul Québec, and the Digital Research Alliance of Canada (alliance.can.ca). O.T. was supported by an Acceleration Consortium (AC) postdoctoral fellowship, and the AC’s Tacozoid cluster was used to carry out this research. S.R. and O.T. also acknowledge the support of an NVIDIA Academic Grant.

## Author Contributions

O.T., J.G. and S.R. conceptualized the research. O.T., J.G. and S.R. designed the methodology. O.T. and S.R. acquired funding. O.T. and J.G. carried out the MD simulations. O.T. designed and trained the ML models. O.T. and J.G. built analytical tools and designed the figures. S.R. supervised the research. O.T., J.G. and S.R. analyzed the data, wrote and revised the manuscript.

---

**Figure 5:**
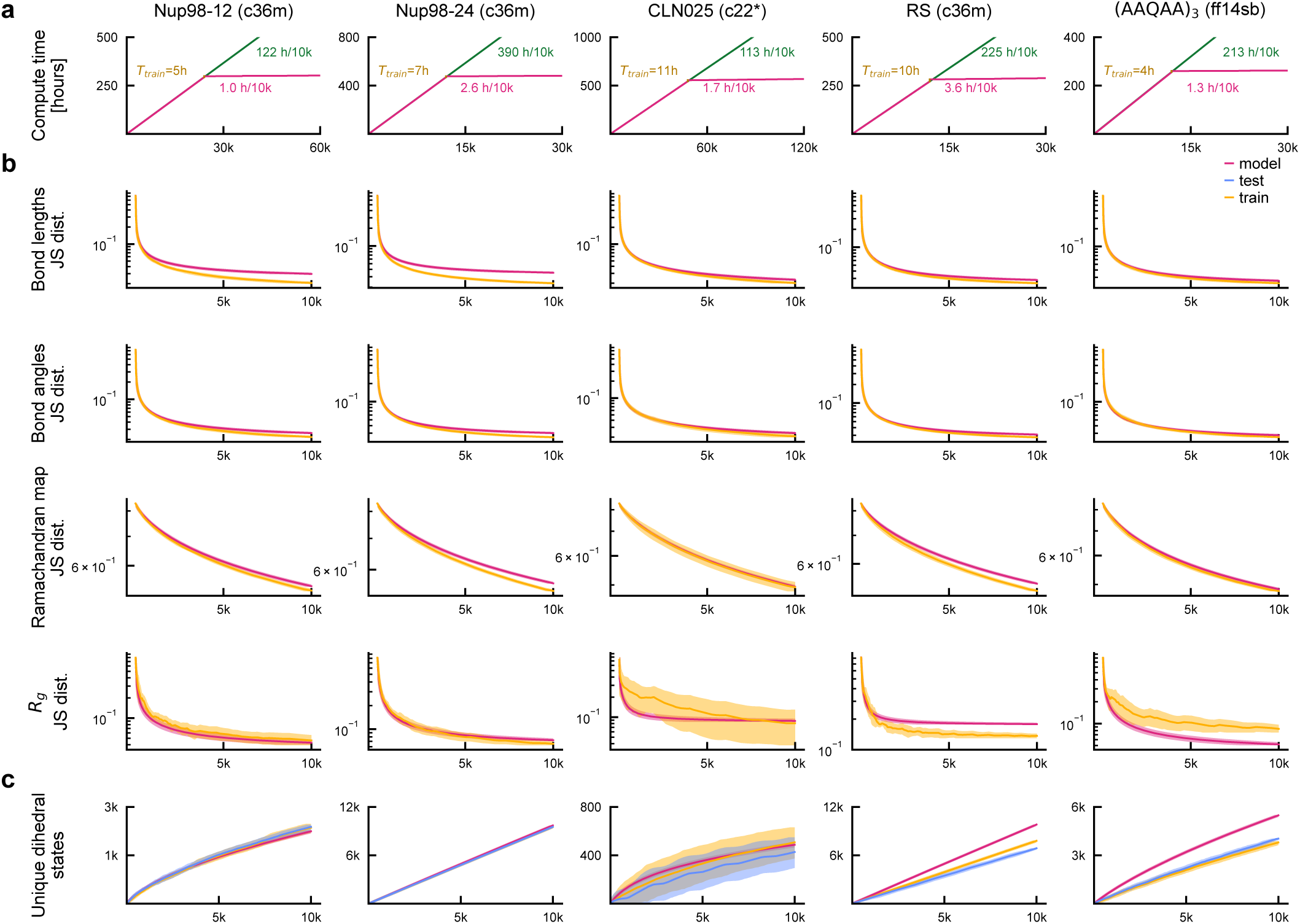
Sampling Efficiency. **a**. Compute time required to generate a given number of samples using MD (green) or the model (pink). For the MD curves, a new sample is counted after every 1 ns of simulation time. The model curves account for the dataset generation time (first phase), model training time *T*_train_ (second phase, vertical segment) and sampling (third phase). The MD and model curves are annotated with their slopes. **b**. For a given metric and system, the JS distance between a given ensemble subset and the respective test set is quantified as a function of the number of structures in the subset. The JS distance curves of model and train subsets are compared. For bond lengths and bond angles, the reported JS distance is an average over all the 1D marginal distances of each quantity. Figure legend applies to all panels in **b** and **c**. **c**. The number of unique backbone dihedral states (defined in Supplementary Note F.4) in a given subset is shown as a function of the number of structures in the subset. Model, train and test subsets are compared. Lines and shaded areas in **b** and **c** denote the bootstrap estimate of the means and standard errors (Supplementary Note F.2).

## References

(1) Tsang, B.; Pritišanac, I.; Scherer, S. W.; Moses, A. M.; Forman-Kay, J. D. Phase Separation as a Missing Mechanism for Interpretation of Disease Mutations. Cell 2020, 183, 1742–1756, DOI: 10.1016/j.cell.2020.11.050.

(2) Holehouse, A. S.; Kragelund, B. B. The Molecular Basis for Cellular Function of Intrinsically Disordered Protein Regions. Nature Reviews Molecular Cell Biology 2024, 25, 187–211, DOI: 10.1038/s41580-023-00673-0.

(3) Uversky, V. N.; Oldfield, C. J.; Dunker, A. K. Intrinsically Disordered Proteins in Human Diseases: Introducing the D2 Concept. Annual Review of Biophysics 2008, 37, 215–246, DOI: 10.1146/annurev.biophys.37.032807.125924.

(4) Mittag, T.; Forman-Kay, J. D. Atomic-Level Characterization of Disordered Protein Ensembles. Current Opinion in Structural Biology 2007, 17, 3–14, DOI: 10.1016/j.sbi.2007.01.009.

(5) Janson, G.; Valdes-Garcia, G.; Heo, L.; Feig, M. Direct Generation of Protein Conformational Ensembles via Machine Learning. Nature Communications 2023, 14, 774, DOI: 10.1038/s41467-023-36443-x.

(6) Tesei, G.; Trolle, A. I.; Jonsson, N.; Betz, J.; Knudsen, F. E.; Pesce, F.; Johansson, K. E.; Lindorff-Larsen, K. Conformational Ensembles of the Human Intrinsically Disordered Proteome. Nature 2024, 626, 897–904, DOI: 10.1038/s41586-023-07004-5.

(7) Zheng, S.; He, J.; Liu, C.; Shi, Y.; Lu, Z.; Feng, W.; Ju, F.; Wang, J.; Zhu, J.; Min, Y.; Zhang, H.; Tang, S.; Hao, H.; Jin, P.; Chen, C.; Noé, F.; Liu, H.; Liu, T.-Y. Predicting Equilibrium Distributions for Molecular Systems with Deep Learning. Nature Machine Intelligence 2024, 6, 558–567, DOI: 10.1038/s42256-024-00837-3.

(8) Gu, X.; Aranganathan, A.; Tiwary, P. Empowering AlphaFold2 for Protein Conformation Selective Drug Discovery with AlphaFold2-RAVE. eLife 2024, 13, RP99702, DOI: 10.7554/eLife.99702.

(9) Janson, G.; Feig, M. Generation of Protein Dynamics by Machine Learning. Current Opinion in Structural Biology 2025, 93, 103115, DOI: 10.1016/j.sbi.2025.103115.

(10) Aranganathan, A.; Gu, X.; Wang, D.; Vani, B. P.; Tiwary, P. Modeling Boltzmann-Weighted Structural Ensembles of Proteins Using Artificial Intelligence–Based Methods. Current Opinion in Structural Biology 2025, 91, 103000, DOI: 10.1016/j.sbi.2025.103000.

(11) Ghafouri, H.; Kadeřávek, P.; Melo, A. M.; Aspromonte, M. C.; Bernado, P.; Cortés, J.; Dosztányi, Z.; Erdős, G.; Feig, M.; Janson, G.; Lindorff-Larsen, K.; Mulder, F. A. A.; Nagy, P.; Pestell, R.; Piovesan, D.; Schiavina, M.; Schuler, B.; Sibille, N.; Tesei, G.; Tompa, P.; Vendruscolo, M.; Vondrasek, J.; Vranken, W.; Zidek, L.; Tosatto, S. C. E.; Monzon, A. M. Toward a Unified Framework for Determining Conformational Ensembles of Disordered Proteins. Nature Methods 2026, 23, 705–719, DOI: 10.1038/s41592-026-03003-2.

(12) Schnapka, V.; Morozova, T. I.; Sen, S.; Bonomi, M. Atomic Resolution Ensembles of Intrinsically Disordered Proteins with Alphafold. Nature Communications 2026, 17, 2399, DOI: 10.1038/s41467-026-69172-y.

(13) Tang, C.; Pandey, M. P.; Chen, C. G.; Megías, A.; Dehez, F.; Chipot, C. Breaking timescales with generative sampling of conformational transitions. Nature 2026, 657, 944–949, DOI: 10.1038/s41586-026-11025-1.

(14) Liu, Z. H.; Zhang, O.; De Castro, S.; Sun, K.; Ghafouri, H.; Attafi, O. A.; Fawzi, N. L.; Tosatto, S. C. E.; Monzon, A. M.; Moses, A. M.; Head-Gordon, T.; Forman-Kay, J. D. AlphaFlex: Ensembles of the human proteome representing disordered regions. bioRxiv, 2025; DOI: 10.1101/2025.11.24.690279.

(15) Dauparas, J.; Anishchenko, I.; Bennett, N.; Bai, H.; Ragotte, R. J.; Milles, L. F.; Wicky, B. I. M.; Courbet, A.; de Haas, R. J.; Bethel, N.; Leung, P. J. Y.; Huddy, T. F.; Pellock, S.; Tischer, D.; Chan, F.; Koepnick, B.; Nguyen, H.; Kang, A.; Sankaran, B.; Bera, A. K.; King, N. P.; Baker, D. Robust Deep Learning–Based Protein Sequence Design Using ProteinMPNN. Science 2022, 378, 49–56, DOI: 10.1126/science.add2187.

(16) Ingraham, J. B.; Baranov, M.; Costello, Z.; Barber, K. W.; Wang, W.; Ismail, A.; Frappier, V.; Lord, D. M.; Ng-Thow-Hing, C.; Van Vlack, E. R.; Tie, S.; Xue, V.; Cowles, S. C.; Leung, A.; Rodrigues, J. V.; Morales-Perez, C. L.; Ay-oub, A. M.; Green, R.; Puentes, K.; Oplinger, F.; Panwar, N. V.; Obermeyer, F.; Root, A. R.; Beam, A. L.; Poelwijk, F. J.; Grigoryan, G. Illuminating Protein Space with a Programmable Generative Model. Nature 2023, 623, 1070–1078, DOI: 10.1038/s41586-023-06728-8.

(17) Lin, Z.; Akin, H.; Rao, R.; Hie, B.; Zhu, Z.; Lu, W.; Smetanin, N.; Verkuil, R.; Kabeli, O.; Shmueli, Y.; Allan dos Santos Costa; Fazel-Zarandi, M.; Sercu, T.; Candido, S.; Rives, A. Evolutionary-Scale Prediction of Atomic-Level Protein Structure with a Language Model. Science 2023, 379, 1123–1130, DOI: 10.1126/science.ade2574.

(18) Watson, J. L.; Juergens, D.; Bennett, N. R.; Trippe, B. L.; Yim, J.; Eisenach, H. E.; Ahern, W.; Borst, A. J.; Ragotte, R. J.; Milles, L. F.; Wicky, B. I. M.; Hanikel, N.; Pellock, S. J.; Courbet, A.; Sheffler, W.; Wang, J.; Venkatesh, P.; Sappington, I.; Torres, S. V.; Lauko, A.; De Bortoli, V.; Mathieu, E.; Ovchinnikov, S.; Barzilay, R.; Jaakkola, T. S.; DiMaio, F.; Baek, M.; Baker, D. De Novo Design of Protein Structure and Function with RFdiffusion. Nature 2023, 620, 1089–1100, DOI: 10.1038/s41586-023-06415-8.

(19) Abramson, J.; Adler, J.; Dunger, J.; Evans, R.; Green, T.; Pritzel, A.; Ronneberger, O.; Willmore, L.; Ballard, A. J.; Bambrick, J.; Bodenstein, S. W.; Evans, D. A.; Hung, C.- C.; O’Neill, M.; Reiman, D.; Tunyasuvunakool, K.; Wu, Z.; Žemgulytė, A.; Arvaniti, E.; Beattie, C.; Bertolli, O.; Bridgland, A.; Cherepanov, A.; Congreve, M.; Cowen-Rivers, A. I.; Cowie, A.; Figurnov, M.; Fuchs, F. B.; Gladman, H.; Jain, R.; Khan, Y. A.; Low, C. M. R.; Perlin, K.; Potapenko, A.; Savy, P.; Singh, S.; Stecula, A.; Thillaisundaram, A.; Tong, C.; Yakneen, S.; Zhong, E. D.; Zielinski, M.; Žídek, A.; Bapst, V.; Kohli, P.; Jaderberg, M.; Hassabis, D.; Jumper, J. M. Accurate Structure Prediction of Biomolecular Interactions with AlphaFold 3. Nature 2024, 630, 493–500, DOI: 10.1038/s41586-024-07487-w.

(20) Xiao, Y.; Zhao, W.; Zhang, J.; Jin, Y.; Zhang, H.; Ren, Z.; Sun, R.; Wang, H.; Wan, G.; Lu, P.; Luo, X.; Zhang, Y.; Zou, J.; Sun, Y.; Wang, W. Protein Large Language Models: A Comprehensive Survey. Findings of the Association for Computational Linguistics: EMNLP 2025. 2025; pp 23080–23103, DOI: 10.18653/v1/2025.findings-emnlp.1255.

(21) Chakravarty, D.; Miller, J. J.; Teng, D.; Ramahi, Y. O.; Bryant, P.; Neira-Mahuzier, C.; Ramírez-Sarmiento, C. A.; Rauscher, S.; Bowman, G. R.; Tiwary, P.; Porter, L. L. Expanding Protein Structure Prediction into Conformational State Space. arXiv, 2026; DOI: 10.48550/arXiv.2608.02866.

(22) Majewski, M.; Pérez, A.; Thölke, P.; Doerr, S.; Charron, N. E.; Giorgino, T.; Husic, B. E.; Clementi, C.; Noé, F.; De Fabritiis, G. Machine Learning Coarse-Grained Potentials of Protein Thermodynamics. Nature Communications 2023, 14, 5739, DOI: 10.1038/s41467-023-41343-1.

(23) Jing, B.; Berger, B.; Jaakkola, T. AlphaFold Meets Flow Matching for Generating Protein Ensembles. Proceedings of the 41st International Conference on Machine Learning. 2024; Vol. 235; pp 22277–22303.

(24) Lewis, S.; Hempel, T.; Jiménez-Luna, J.; Gastegger, M.; Xie, Y.; Foong, A. Y. K.; Satorras, V. G.; Abdin, O.; Veeling, B. S.; Zaporozhets, I.; Chen, Y.; Yang, S.; Foster, A. E.; Schneuing, A.; Nigam, J.; Barbero, F.; Stimper, V.; Campbell, A.; Yim, J.; Lienen, M.; Shi, Y.; Zheng, S.; Schulz, H.; Munir, U.; Sordillo, R.; Tomioka, R.; Clementi, C.; Noé, F. Scalable Emulation of Protein Equilibrium Ensembles with Generative Deep Learning. Science 2025, 389, eadv9817, DOI: 10.1126/science.adv9817.

(25) Abdin, O.; Kim, P. M. Direct Conformational Sampling from Peptide Energy Landscapes through Hypernetwork-Conditioned Diffusion. Nature Machine Intelligence 2024, 6, 775–786, DOI: 10.1038/s42256-024-00860-4.

(26) Zhu, J.; Li, Z.; Zheng, Z.; Zhang, B.; Zhong, B.; Bai, J.; Hong, X.; Wang, T.; Wei, T.; Yang, J.; Chen, H.-F. Accurate Generation of Conformational Ensembles for Intrinsically Disordered Proteins with IDPFold. Advanced Science 2025, 12, e11636, DOI: 10.1002/advs.202511636.

(27) Taneja, I.; Lasker, K. Machine-Learning-Based Methods to Generate Conformational Ensembles of Disordered Proteins. Biophysical Journal 2024, 123, 101–113, DOI: 10.1016/j.bpj.2023.12.001.

(28) Zhu, J.; Li, Z.; Tong, H.; Lu, Z.; Zhang, N.; Wei, T.; Chen, H.-F. Phanto-IDP: Compact Model for Precise Intrinsically Disordered Protein Backbone Generation and Enhanced Sampling. Briefings in Bioinformatics 2024, 25, bbad429, DOI: 10.1093/bib/bbad429.

(29) Zhu, J.; von Bülow, S.; Liu, H.; Lindorff-Larsen, K.; Chen, H.-F. Extending Conformational Ensemble Prediction to Multidomain Proteins and Protein Complex. bioRxiv, 2026; DOI: 10.64898/2026.01.14.699584.

(30) DeCastro, S.; Zhang, O.; Liu, Z. H.; Forman-Kay, J. D.; Head-Gordon, T. IDPForge: Deep Learning of Proteins with Global and Local Regions of Disorder. bioRxiv, 2026; DOI: 10.64898/2026.03.25.714313.

(31) Novak, B.; Lotthammer, J. M.; Emenecker, R. J.; Holehouse, A. S. Accurate Predictions of Disordered Protein Ensembles with STARLING. Nature 2026, 652, 240–250, DOI: 10.1038/s41586-026-10141-2.

(32) Karras, T.; Aittala, M.; Aila, T.; Laine, S. Elucidating the Design Space of Diffusion-Based Generative Models. Advances in Neural Information Processing Systems. 2022; Vol. 35; pp 26565–26577, DOI: 10.52202/068431-1926.

(33) Meng, E. C.; Goddard, T. D.; Pettersen, E. F.; Couch, G. S.; Pearson, Z. J.; Morris, J. H.; Ferrin, T. E. UCSF ChimeraX: Tools for Structure Building and Analysis. Protein Science 2023, 32, e4792, DOI: 10.1002/pro.4792.

(34) Sohl-Dickstein, J.; Weiss, E.; Maheswaranathan, N.; Ganguli, S. Deep Unsupervised Learning Using Nonequilibrium Thermodynamics. Proceedings of the 32nd International Conference on Machine Learning. 2015; Vol. 37; pp 2256–2265.

(35) Ho, J.; Jain, A.; Abbeel, P. Denoising Diffusion Probabilistic Models. Advances in Neural Information Processing Systems. 2020; Vol. 33; pp 6840–6851.

(36) Ng, S. C.; Güttler, T.; Görlich, D. Recapitulation of Selective Nuclear Import and Export with a Perfectly Repeated 12mer GLFG Peptide. Nature Communications 2021, 12, 4047, DOI: 10.1038/s41467-021-24292-5.

(37) Honda, S.; Akiba, T.; Kato, Y. S.; Sawada, Y.; Sekijima, M.; Ishimura, M.; Ooishi, A.; Watanabe, H.; Odahara, T.; Harata, K. Crystal Structure of a Ten-Amino Acid Protein. Journal of the American Chemical Society 2008, 130, 15327–15331, DOI: 10.1021/ja8030533.

(38) Rauscher, S.; Gapsys, V.; Gajda, M. J.; Zweckstetter, M.; de Groot, B. L.; Grubmüller, H. Structural Ensembles of Intrinsically Disordered Proteins Depend Strongly on Force Field: A Comparison to Experiment. Journal of Chemical Theory and Computation 2015, 11, 5513–5524, DOI: 10.1021/acs.jctc.5b00736.

(39) Shalongo, W.; Dugad, L.; Stellwagen, E. Distribution of Helicity within the Model Peptide Acetyl(AAQAA)3amide. Journal of the American Chemical Society 1994, 116, 8288–8293, DOI: 10.1021/ja00097a039.

(40) Huang, J.; Rauscher, S.; Nawrocki, G.; Ran, T.; Feig, M.; de Groot, B. L.; Grubmüller, H.; MacKerell, A. D. CHARMM36m: An Improved Force Field for Folded and Intrinsically Disordered Proteins. Nature Methods 2017, 14, 71–73, DOI: 10.1038/nmeth.4067.

(41) Piana, S.; Lindorff-Larsen, K.; Shaw, D. E. How Robust Are Protein Folding Simulations with Respect to Force Field Parameterization? Biophysical Journal 2011, 100, L47–L49, DOI: 10.1016/j.bpj.2011.03.051.

(42) McInnes, L.; Healy, J.; Saul, N.; Großberger, L. UMAP: Uniform Manifold Approximation and Projection. Journal of Open Source Software 2018, 3, 861, DOI: 10.21105/joss.00861.

(43) Maier, J. A.; Martinez, C.; Kasavajhala, K.; Wickstrom, L.; Hauser, K. E.; Simmerling, C. ff14SB: Improving the Accuracy of Protein Side Chain and Backbone Parameters from ff99SB. Journal of Chemical Theory and Computation 2015, 11, 3696–3713, DOI: 10.1021/acs.jctc.5b00255.

(44) Gervais, F.; Paquette, J.; Morissette, C.; Krzywkowski, P.; Yu, M.; Azzi, M.; Lacombe, D.; Kong, X.; Aman, A.; Laurin, J.; Szarek, W. A.; Tremblay, P. Targeting Soluble A*β* Peptide with Tramiprosate for the Treatment of Brain Amyloidosis. Neurobiology of Aging 2007, 28, 537–547, DOI: 10.1016/j.neurobiolaging.2006.02.015.

(45) Lazar, T.; Connor, A.; DeLisle, C. F.; Burger, V.; Tompa, P. Targeting Protein Disorder: The next Hurdle in Drug Discovery. Nature Reviews Drug Discovery 2025, 24, 743–763, DOI: 10.1038/s41573-025-01220-6.

(46) Tunyasuvunakool, K.; Adler, J.; Wu, Z.; Green, T.; Zielinski, M.; Žídek, A.; Bridgland, A.; Cowie, A.; Meyer, C.; Laydon, A.; Velankar, S.; Kleywegt, G. J.; Bateman, A.; Evans, R.; Pritzel, A.; Figurnov, M.; Ronneberger, O.; Bates, R.; Kohl, S. A. A.; Potapenko, A.; Ballard, A. J.; Romera-Paredes, B.; Nikolov, S.; Jain, R.; Clancy, E.; Reiman, D.; Petersen, S.; Senior, A. W.; Kavukcuoglu, K.; Birney, E.; Kohli, P.; Jumper, J.; Hassabis, D. Highly Accurate Protein Structure Prediction for the Human Proteome. Nature 2021, 596, 590–596, DOI: 10.1038/s41586-021-03828-1.

(47) Anderson, B. D. Reverse-Time Diffusion Equation Models. Stochastic Processes and their Applications 1982, 12, 313–326, DOI: 10.1016/0304-4149(82)90051-5.

(48) Särkkä, S.; Solin, A. Applied Stochastic Differential Equations, first edition. ed.; Institute of Mathematical Statistics Textbooks; Cambridge University Press, 2019; DOI: 10.1017/9781108186735.

(49) Xiang, S.; Gapsys, V.; Kim, H.-Y.; Bessonov, S.; Hsiao, H.-H.; Möhlmann, S.; Klaukien, V.; Ficner, R.; Becker, S.; Urlaub, H.; Lührmann, R.; de Groot, B.; Zweckstetter, M. Phosphorylation Drives a Dynamic Switch in Serine/Arginine-Rich Proteins. Structure 2013, 21, 2162–2174, DOI: 10.1016/j.str.2013.09.014.

(50) Lindorff-Larsen, K.; Piana, S.; Dror, R. O.; Shaw, D. E. How Fast-Folding Proteins Fold. Science 2011, 334, 517–520, DOI: 10.1126/science.1208351.

