## Supplementary Information for "Generating Structural Ensembles of Disordered Proteins with Diffusion Models"

### A Coarse-Graining

#### A.1 Defining the Coarse-grained Beads

Two coarse-grained (CG) beads are defined for each residue, one for the backbone and one for the side chain. The sizes of these beads were determined using the Top2018 dataset,[1] a curated set of crystal structures from the Protein Data Bank (PDB). The center of the backbone bead is located at the C$\alpha$ atom. The bead's radius is based on the maximum distance between the C$\alpha$ atom and any backbone atom, including hydrogens. The maximum of these distances across all curated structures was found to be $\sim 2.6$ Å. Consequently, the backbone bead's radius was set to 2.7 Å.

For side chains, a distinct CG bead with an associated center and radius is defined for each of the 20 conventional amino acid residues. The properties of each CG bead are defined through an optimization procedure that identifies the smallest bead capable of encapsulating the entire side chain for all geometries of that residue type observed in the Top2018 dataset. First, for a given residue in a given structure, a bead is centered on each heavy atom in the side chain. The radius of each bead is defined as the maximum distance between its center and any side-chain atom, including hydrogens. To account for the variability of side-chain geometries, this process is iterated over all structures in the Top2018 dataset, and the largest radius found for each choice of center atom in each residue type is used to define a candidate CG bead. Finally, considering all candidate CG beads for a given residue type, the bead with the smallest radius defines the side-chain CG bead for that residue type.

The CG bead definition process for the side chains corresponds to a min-max optimization over all possible bead centers and radii. Specifically, let $\mathbf{x}_i^{(J)}$ correspond to the position of atom $i$ in the $J^{\text{th}}$ sample of the curated dataset of crystal structures. For simplicity, we assume that all samples have a single topology for each residue type, i.e. each residue has a single protonation state. Let $I_\alpha$ denote the set of all side-chain atom indices in the molecular graph of residue $\alpha$, where $\alpha \in \{1, \ldots, 20\}$ indexes all possible residue types, and let $H$ denote the index set of all heavy atoms. Then, the atom index of the CG bead's center $i_\alpha$ and its radius $\tilde{r}_\alpha$ are determined as follows:

$$\begin{aligned} i_\alpha &= \underset{i \in I_\alpha \cap H}{\arg\min} \max_{J,\ j \in I_\alpha} \left\| \mathbf{x}_i^{(J)} - \mathbf{x}_j^{(J)} \right\| \\ \tilde{r}_\alpha &= \min_{i \in I_\alpha \cap H} \max_{J,\ j \in I_\alpha} \left\| \mathbf{x}_i^{(J)} - \mathbf{x}_j^{(J)} \right\| \end{aligned} \tag{1}$$

For the special case of glycine, which has no heavy atoms in its side chain, this process is modified. The side-chain CG bead of glycine is located on the backbone carbonyl $C$ atom with a radius of 0.0 Å. This choice is arbitrary, but it simplifies the data structure by ensuring that all coarse-grained protein structures have $2N$ nodes, where $N$ denotes the number of residues. During training and sampling, the coarse-structure model learns to position the CG side-chain bead for each glycine residue. However, when a given set of predicted CG bead positions is used to sample heavy-atom positions, the positions of glycine side-chain CG beads are discarded.

Finally, the radius of each side-chain CG bead is ceiled to the next highest multiple of 0.1 Å to avoid floating-point errors:

$$r_\alpha = \left\lceil \frac{\tilde{r}_\alpha}{a} \right\rceil a, \quad a = 0.1\,\text{Å} \tag{2}$$

For each residue, the atom at which the side-chain CG bead's center is located and the bead's radius are given in Supplementary Table 1.

#### A.2 Complexity Analysis of Graph Convolutions

In the CG model, graph convolutions are performed for all possible pairs of CG beads, implying a complexity of $O((2N)^2)$, where $N$ denotes the number of residues in the sequence. In the fine-structure model, graph convolutions are performed for each pair of heavy atoms whose CG beads are in contact. Two CG beads located at positions $\mathbf{x}_1, \mathbf{x}_2$ with radii $r_1, r_2$

Supplementary Table 1: **Definition of the Coarse-grained Beads**
For each of the 20 commonly occurring amino acid residues, the atom at which the side-chain CG bead's center is located and the bead's radius are provided. The atom names follow the PDB nomenclature.

| Residue | Center Atom (PDB) | Radius (Å) |
|---|---|---|
| ALA | CB | 1.1 |
| ARG | CD | 5.1 |
| ASN | CG | 2.3 |
| ASP | CG | 2.3 |
| CYS | CB | 2.5 |
| GLN | CG | 3.6 |
| GLU | CG | 3.4 |
| GLY | C | 0.0 |
| HIS | CG | 3.4 |
| ILE | CB | 3.7 |
| LEU | CG | 2.8 |
| LYS | CD | 3.8 |
| MET | SD | 4.0 |
| PHE | CD1 | 4.0 |
| PRO | CG | 2.3 |
| SER | CB | 2.2 |
| THR | CB | 2.3 |
| TRP | CD2 | 4.0 |
| TYR | CD2 | 4.2 |
| VAL | CB | 2.3 |

are considered in contact when the following condition is satisfied:

$$\|\mathbf{x}_1 - \mathbf{x}_2\| < r_1 + r_2 + b \tag{3}$$

where $b$ is a buffer distance set to 2.0 Å to capture some of the dispersion interactions between the atoms in these residues.

Let $n_i$ denote the number of heavy atoms approximated by the $i^{\text{th}}$ CG bead of a given protein structure. For backbone CG beads, $n_i = 4$, while for side-chain CG beads, $0 \leq n_i \leq 10$. For a given protein structure with CG adjacency matrix $A$, the computational complexity of one layer of the heavy-atom graph neural network is $O(\sum_{i,j,A_{ij}=1} n_i n_j)$, which strongly depends on the spatial sparsity of the protein structure.

To simplify the complexity analysis below, let us separate the number of heavy atoms in the backbone (four per residue) and side chain ($n_{s,i}$), where $i$ now indexes the residues in the protein sequence. Furthermore, let $\langle n_{s,i}\rangle$ denote the average number of side-chain heavy atoms per residue. Because complexity depends on spatial sparsity, we consider two cases:

- For sparse (expanded) protein structures, only CG beads that are adjacent along the protein sequence interact with each other. In this case, the graph convolution complexity can be as low as $O(\sum_i (4+n_{s,i})^2 + (4+n_{s,i})(4+n_{s,i+1}) + (4+n_{s,i})(4+n_{s,i-1})) = O(3(4+\langle n_{s,i}\rangle)^2 N)$.

- For dense (compact) protein structures, the complexity is larger than the sparse case, but remains much smaller than $O(N^2)$. The number of interacting CG beads depends on the local packing density. The packing fraction of protein hydrophobic cores is 0.56,[2] and the number of neighboring residues in compact structures is usually less than twelve.[3] The complexity of graph convolutions can be as large as $O(12(4+\langle n_{s,i}\rangle)^2 N)$ if we assume each residue has twelve neighbors. Even with this conservative estimate, the graph convolution complexity remains $O(N)$, but with a larger prefactor compared to the case of sparse structures.

### B Ensemble Generation Strategy

#### B.1 Relating the Diffusion Model SDE to the Noise Schedule

The diffusion process can be formalized through the following stochastic differential equation (SDE):

$$d\mathbf{x} = h(t)\mathbf{x}dt + g(t)d\mathbf{w}_t \tag{4}$$

where $\mathbf{w}_t$ is a standard Wiener process and $h(t), g(t)$ are predetermined functions of time $t \in [t_{\min}, t_{\max}]$. The probability density function $p(\mathbf{x}, t)$ of obtaining a given value $\mathbf{x}$ at time

$t$ is given by:

$$p(\mathbf{x}, t) = \mathcal{N}(\mathbf{x}; \alpha(t)\mathbf{x}_0, \sigma^2(t)I) \tag{5}$$

$$\alpha(t) = \exp\left(\int_0^t h(s)ds\right), \quad \sigma^2(t) = \alpha(t)^2 \int_0^t \frac{g(s)^2}{\alpha(s)^2} ds \tag{6}$$

where $\mathcal{N}(\mathbf{x}; \mu, \sigma^2)$ denotes a normal distribution with mean $\mu$ and variance $\sigma^2$, $\mathbf{x}_0$ denotes the initial conformation and the functions $(\alpha(t), \sigma(t))$ correspond to the noise schedule. Inverting eq. (6) allows one to relate the SDE functions $(g(t), h(t))$ to the noise schedule:

$$h(t) = \frac{\dot{\alpha}}{\alpha}, \quad g(t) = \sqrt{2\sigma^2\left(\frac{\dot{\sigma}}{\sigma} - \frac{\dot{\alpha}}{\alpha}\right)} \tag{7}$$

#### B.2 Training

The denoising models $D_\theta(\mathbf{x}, t)$ are trained by minimizing the conventional diffusion loss function:

$$\mathcal{L}_{\text{diff}} = \mathop{\mathbb{E}}_{\substack{\mathbf{x}_0 \sim p_{\text{data}},\, t \sim p_n \\ \mathbf{x} \sim \mathcal{N}(\alpha(t)\mathbf{x}_0, \sigma^2(t)\mathbf{I})}} \left[\lambda(t) \left\| D_\theta(\mathbf{x}, t) - \mathbf{x}_0 \right\|^2\right] \tag{8}$$

where $p_{\text{data}}$ denotes the training data distribution, $p_n$ denotes the denoising time distribution, and $\lambda(t)$ is a time-dependent function used to reweight the importance of each time across the temporal range. In practice, $p_n$ is set to a log-uniform distribution in the range $[t_{\min}, t_{\max}]$, whose boundaries are treated as hyperparameters.

To stabilize training and focus learning on the most relevant denoising times, Karras et al.[4] suggest preconditioning the denoiser's input and output in the following manner:

$$D_\theta(\mathbf{x}, t) = c_{\text{skip}}(t)\mathbf{x} + c_{\text{out}}(t) F_\theta(c_{\text{in}}(t)\mathbf{x}, t) \tag{9}$$

$$c_{\text{skip}}(t) = \frac{\sigma_{\text{data}}^2}{\sigma_{\text{data}}^2 + \sigma^2(t)}, \quad c_{\text{out}}(t) = \frac{\sigma_{\text{data}}\sigma(t)}{\sqrt{\sigma_{\text{data}}^2 + \sigma^2(t)}}, \quad c_{\text{in}}(t) = \frac{1}{\sqrt{\sigma_{\text{data}}^2 + \sigma^2(t)}} \tag{10}$$

where $F_\theta(\mathbf{y}, t)$ is a neural network parameterized by $\theta$ and Karras' prescribed loss function weight is:

$$\lambda(t) = \frac{1}{c_{\text{out}}(t)^2} \tag{11}$$

In this work, the value of $\sigma_{\text{data}}$ depends on the training dataset structures and its definition differs for the coarse-structure model versus the fine structure model. For the coarse-structure model, $\sigma_{\text{data}}$ denotes the root mean square deviation (RMSD) of the atom positions about their centroid, averaged across all three dimensions:

$$\sigma_{\text{data,CG}} = \sqrt{\frac{1}{3|J_{\text{train}}|} \sum_{j \in J_{\text{train}}} \sum_{i=1}^{n} \|\mathbf{x}_i^j - \bar{\mathbf{x}}^j\|^2} \tag{12}$$

For the fine-structure model, $\sigma_{\text{data}}$ denotes the RMSD of atomic coordinates centered onto their coarse-grained bead representative:

$$\sigma_{\text{data,HA}} = \sqrt{\frac{1}{3|J_{\text{train}}|} \sum_{j \in J_{\text{train}}} \sum_{i=1}^{n} \|\mathbf{x}_i^j - \mathbf{c}_{\mathcal{C}(i)}^j\|^2} \tag{13}$$

where $\mathcal{C}(i)$ denotes the CG bead index of atom $i$.

The functional form of the $c_{\text{skip}}$, $c_{\text{out}}$, $c_{\text{in}}$ functions serves multiple purposes. First, $c_{\text{skip}}(t)$ monotonically interpolates between the input $\mathbf{x}$ and the neural network's output since $\sigma(t)$ is monotonic. For $t \sim 0$, the network's input $\mathbf{x}$ is a close approximation of the denoised coordinates $\mathbf{x}_0$. $c_{\text{skip}}(t)$ incorporates this prior knowledge to facilitate the learning of $F_\theta(\mathbf{y}, t)$. For $t \sim 0$, $c_{\text{skip}}(t) \sim 1$ and therefore only small differences $(\mathbf{x}_0 - c_{\text{skip}}(t)\mathbf{x})$ need to be learned by $F_\theta(\mathbf{y}, t)$. For large noise levels ($t \gg 1$), the input is dominated by noise and is therefore not a reliable approximation of $\mathbf{x}_0$. In this regime, $c_{\text{skip}}(t) \sim 0$, which implies that the denoiser's output is entirely determined by the network's output. In addition, $c_{\text{in}}(t)$ rescales the input of the network to ensure that it has unit variance, enabling faster training. Finally, $c_{\text{out}}(t)$, together with the specific form of $c_{\text{skip}}(t)$, ensures that the effective target of the neural network under the squared error loss (see eq. (8)) has a minimal variance, which minimizes noise in the parameter gradients during learning. We refer the reader to Karras et al.[4] for a detailed discussion of these parametric choices.

Motivated by the results of Karras et al.,[4] we choose a linear-in-time noise schedule, which induces the following associated parametric functions:

$$\alpha(t) = 1, \quad \sigma(t) = t, \quad h(t) = 0, \quad g(t) = \sqrt{2t} \tag{14}$$

This noise schedule is used to train all models. The ranges of the noise schedule for the coarse-structure and fine-structure models differ. For the coarse-structure model, $[t^{\text{CG}}_{\text{min}}, t^{\text{CG}}_{\text{max}}] = [0.1\,\text{Å}, 50\,\text{Å}]$ is used, while for the fine-structure model, $[t^{\text{HA}}_{\text{min}}, t^{\text{HA}}_{\text{max}}] = [0.002\,\text{Å}, 5\,\text{Å}]$ is used. Moreover, during sampling, the range of the noise schedule is discretized in logarithmic space:

$$t_i = e^{a\tau_i + b}, \quad \tau_i = \frac{i}{N}, \quad i \in \mathbb{Z}, 0 \leq i \leq N, \quad a = \log\left(\frac{\sigma_{\text{min}}}{\sigma_{\text{max}}}\right), \quad b = \log(\sigma_{\text{max}}) \tag{15}$$

where $N$ is the total number of temporal intervals. During training, a uniform distribution for $\tau$, i.e. $p(\tau) = \mathcal{U}(0, 1)$, is used to sample the noising times of each batch element. This ensures that all noising times are sampled with equal probability in logarithmic space. Using the change of variable formula, this induces the following $p_n(t)$ distribution:

$$p(\tau) = \mathcal{U}(0, 1) \to p_n(t) = \frac{1}{|a|t} \tag{16}$$

The noise schedule profiles for the coarse-structure and fine-structure models are given as a function of $\tau$ in Supplementary Fig. 1.

Models are trained to minimize the denoising loss function eq. (8) using stochastic gradient-based optimization. The Adam optimizer[5] is used with parameters $\beta_1 = \beta_2 = 0.9$ and a batch size of 64 structures. A parameter update is performed using gradients calcu-

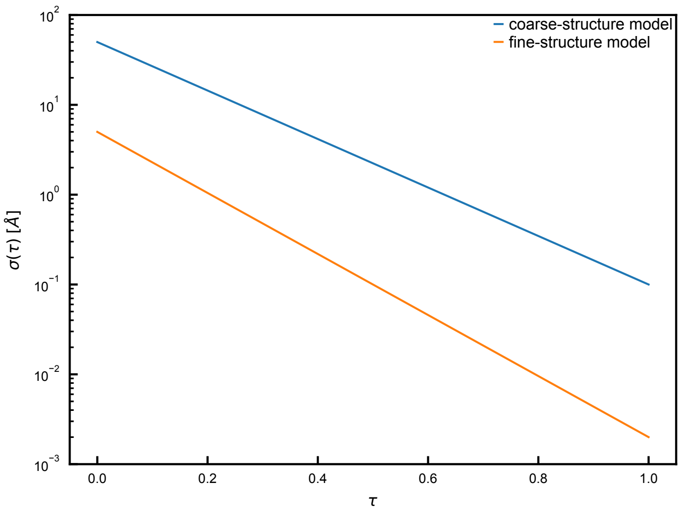

Supplementary Figure 1: **Noise Schedule Profiles**
The value of the noise standard deviation of the noise schedule $\sigma(t)$ is shown as a function of the parametric time $\tau$ for both the coarse-structure and fine-structure models.

lated on one batch of structures. Batches are formed by sampling structures randomly from the training dataset without replacement. Once all structures have been sampled, they are placed back in the pool of samples, and this cycle repeats until the end of the training. All models are trained for 4000 epochs, where one epoch corresponds to 32 parameter updates. The learning rate is determined by a piecewise function of the epoch number, consisting of a linear warm-up period, a plateau and an exponential decay phase (Supplementary Fig. 2). Note that the learning rate schedules of the coarse-structure and fine-structure models differ.

To ensure a smooth progression of the model parameters, an exponential moving average of the parameters is maintained throughout training and used to evaluate validation metrics and produce final samples. The exponential moving average is updated after every optimizer step $j$ as follows:

$$\bar{w}_i^{j+1} = \beta \bar{w}_i^j + (1 - \beta) w_i^j \tag{17}$$

where $w_i, \bar{w}_i$ denote the $i^{\text{th}}$ model parameter and its average, respectively. For all training runs, a decay rate of $\beta = 0.999$ is used.

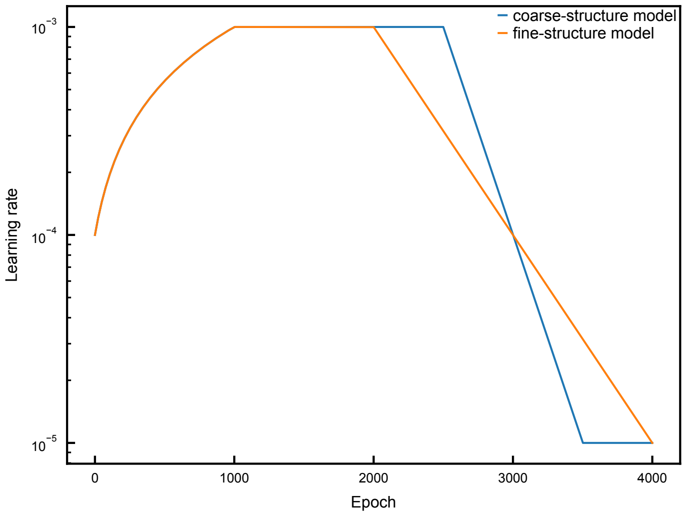

Supplementary Figure 2: **Learning Rate Schedules**
The learning rate schedules for both the coarse-structure and fine-structure models are shown.

#### B.3 Monitored Training Metrics

In addition to monitoring the diffusion loss on the validation split of the dataset, the model's accuracy is assessed every 50 epochs using several statistics of the structural and energy metrics (see Supplementary Appendix E for definitions of metrics). These statistics are calculated on an ensemble of 100 structures generated with 100 denoising steps. When training the fine-structure model, a fixed subset of the validation dataset is used to define the required input CG bead positions.

For a given metric $x$, the errors in the average ($\langle x \rangle$) and standard deviation ($\mathrm{std}(x)$) are calculated relative to the training and validation datasets:

$$\Delta_{\mathrm{train}}\langle x \rangle = \frac{\langle x \rangle_{\mathrm{sample}}}{\langle x \rangle_{\mathrm{train}}} - 1, \qquad \Delta_{\mathrm{valid}}\langle x \rangle = \frac{\langle x \rangle_{\mathrm{sample}}}{\langle x \rangle_{\mathrm{valid}}} - 1 \tag{18}$$

$$\Delta_{\mathrm{train}}\mathrm{std}(x) = \frac{\mathrm{std}(x)_{\mathrm{sample}}}{\mathrm{std}(x)_{\mathrm{train}}} - 1, \qquad \Delta_{\mathrm{valid}}\mathrm{std}(x) = \frac{\mathrm{std}(x)_{\mathrm{sample}}}{\mathrm{std}(x)_{\mathrm{valid}}} - 1 \tag{19}$$

The tracked metrics are a subset of those used to assess the generated samples (Supplementary Fig. 3, Supplementary Fig. 4). In particular, during training of the fine-structure model, energy-based metrics still improve after the diffusion loss saturates to its final value (Supplementary Fig. 4). Therefore, saturation of the energy-based metrics is used to determine a suitable number of training epochs. Note that the relative errors of most energy-based metrics reach negative values since a 100-sample subset is insufficient to probe the higher moments of the training and validation dataset distributions. Finally, changes in the tracked metrics are used to optimize the model's architecture and training strategy.

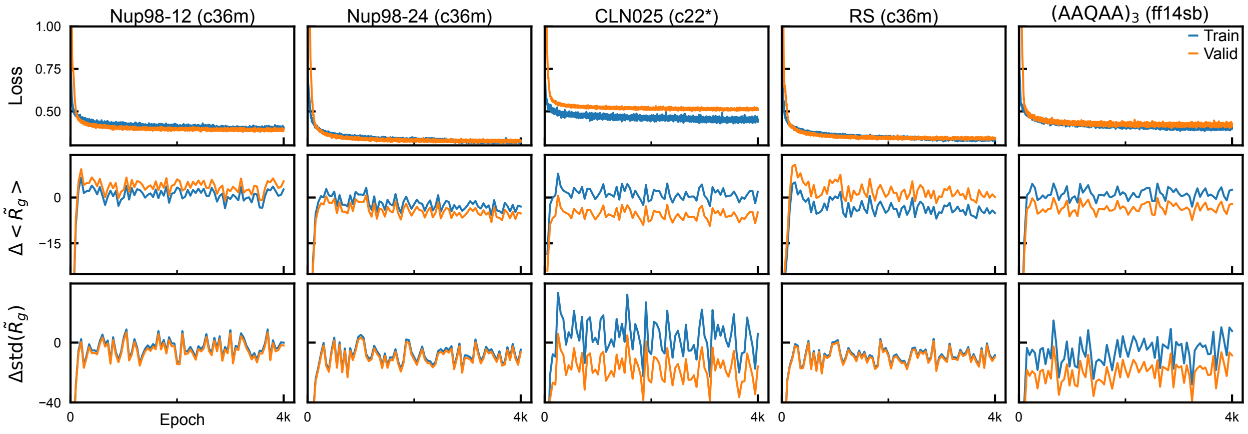

Supplementary Figure 3: **Training Metrics of the Coarse-Structure Model**
Tracked metrics of coarse-structure model training are shown relative to the training and validation datasets of all studied systems. $\tilde{R}_g$ denotes the radius of gyration with unit masses, which weighs all positions equally.

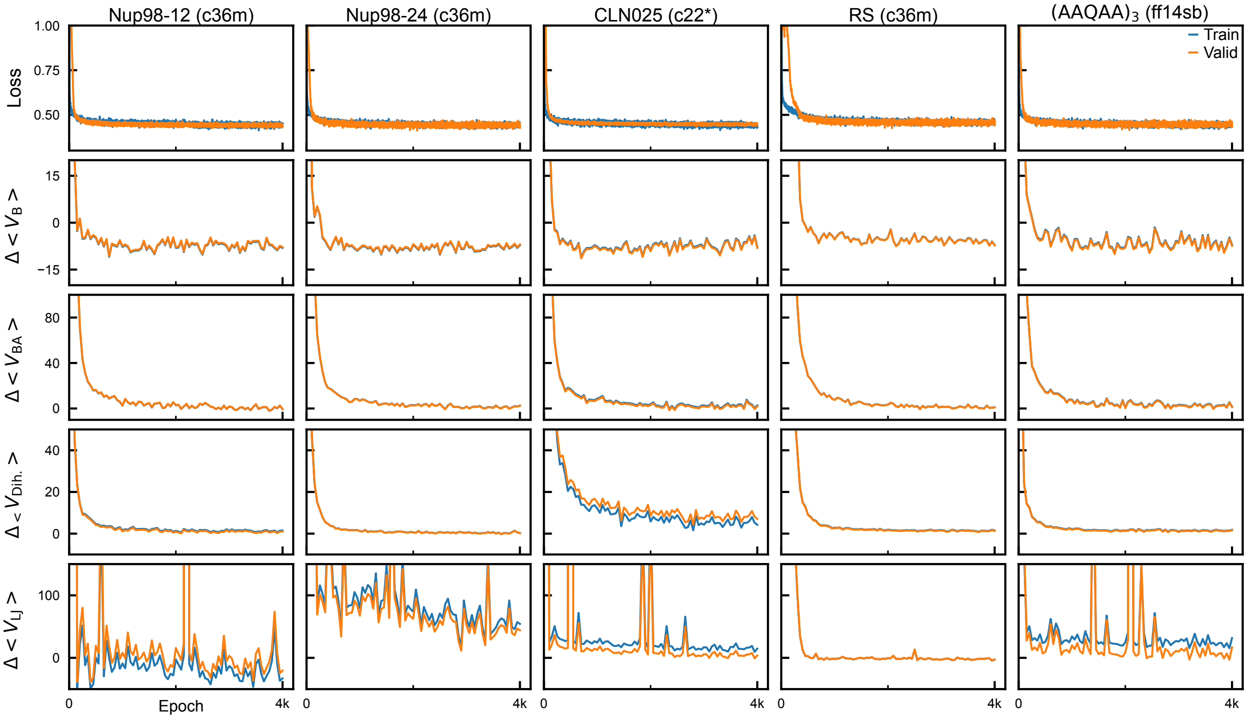

Supplementary Figure 4: **Training Metrics of the Fine-Structure Model**
Tracked metrics (rows) of fine-structure model training are shown relative to the training and validation datasets for all tested systems (columns). The bond energy ($V_{\mathrm{B}}$), bond angle energy ($V_{\mathrm{BA}}$), dihedral angle energy ($V_{\mathrm{Dih.}}$) and Lennard-Jones energy ($V_{\mathrm{LJ}}$) relative errors are shown. Definitions of the energy metrics are given in Supplementary Appendix E.2.

#### B.4 Sampling

To sample structures from the generative model, the reverse diffusion process is numerically approximated using the Euler-Maruyama integrator,[6] where the time $t$ is log-linearly discretized (eq. (15)) over the range of the denoising time sampling distribution used during training, i.e. $[\sigma_{\min}, \sigma_{\max}]$:

$$\begin{aligned}
\mathbf{x}^{(i+1)} &= \mathbf{x}^{(i)} + \left(h(t_i)\mathbf{x}^{(i)} - g(t_i)^2 \nabla_{\mathbf{x}} \log(\hat{p}(\mathbf{x}^{(i)}, t_i))\right) \Delta t_i + g(t_i)\sqrt{|\Delta t_i|}\epsilon_i, \qquad 0 \leq i < N \\
&= \mathbf{x}^{(i)} - 2t_i \Delta t_i \nabla_{\mathbf{x}} \log(\hat{p}(\mathbf{x}^{(i)}, t_i)) + \sqrt{2t_i|\Delta t_i|}\epsilon_i \qquad (20)
\end{aligned}$$

$$\Delta t_i = t_{i+1} - t_i < 0, \quad \epsilon_i \sim \mathcal{N}(0, 1) \qquad (21)$$

where $i$ increases from 0 to $N-1$ during sampling as it corresponds to the reverse direction of the diffusion process (Supplementary Fig. 1).

The initial node positions of the coarse and fine structures are sampled differently. For coarse structures, initial conformations are determined by sampling from an approximation of the position probability density (eq. (5)) at the maximum noise level $\sigma(t_{\max}^{\mathrm{CG}})$:

$$\mathbf{x}_{\mathrm{CG},i}^{(0)} \sim \widetilde{p}(\mathbf{x}, t_{\max}^{\mathrm{CG}}) = \mathcal{N}(\mathbf{x}; 0, \sigma^2(t_{\max}^{\mathrm{CG}})I) \qquad (22)$$

where the same distribution is used for all CG beads $i$ in the given protein structure. For fine structures, heavy-atom positions are initialized such that the expected centroid of the atoms approximated by each CG bead matches the given CG bead position $\mathbf{c}_j$:

$$\mathbf{x}_{\mathrm{HA},i}^{(0)} \sim \widetilde{p}(\mathbf{x}, t_{\max}^{\mathrm{HA}}) = \mathcal{N}(\mathbf{x}; \mathbf{c}_{\mathcal{C}(i)}, \sigma^2(t_{\max}^{\mathrm{HA}})I) \qquad (23)$$

where $\mathcal{C}(i)$ denotes the CG bead index of atom $i$.

Following the Chroma model,[7] the sampling SDE is modified with additional equilibration to efficiently sample the marginal distributions $\hat{p}(\mathbf{x}, t_i)$. This is implemented by rescaling the magnitude of the Langevin steps, which is valid provided that the time intervals are small, i.e. $|\Delta t_i| \ll 1$:

$$d\mathbf{x} = \left[h(t)\mathbf{x} - \left(1 + \frac{\psi}{2}\right) g(t)^2 \nabla_{\mathbf{x}} \log(\hat{p}(\mathbf{x}, t))\right] dt + \sqrt{1+\psi} g(t) d\mathbf{w}_t \tag{24}$$

where $\psi$ is a parameter that controls the rate of equilibration, with $\psi = 0$ corresponding to no equilibration. As seen in eq. (24), $\psi$ effectively increases the magnitude of the score function and the Wiener process terms. The increased magnitude of the Wiener process step enhances the diversity of the sampled paths by preventing the sampling process from falling into local minima of the marginal distributions $\hat{p}(\mathbf{x}, t)$. The increased magnitude of the score function is necessary to counteract the increase in the added noise, ensuring convergence of the sampling process.

When sampling CG structures, a value of $\psi = 10$ is used, while $\psi = 2$ is used to sample heavy-atom positions. These values are suitable for the goals of the present work, but would likely require fine-tuning for other purposes. The effect of $\psi$ on the force field energies is assessed for the Nup98-12 system for $\psi \in \{0, 1, 2, 5, 10\}$ (Supplementary Figs. 5 and 6). When assessing the sampling of heavy-atom structures, we find that $\psi$ most significantly affects the bond energy distribution and a value of $\psi \geq 2$ provides reasonably-equilibrated distributions (Supplementary Fig. 5). To minimize discretization errors in the sampling SDE, the lowest value of $\psi$ ($\psi = 2$) that achieves an accurate bond energy distribution is used. When assessing the sampling of CG structures, a value of $\psi = 10$ provides the best agreement across the bond length, bond angle and dihedral angle energies (Supplementary Fig. 6). $\psi = 10$ is, therefore, used to sample all CG structures.

Finally, each generated dataset contains the same number of samples as its corresponding training dataset. This ensures that the discrepancies observed between the generated and training datasets are not caused by differences in the dataset size. The only exception is the CLN025 experiments with different RMSD thresholds, for which the number of generated structures is fixed at 25,000 (see Supplementary Appendix F.5). The dataset size of all generated datasets is summarized in Supplementary Table 8.

#### B.5 Conditioners

When sampling heavy-atom structures, a set of conditions $C$ on the geometries is used to guide the denoising trajectories towards physically relevant landscapes. These conditions are applied using the conditioner framework of the Chroma model.[7] These conditioners essentially modify the learned score function with additive terms. This is a consequence of Bayes' theorem since the score of the conditional marginal distributions $\hat{p}(\mathbf{x}, t|C)$ is expressed as the sum of the unconditional score and a conditioning gradient:

$$\nabla_{\mathbf{x}} \log(\hat{p}(\mathbf{x}, t|C)) = \nabla_{\mathbf{x}} \log(\hat{p}(\mathbf{x}, t)) + \nabla_{\mathbf{x}} \log(p(C|\mathbf{x}, t)) \tag{25}$$

In our approach, all sampling conditioners restrain the range of a predetermined set of pairwise distances in the structure, i.e. $\{l_{ij} < \|\mathbf{x}_i^{(0)} - \mathbf{x}_j^{(0)}\| < u_{ij} | (i,j) \in I\}$. To derive expressions for the conditioning gradients implied by these conditioners, an expression for the posterior distribution $p(\mathbf{x}^{(0)}|\mathbf{x}, t)$ is needed. The posterior distribution is derived in Ingraham et al.[7] for the variance-preserving case with a noising kernel that has a non-trivial covariance matrix. We take similar steps to derive $p(\mathbf{x}^{(0)}|\mathbf{x}, t)$ in the variance-exploding case with a uniform noising covariance matrix. First, we assume a normal prior distribution for $\mathbf{x}^{(0)}$:

$$p(\mathbf{x}^{(0)}) = \mathcal{N}(\mathbf{x}^{(0)}; \mu_{\mathbf{0}}, \sigma_0^2 I) \tag{26}$$

where $\mu_{\mathbf{0}}$ and $\sigma_0$ control the location and size of the prior distribution. In practice, $\mu_{\mathbf{0}} = \mathbf{0}$ is used and $\sigma_0$ is estimated with the standard deviation of the nodes' positions in the structures of the training dataset. Using this prior and the uniform noising kernel, the

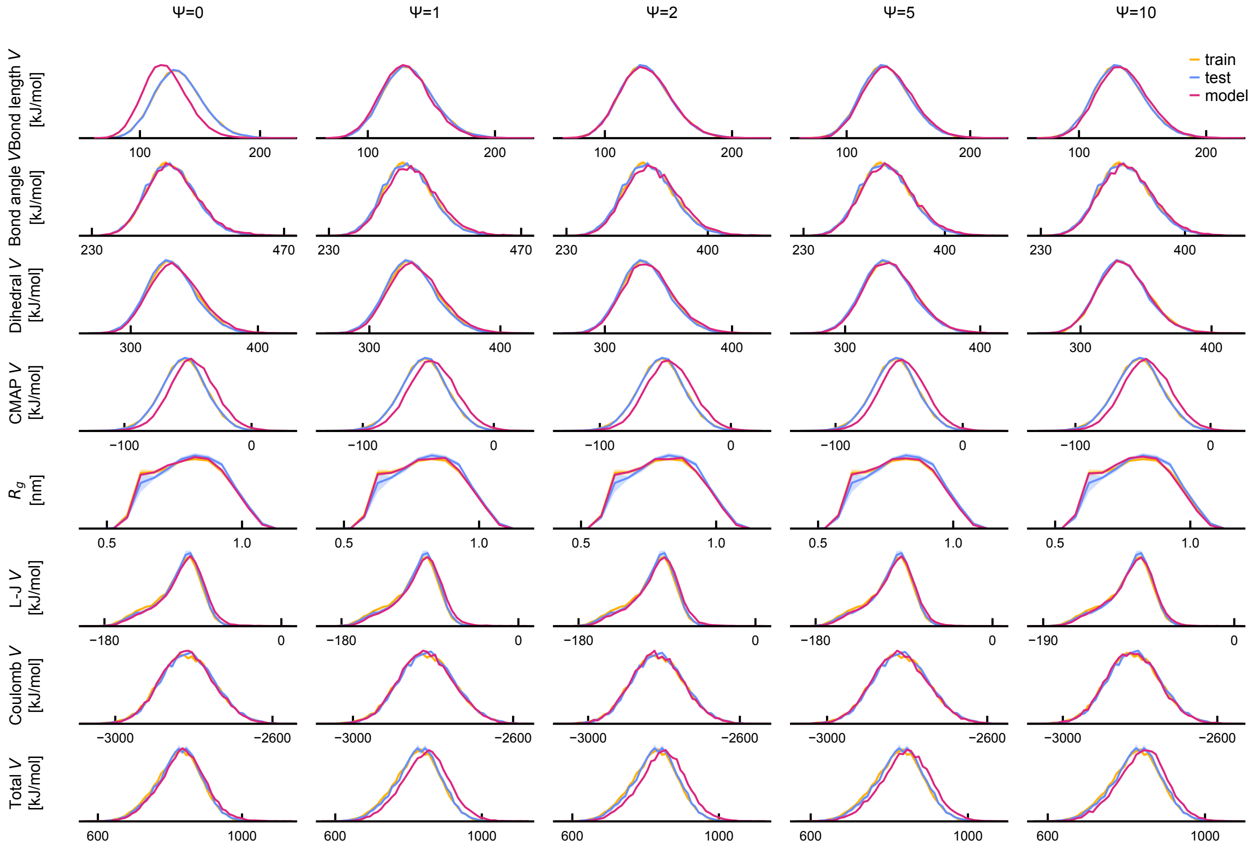

Supplementary Figure 5: **Effect of equilibration rate $\psi$ on fine-structure sampling** Probability densities of several fine- and coarse-structure metrics (rows) are shown for various amounts of equilibration ($\psi$) used during fine-structure sampling (columns). Supplementary Appendix E details how these metrics were computed. The distributions for the model, training and test sets are shown in each case. Distributions for the training and test sets are averaged over independent MD trajectories, and the standard error of the mean is shown (shaded area). The CG bead positions used to initialize the heavy-atom structures were taken from the training dataset.

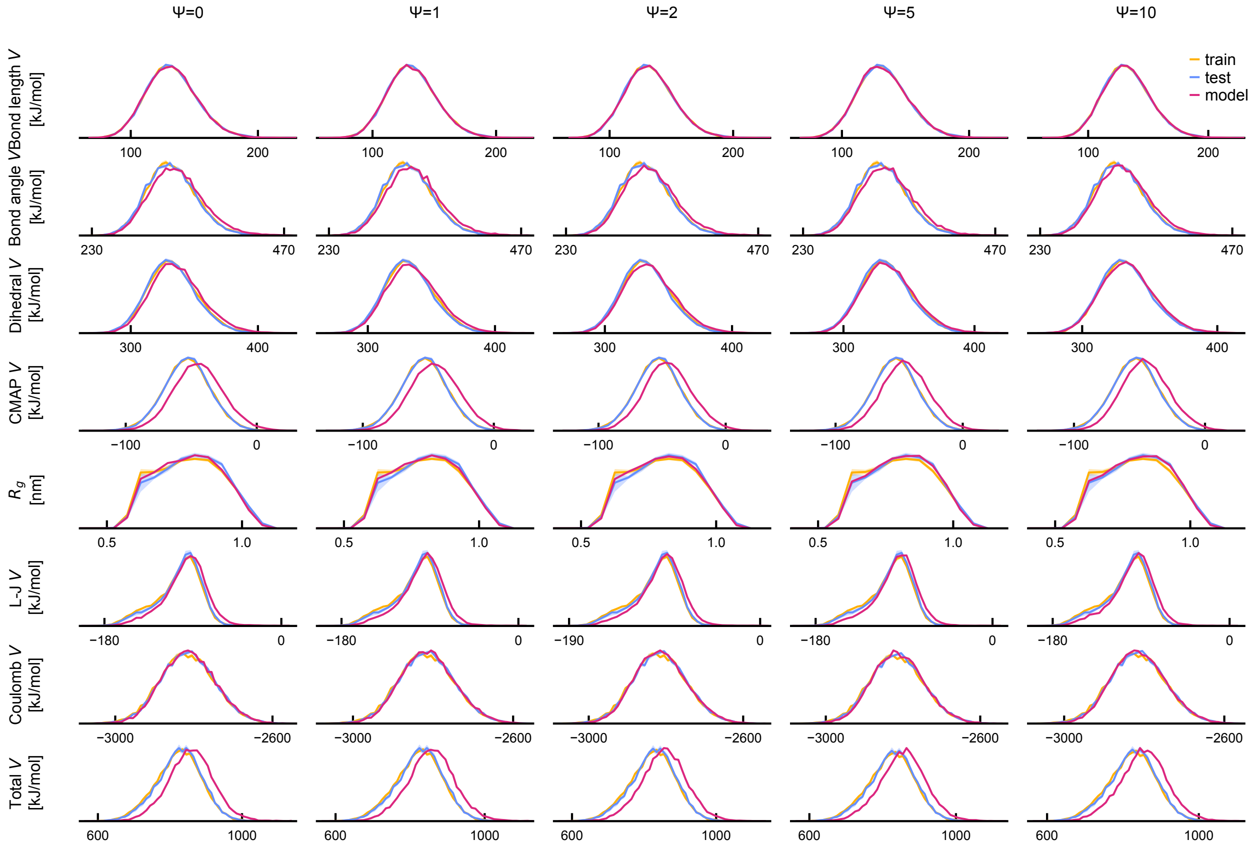

Supplementary Figure 6: **Effect of equilibration rate $\psi$ on coarse-structure sampling** Probability densities of several fine- and coarse-structure metrics (rows) are shown for various amounts of equilibration ($\psi$) used during coarse-structure sampling (columns). Supplementary Appendix E details how these metrics were computed. The distributions for the model, training and test sets are shown in each case. Distributions for the training and test sets are averaged over independent MD trajectories, and the standard error of the mean is shown (shaded area). A value of $\psi = 2$ was used to sample the heavy-atom positions.

posterior distribution of $\mathbf{x}^{(0)}$ is derived as follows:

$$
\begin{aligned}
p(\mathbf{x}^{(0)}|\mathbf{x}, t) &\propto p(\mathbf{x}, t|\mathbf{x}^{(0)})p(\mathbf{x}^{(0)}) && (27)\\
&\propto \mathcal{N}(\mathbf{x}; \mathbf{x}^{(0)}, \sigma^2(t)I)\,\mathcal{N}(\mathbf{x}^{(0)}; \mathbf{0}, \sigma_0^2 I) && (28)\\
&\propto \mathcal{N}(\mathbf{x}^{(0)}; \mathbf{x}, \sigma^2(t)I)\,\mathcal{N}(\mathbf{x}^{(0)}; \mathbf{0}, \sigma_0^2 I) && (29)\\
&= \mathcal{N}\left(\mathbf{x}^{(0)}; a_t\mathbf{x}, a_t\sigma^2(t)I\right), \quad \text{where} \quad a_t = \frac{\sigma_0^2}{\sigma_0^2 + \sigma^2(t)} && (30)
\end{aligned}
$$

where the last step follows from the well-known expression for combining a product of multivariate normal distributions:[8]

$$
\begin{aligned}
\mathcal{N}(\mathbf{x}; \mu_1, \Sigma_1)\mathcal{N}(\mathbf{x}; \mu_2, \Sigma_2) &\propto \mathcal{N}(\mathbf{x}; \mu_3, \Sigma_3) && (31)\\
\Sigma_3 &= (\Sigma_1^{-1} + \Sigma_2^{-1})^{-1} && (32)\\
\mu_3 &= \Sigma_3(\Sigma_1^{-1}\mu_1 + \Sigma_2^{-1}\mu_2) && (33)
\end{aligned}
$$

In practice, we use a non-informative prior with $\sigma_0 = \frac{1}{\sqrt{3}}$nm, which implies that structures generated from the prior's distribution have an expected unweighted radius of gyration of approximately 1 nm. This choice of $\sigma_0$ implies that $a_t$ rapidly converges to 1 when sampling the fine structure. Indeed, as seen in the fine-structure noise schedule (Supplementary Fig. 1), $\sigma(t) \approx 0.1$ nm at $\tau \approx 0.2$, which implies that $a_t \approx 0.97$ at this point and increases toward 1 as sampling progresses to smaller length scales ($\tau \to 1$).

To restrain pairwise distances of denoised positions $\mathbf{v}_{ij}^{(0)} = \mathbf{x}_i^{(0)} - \mathbf{x}_j^{(0)}$, eq. (30) is used to derive the posterior distribution of $\mathbf{v}_{ij}^{(0)}$:

$$
p(\mathbf{v}_{ij}^{(0)}|\mathbf{x}, t) = \mathcal{N}\left(\mathbf{v}_{ij}^{(0)}; a_t(\mathbf{x}_i - \mathbf{x}_j), 2a_t\sigma^2(t)I\right) \qquad (34)
$$

This implies that $\frac{\|\mathbf{v}_{ij}^{(0)}\|}{\sqrt{2a_t}\sigma(t)}$ has a non-central chi distribution. We use this property to enforce the distance $\|\mathbf{x}_i^{(0)} - \mathbf{x}_j^{(0)}\|$ to be within a fixed range $[l_{ij}, u_{ij}]$:

$$\log(p(\{l_{ij} < \|\mathbf{x}_i^{(0)} - \mathbf{x}_j^{(0)}\| < u_{ij}\}|\mathbf{x}, t)) = \log\left(\mathrm{CDF}_{\chi'}\left(\frac{u_{ij}}{\sqrt{2a_t}\sigma(t)}, \frac{\sqrt{a_t}\|\mathbf{x}_i - \mathbf{x}_j\|}{\sqrt{2}\sigma(t)}, 3\right)\right.$$
$$\left. - \mathrm{CDF}_{\chi'}\left(\frac{l_{ij}}{\sqrt{2a_t}\sigma(t)}, \frac{\sqrt{a_t}\|\mathbf{x}_i - \mathbf{x}_j\|}{\sqrt{2}\sigma(t)}, 3\right)\right) \quad (35)$$

where $\mathrm{CDF}_{\chi'}(x, \lambda, k)$ corresponds to the cumulative distribution function of the non-central chi distribution with $k$ degrees of freedom and non-centrality parameter $\lambda$. Practically, $\mathrm{CDF}_{\chi'}(x, \lambda, k)$ is expressed in terms of the MarcumQ function:

$$\mathrm{CDF}_{\chi'}(x, \lambda, k) = 1 - \mathrm{MarcumQ}_{\frac{k}{2}}(\lambda, x) \quad (36)$$

and for the special case of $k = 3$, $\mathrm{MarcumQ}_{\frac{k}{2}}(a, b)$ has the following closed form:

$$\mathrm{MarcumQ}_{\frac{3}{2}}(a, b) = \frac{1}{2}\left(\mathrm{erfc}\left(\frac{b-a}{\sqrt{2}}\right) + \mathrm{erfc}\left(\frac{b+a}{\sqrt{2}}\right)\right) + \frac{\sqrt{2}}{\sqrt{\pi}a}\sinh(ab)\, e^{-\frac{a^2+b^2}{2}} \quad (37)$$

where $\mathrm{erfc}(x)$ is the complementary error function. In practice, we use this form to implement a custom Python function that computes $\log(\mathrm{CDF}_{\chi'})$ in a numerically stable manner. This is necessary since, during sampling, accurate gradients of $\log(p(\{l_{ij} < \|\mathbf{x}_i^{(0)} - \mathbf{x}_j^{(0)}\| < u_{ij}\}|\mathbf{x}, t))$ must be calculated for the entire range of $\sigma(t)$.

Using these pairwise distance conditioners, four kinds of restraints are imposed on the geometry of sampled heavy-atom structures: 1) coarse-grained bead size restraints, 2) bond length restraints, 3) bond angle restraints and 4) Lennard-Jones pair distance restraints.

1. Coarse-grained bead size restraints

   When the positions of heavy atoms are denoised, heavy atoms can drift outside of their respective coarse-grained bead neighborhood, which is fixed and determined by the given coarse-structure. This creates discrepancies between the graph topology of the denoised structure and the one defined by the given coarse-structure. To prevent this, heavy atoms that belong to a given coarse-grained bead are restrained to remain in the neighborhood of the given bead position. This is ensured by imposing an upper bound on the distance between heavy-atom positions and their respective beads' positions. Specifically, given a set of coarse-grained bead positions $\{\mathbf{c}_j | j \in I\}$ and heavy-atom positions $\mathbf{x}_i$, the log-likelihood of this conditioner is given by:

$$\log(p(\|\mathbf{x}_i^{(0)} - \mathbf{c}_{\mathcal{C}(i)}\| < r_{\mathcal{C}(i)} + b|\mathbf{x}, t)) = \log\left(\mathrm{CDF}_{\chi'}\left(\frac{r_{\mathcal{C}(i)} + b}{\sqrt{a_t}\sigma(t)}, \frac{\sqrt{a_t}\|\mathbf{x}_i - \mathbf{c}_{\mathcal{C}(i)}\|}{\sigma(t)}, 3\right)\right)$$

where $r_{\mathcal{C}(i)}$ denotes the radius of the coarse-grained bead of atom $i$ and $b = 1\,\text{Å}$ is a buffer distance that was fine-tuned to relax the confining force of this conditioner. Note that compared to eq. (35), the denominators in the CDF are missing a factor of $\sqrt{2}$ since the conditioned distances only have a single free endpoint.

2. Bond length restraints

These restraints ensure that bond lengths in the final samples are within a physically realistic range. The allowable bond-length range for each bond type is defined a priori by analyzing the corresponding bond-length distribution in the structures of the training dataset. More precisely, for each bond $(i, j)$ between atom $i$ and $j$, lower and upper bounds of the bond length are determined by the sample mean $\hat{\mu}_{ij}$ and sample standard deviation $\hat{\sigma}_{ij}$ of that bond across the training dataset:

$$l_{ij} = \hat{\mu}_{ij} - n\hat{\sigma}_{ij} \quad (38)$$

$$u_{ij} = \hat{\mu}_{ij} + n\hat{\sigma}_{ij} \quad (39)$$

where $n$ is the sigma level, which controls the extension of the range in units of the sample standard deviation. Once these bounds are calculated, a conditioner is added for each bond in the protein's topology using eq. (35). A sigma level of $n = 5$ is used in all sampling runs.

3. Bond angle restraints

These restraints ensure that bond angles defined in the protein's force field topology are sampled in a realistic range. A bond angle $\theta_{ijk}$ is defined by a trio of atoms $(i, j, k)$; atoms $i, k$ are each bonded to atom $j$. To restrict the range of bond angles in the final samples, lower and upper bounds are imposed on the distance between atoms $i$ and $k$, i.e. $d_{ik} = \|\mathbf{x}_i^{(0)} - \mathbf{x}_k^{(0)}\|$. Since the bond length conditioner already imposes restraints on the range of $d_{ij}$ and $d_{jk}$, restricting the range of $d_{ik}$ will induce restraints on $\theta_{ijk}$. The lower and upper bounds of each $d_{ik}$ are defined analogously to the bond length conditioner using statistics of the corresponding distances estimated from the training dataset and the same sigma level ($n = 5$). An alternative method to restrict the bond angle range would be to impose conditions on the angle itself $\theta_{ijk}^{(0)}$. However, there exists no closed-form formula for $p(\theta_{ijk}^{(0)}|\mathbf{x}, t)$, preventing the fast calculation of gradients of the associated probability ($p(l_{ijk} < \theta_{ijk}^{(0)} < u_{ijk}|\mathbf{x}, t)$), which would be needed to define the conditioning score.

4. Lennard-Jones pair distance restraints

These restraints ensure that atoms do not approach each other too closely, which would otherwise result in high energies due to the $r^{-12}$ term in the Lennard-Jones (LJ) potential. To simplify these restraints, a global minimum is imposed on all LJ pair distances. This minimum is set to 2.0 Å, a value smaller than the closest LJ pair distance observed across all datasets. In addition, to accelerate the calculation of the conditioner's force, only distances between atom pairs identified as adjacent in the fixed adjacency matrix are conditioned.

To combine the set of all conditions $\{C_j\}$, eq. (25) suggests adding all conditioning gradients:

$$\nabla_{\mathbf{x}} \log(\hat{p}(\mathbf{x}, t|\{C_j\})) \approx \nabla_{\mathbf{x}} \log(\hat{p}(\mathbf{x}, t)) + \sum_j \nabla_{\mathbf{x}} \log(\hat{p}(C_j|\mathbf{x}, t)) \tag{40}$$

This is not true in general, since the set of conditioned variables that define the conditions can be correlated, which occurs when they are derived from a shared set of atom positions (e.g. lengths of two bonds with a common atom). However, since the sampling procedure is inherently noisy and since the conditions that we impose are relatively mild (e.g. sigma-level of 5 for the bond length and bond angle conditioners), we expect that summing the conditioners' gradients independently still provides beneficial steering of the sampling process, especially in the early phase of sampling. To verify this assumption, the energy of each conditioner was recorded when sampling the fine structure of the Nup98-12 system starting from coarse structures gathered from the training dataset (Supplementary Fig. 7). As expected, the energies of the CG bead size and the Lennard-Jones conditioners are non-zero during the early phase of sampling, where large length scales are denoised and vanish to zero as the sampling progresses towards smaller length scales. On the other hand, the energies of the bond length and bond angle conditioners vanish to zero at a slower rate, as they are restricting features associated with smaller length scales. Finally, the energy of all conditioners vanishes to zero before the diffusion energy, indicating that the model itself contributes the most to the gradients in the later stages of sampling.

#### B.6 Pruning

After generating a set of samples, structures that have non-physical values for any of the potential energies assessed are removed from the set. $N_E = 7$ energy types are used to prune the model ensembles, which include the bond energy, bond angle energy, dihedral angle energy, correction-map (CMAP) energy (when applicable), Lennard-Jones energy, Coulomb energy and total potential energy. For force fields without a CMAP correction, $N_E = 6$.

More specifically, let $\{\mathbf{x}^j\}$ denote the set of atomic positions in a given structure $j$ and let $E_k(\{\mathbf{x}^j\})$ denote the $k^{\text{th}}$ energy type ($k \in \{1, \ldots, N_E\}$) of the $j^{\text{th}}$ structure in a given

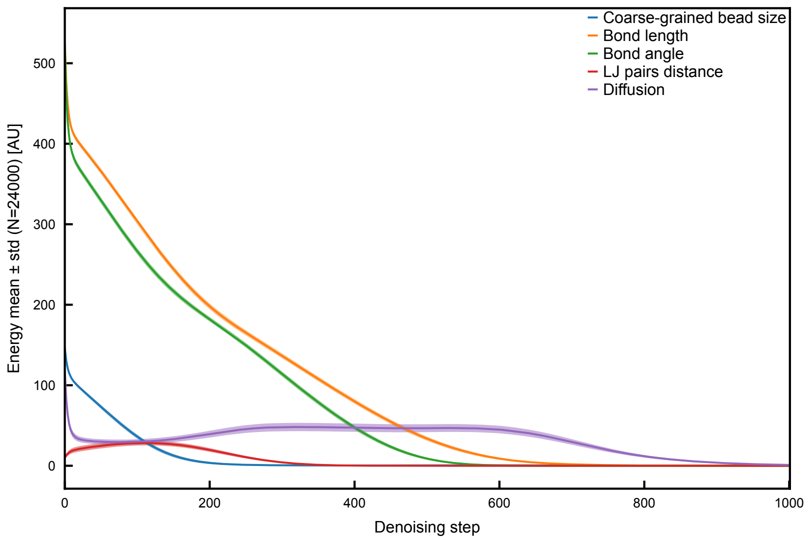

Supplementary Figure 7: **Energy of Sampling Conditioners**
The average energy of each conditioner is shown against the denoising step for the case of sampling the heavy-atom structure of the Nup98-12 system. The diffusion energy is also shown for relative comparison. The shaded area indicates one standard deviation around the mean. The energy statistics are evaluated using 24000 sampling trajectories, where 0.1% of the highest energies are omitted for computing each statistic at each denoising step. This is done to reduce the impact on the mean and standard deviation of outlier conditioner energies.

dataset. The Z-score of the $j^{\text{th}}$ structure on the $k^{\text{th}}$ energy is defined as:

$$z_k^j = \frac{E_k(\{\mathbf{x}^j\}) - \mu_k}{\sigma_k} \tag{41}$$

$$\mu_k = \frac{1}{|J_{\text{train}}|} \sum_{j \in J_{\text{train}}} E_k(\{\mathbf{x}^j\}), \qquad \sigma_k = \sqrt{\frac{1}{|J_{\text{train}}| - 1} \sum_{j \in J_{\text{train}}} (E_k(\{\mathbf{x}^j\}) - \mu_k)^2} \tag{42}$$

where $J_{\text{train}}$ denotes the set of all structures in the training dataset. Then, the set of structure indices that are retained after pruning $J_{\text{pruned}}$ is defined as follows:

$$J_{\text{pruned}} = \left\{ j \in J_{\text{model}} \middle| \forall k \in \{1, \ldots, N_E\}, |z_k^j| < 5 \right\} \tag{43}$$

where $J_{\text{model}}$ denotes the set of all structures in the model dataset. The percentage of pruned structures in the model ensembles used for the coarse and fine structure assessments is provided in Supplementary Table 2.

Supplementary Table 2: **Percentage of Pruned Samples in Model Ensembles**

| | Pruned Samples (%) |
|---|---|
| Nup98-12 | 0.67 |
| Nup98-24 | 0.73 |
| CLN025 | 0.48 |
| RS peptide | 0.53 |
| `(AAQAA)`$_3$ | 0.61 |

#### B.7 Technical Details

The codebase was written in Python (3.11.15). The network architecture was written with `torch(2.8.0)`[9] and the following packages were used for data processing and analysis: `matplotlib(3.8.4)`,[10] `mdanalysis(2.9.0)`,[11,12] `networkx(3.5)`,[13] `numpy(1.26.4)`,[14] `numba(0.61.2)`,[15] `openmm(8.1.1)`,[16] `scipy(1.11.3)`,[17] `umap_learn(0.5.7)`.[18]

Prior to training and sampling, functions and models were JIT-compiled using `torch.compile`. Automatic mixed precision (AMP) using 16-bit floating-point precision is used for predicting heavy-atom positions. The final predicted atom positions are represented using 32-bit floating-point precision. Finally, benchmarking was performed on a single workstation equipped with 8-core AMD Ryzen 7 5800X3D processor and an NVIDIA RTX 6000 Ada Generation graphics card.

### C Network Architecture

This section details the inner workings of the denoiser $D_\theta(\mathbf{x}, t)$ parameterized by $\theta$. Its architecture was inspired by previous work.[4,7,19] $D_\theta(\mathbf{x}, t)$ is modeled using a graph neural network (GNN) with SE(3) equivariant message-passing layers. The network architecture is summarized in algorithm 1, where colored functions contain trainable parameters and each layer $i$ consists of an equivariant graph convolution (EGC) followed by a cross-product module (CPM). The architecture of these modules, along with the preprocessing and post-processing steps, is described in the next sections. The denoiser's inputs are: 1) a set of noisy atom positions $\{\mathbf{x}_i : i \in I\}$ defining a protein structure with 2) amino acid sequence $S$; 3) $t$, the diffusion time, which informs the denoiser about the noise level in the input positions; 4) $A_{ij} \in \{0, 1\}$, the adjacency matrix where $A_{ij} = 1$ indicates that atoms $i$ and $j$ are connected when passing messages; 5) $\mathbf{p}_{\text{ff}}$, force field parameters (atom types, Lennard-Jones parameters and partial charges) and 6) $T$, the set of triplet node indices $T = \{(i, j, k) : i, j, k \in I\}$ that define the cross-product update (see Supplementary Appendix C.4).

**Algorithm 1** Denoiser architecture

**Require:** $\mathbf{x}$ (atom positions), $t$(diffusion time), $S$(protein sequence),
$A$(adjacency matrix), $T$(triplet node indices), $\mathbf{p}_{\text{ff}}$(force field parameters)

$\quad {}^{0}\mathbf{x}, \mathbf{h}, \mathbf{e}, T \leftarrow \text{PreProcess}(\mathbf{x}, t, S, \mathbf{p}_{\text{ff}})$

$\quad {}^{0}\mathbf{h} \leftarrow \text{NodeFeatureEncoder}(\mathbf{h})$

$\quad {}^{0}\mathbf{e} \leftarrow \text{EdgeFeatureEncoder}(\mathbf{e})$

$\quad$ **for** $i = 0, \ldots, N-1$ **do**

$\qquad {}^{i+1}\tilde{\mathbf{x}}, {}^{i+1}\tilde{\mathbf{h}}, {}^{i+1}\tilde{\mathbf{e}} \leftarrow \text{EGC}({}^{i}\mathbf{x}, {}^{i}\mathbf{h}, {}^{i}\mathbf{e}, A)$

$\qquad {}^{i+1}\mathbf{x}, {}^{i+1}\mathbf{h}, {}^{i+1}\mathbf{e} \leftarrow \text{CPM}({}^{i+1}\tilde{\mathbf{x}}, {}^{i+1}\tilde{\mathbf{h}}, {}^{i+1}\tilde{\mathbf{e}}, T)$

$\quad \mathbf{x}_{\text{out}} \leftarrow \text{PostProcess}({}^{N}\mathbf{x}, \mathbf{x}, t)$

$\quad$ **return** $\mathbf{x}_{\text{out}}$

Although the preconditioned denoiser form suggested by Karras et al.[4] (eq. (9)) controls the input and output variance, it breaks translation equivariance of the denoiser, i.e. $D_\theta(\mathbf{x} + \mathbf{y}, t) \neq D_\theta(\mathbf{x}, t) + \mathbf{y}$. To remedy this, we apply Karras' denoiser preconditioning on the centroid-free atom positions:

$$D_\theta(\mathbf{x}_i, t) = \mathbf{c} + c_{\text{skip}}(t)(\mathbf{x}_i - \mathbf{c}) + c_{\text{out}}(t)\left(F_\theta(c_{\text{in}}(t)\mathbf{x}_i, t) - c_{\text{in}}(t)\mathbf{x}_i\right), \qquad \mathbf{c} = \frac{1}{n}\sum_{i=1}^{n} \mathbf{x}_i \tag{44}$$

where $i$ indexes the $n$ atoms in a given structure and $\mathbf{c}$ is the centroid of the structure (see Supplementary Appendix C.1). With this modified preconditioned form, the SE(3) equivariance of $D_\theta$ is dictated by the equivariance of $F_\theta$.

#### C.1 Preprocessing

The main purpose of the preprocessing steps is to rescale the input positions and prepare features of the graph neural network. In general, a graph neural network has node features ($\mathbf{h}_i$) and edge features ($\mathbf{e}_{ij}$), where $i, j$ index nodes in the input graph. Although node positions $\mathbf{x}_i$ are strictly speaking node features, we will denote them differently than other node features $\mathbf{h}_i$ since their transformation is constrained by the SE(3) equivariance requirement.

First, the input node coordinates are scaled with $c_{\text{in}}(t)$ (see eq. (10)) to normalize their variance:

$$^{0}\mathbf{x}_i = c_{\text{in}}(t)\mathbf{x}_i \tag{45}$$

Second, five types of node features are defined:

1. Atom type, $a_i$

   This categorical feature encodes the atom type defined in the force-field-dependent topology. It is one-hot encoded and includes all possible atom types defined in the force field.

2. Residue type, $r_i$

   This categorical feature encodes the type of amino acid residue that each atom belongs to. It is one-hot encoded with a dimension of 20 to cover all 20 conventional amino acids.

3. Sequence type, $s_i$

   This ternary one-hot feature encodes whether the residue to which each atom belongs is terminal or non-terminal. Its definition is:

$$s_i = \begin{cases} (1,0,0) & \text{if atom } i \text{ is in the N-terminal residue} \\ (0,0,1) & \text{if atom } i \text{ is in the C-terminal residue} \\ (0,1,0) & \text{otherwise} \end{cases} \tag{46}$$

4. Backbone type, $bb_i$

   This binary feature encodes whether a given atom belongs to the backbone (1) or side-chain (0) of its amino acid residue.

5. Logged diffusion time $\tilde{t}$

   This floating-point scalar feature encodes the expected level of noise in the input positions in logarithmic scale: $\tilde{t} = \frac{\log(t)}{4}$.

For coarse-structure models, all node features, except $bb_i$, are concatenated and given to the graph neural network. For fine-structure models, the atom type, backbone type and logged diffusion time features are used. The omission of the residue type node feature for the fine-structure model forces the model to learn to position heavy atoms irrespective of the residues that they belong to. In other words, the model learns to utilize the chemical environment of each atom to predict its position.

Finally, the following edge features are defined:

1. Pairwise distance ${}^0d_{ij}$

   Distance between node $i$ and $j$, i.e. ${}^0d_{ij} = \|{}^0\mathbf{x}_i - {}^0\mathbf{x}_j\|$. Note that the scaled node positions ${}^0\mathbf{x}_i$ are used in defining the distance.

2. Inverse pairwise distance ${}^0d_{ij}^{-1}$

   Clipped inverse distance between node $i$ and $j$, i.e. ${}^0d_{ij}^{-1} = \frac{1}{{}^0d_{ij}+\epsilon}$ where $\epsilon = 0.5$ is used. $\epsilon \neq 0$ is necessary to avoid numerical instability in cases where two nodes are very close to one another, which can occur due to noise added to the positions.

3. Sequence position difference $spd_{ij}$

   This ternary one-hot-encoded feature encodes the absolute difference in position, along the protein sequence, of two given nodes $i, j$. Specifically,

$$spd_{ij} = \begin{cases} (1,0,0) & \text{if} \quad |\text{ResInd}(i) - \text{ResInd}(j)| = 0 \\ (0,1,0) & \text{if} \quad |\text{ResInd}(i) - \text{ResInd}(j)| = 1 \\ (0,0,1) & \text{if} \quad |\text{ResInd}(i) - \text{ResInd}(j)| > 1 \end{cases} \tag{47}$$

   where $\text{ResInd}(i)$ denotes the residue index of atom $i$.

4. Bond $b_{ij}$

   This binary feature encodes whether two atoms are bonded to each other as defined in the system's topology.

5. Lennard-Jones (LJ) parameters $[\epsilon_{ij}, \sigma_{ij}]$

   These floating-point features correspond to the force-field-dependent Lennard-Jones parameters that define the Lennard-Jones energy between atom $i$ and $j$:

$$V_{\text{LJ}}(\mathbf{x}_i, \mathbf{x}_j) = 4\epsilon_{ij}\left(\left(\frac{\sigma_{ij}}{d_{ij}}\right)^{12} - \left(\frac{\sigma_{ij}}{d_{ij}}\right)^{6}\right), \quad d_{ij} = \|\mathbf{x}_i - \mathbf{x}_j\| \tag{48}$$

6. Charge product $qq_{ij}$

   This floating-point feature corresponds to the product of the force-field-dependent partial charges of adjacent atoms, i.e. $qq_{ij} = q_i q_j$. This feature is meant to encode the charge-dependent term of the Coulomb interaction between pairs of atoms.

For coarse-structure models, only the pairwise distance, inverse pairwise distance and sequence position difference are used as edge features; fine-structure models use all six edge features. The preprocessing steps are summarized in algorithm 2. The construction of the triplet node index sets is explained in Supplementary Appendix C.4.

**Algorithm 2** Denoiser preprocessing

**Require:** $\mathbf{x}$ (atom positions), $t$ (diffusion time), $S$(protein sequence), $\mathbf{p}_{\mathrm{ff}}$ (force field parameters)

**for** $i = 1, \ldots, n$ **do**

  ${}^0\mathbf{x}_i \leftarrow c_{\mathrm{in}}(t)\mathbf{x}_i$

$a, r, s, \tilde{t} \leftarrow ComputeNodeFeatures(\mathbf{p}_{\mathrm{ff}}, S, t)$

${}^0d, {}^0d^{-1}, spd, b, \epsilon, \sigma, qq \leftarrow ComputeEdgeFeatures({}^0\mathbf{x}, \mathbf{p}_{\mathrm{ff}}, S)$

### Concatenate node and edge features

**if** coarse-structure model **then**

  $\mathbf{h} \leftarrow [a, r, s, \tilde{t}]$

  $\mathbf{e} \leftarrow [{}^0d, {}^0d^{-1}, spd]$

  $T \leftarrow ComputeCoarseGrainedBeadTripletIndices(S)$

**else**

  $\mathbf{h} \leftarrow [a, bb, \tilde{t}]$

  $\mathbf{e} \leftarrow [{}^0d, {}^0d^{-1}, spd, b, \epsilon, \sigma, qq]$

  $T \leftarrow ComputeHeavyAtomTripletIndices(S)$

**return** ${}^0\mathbf{x}, \mathbf{h}, \mathbf{e}$, T

#### C.2 Feature Encoders

Before evaluating the graph convolution layers, the input node and edge features are passed through encoders to map them into their respective latent spaces. The dimension of the node and edge features' latent space is kept constant thereafter. The architecture of the encoders is given by a single fully-connected layer with a learnable bias (see algorithm 3).

**Algorithm 3** Feature Encoders

**Require:** $\mathbf{h}$ (input node features), $\mathbf{e}$ (input edge features)

${}^0\mathbf{h} \leftarrow \mathrm{LinearWithBias}(\mathbf{h})$

${}^0\mathbf{e} \leftarrow \mathrm{LinearWithBias}(\mathbf{e})$

**return** ${}^0\mathbf{h}, {}^0\mathbf{e}$

#### C.3 Equivariant Graph Convolution

The core elements of the graph neural network architecture are the equivariant graph convolution (EGC) layers. These layers update each given node's position by accumulating SE(3) equivariant messages sent from all adjacent nodes. In the coarse-structure model, nodes are adjacent to all other nodes (fully-connected graph), while nodes in the fine-structure model exchange messages with a local neighborhood of nodes defined by the adjacency of the given coarse-grained beads (see Supplementary Appendix A). In addition, node features and edge features are updated using conventional graph convolutions. Details of the EGC architecture are given in algorithm 4.

**Algorithm 4** Equivariant Graph Convolution

**Require:** $\mathbf{x}$ (atom positions), $\mathbf{h}$ (node features), $\mathbf{e}$ (edge features), $A$(adjacency matrix)

  # Concatenate input of $\phi$ modules (see algorithm 7) for each message direction.
  **for** $i = 1, \dots, n$ **do**
    $\mathcal{N}(i) \leftarrow \{j | A_{ij} = 1\}$ # Construct set of neighboring node indices
    **for** $j \in \mathcal{N}(i)$ **do**
      $\mathbf{q}_{ij} \leftarrow [\mathbf{h}_i, \mathbf{h}_j, \mathbf{e}_{ij}, \|\mathbf{x}_i - \mathbf{x}_j\|]$

  # Update node positions
  **for** $i = 1, \dots, n$ **do**
    # Compute position attention weights
    **for** $j \in \mathcal{N}(i)$ **do**
      $\tilde{w}_{ij} = \phi_{w,x}(\mathbf{q}_{ij})$
    $W_i \leftarrow \sum_k \exp(\tilde{w}_{ik})$
    **for** $j \in \mathcal{N}(i)$ **do**
      $w_{ij} = \frac{\exp(\tilde{w}_{ij})}{W_i}$
    $\mathbf{x}_i \leftarrow \mathbf{x}_i + \sum_{j \in \mathcal{N}(i)} (\mathbf{x}_i - \mathbf{x}_j) w_{ij} \phi_x(\mathbf{q}_{ij})$ # SE(3) equivariant update

  # Update node features
  **for** $i = 1, \dots, n$ **do**
    # Compute node feature attention weights
    **for** $j \in \mathcal{N}(i)$ **do**
      $\tilde{w}_{ij} = \phi_{w,\text{msg}}(\mathbf{q}_{ij})$
    $W_i \leftarrow \sum_k \exp(\tilde{w}_{ik})$
    **for** $j \in \mathcal{N}(i)$ **do**
      $w_{ij} = \frac{\exp(\tilde{w}_{ij})}{W_i}$
    $\mathbf{h}_i \leftarrow \mathbf{h}_i + \phi_h([\mathbf{h}_i, \sum_{j \in \mathcal{N}(i)} w_{ij} \phi_{\text{msg}}(\mathbf{q}_{ij})])$

  # Update edge features
  **for** $i = 1, \dots, n$ **do**
    **for** $j \in \mathcal{N}(i)$ **do**
      **if** $i < j$ **then**
        $\mathbf{e}_{ij} \leftarrow \mathbf{e}_{ij} + \phi_e([|\mathbf{h}_i - \mathbf{h}_j|, \mathbf{e}_{ij}, \|\mathbf{x}_i - \mathbf{x}_j\|])$
      **else**
        $\mathbf{e}_{ij} \leftarrow \mathbf{e}_{ji}$
  **return** $\mathbf{x}, \mathbf{h}, \mathbf{e}$

#### C.4 Cross-product Module

As shown previously,[20,21] model layers that perform tensor products of irreducible representations of rotation groups efficiently learn 3D geometric features. For learning protein structures, including these products in the model architecture is essential to properly predict their chiral features.

The cross-product module utilizes one type of these tensor products: the cross product of two vectors. Specifically, it updates node positions using a weighted sum of cross products of neighboring nodes' positions. The index set of node triplets $T$ that form the accumulated cross products are fixed and determined by the molecular graph of the given protein and the type of the predicted structure (coarse or fine).

For fine-structure models, the set of triplets is built by enumerating all distinct pairs of atoms in the bonded neighborhood of each atom in the molecular graph. Specifically, if $B_i$ denotes the index set of atoms bonded to atom $i$ in the molecular graph $G$, the set of cross-product triplet indices $T_{\mathrm{HA}}$ is:

$$T_{\mathrm{HA}} = \{(i, j, k) | i \in G, j, k \in B_i, j \neq k\} \tag{49}$$

For the coarse-structure model, the triplets are formed out of combinations of two pairs of adjacent backbone ($c_i$) or side-chain ($s_i$) coarse-grained beads that share a common coarse-grained bead. Two coarse-grained beads are considered adjacent when the residues that they belong to are contiguous along the protein sequence. In addition, for defining the set of triplets, coarse-grained beads of terminal residues are also treated as adjacent and coarse-grained beads that represent side chains are not considered adjacent in order to reduce the combinatorial complexity of the triplets' definition.

Specifically, let $\tilde{A}_{\mathrm{CG}}$ denote the adjacency matrix used to define the triplet indices $T_{\mathrm{CG}}$ for the ordered set of coarse-grained beads $\{c_1, s_1, c_2, s_2, \ldots, c_N, s_N\}$ in a sequence of $N$ residues.

Applying the aforementioned rules of adjacency, $\tilde{A}_{\text{CG}}$ takes the following form:

$$\tilde{A}_{\text{CG}} = \begin{pmatrix} 0 & 1 & 0 & 1 & 0 & \dots & 1 & 1 \\ 1 & 0 & 1 & 0 & 0 & \dots & 1 & 0 \\ 1 & 1 & 0 & 1 & 1 & \dots & 0 & 0 \\ 1 & 0 & 1 & 0 & 1 & \dots & 0 & 0 \\ 0 & 0 & 1 & 1 & 0 & \dots & 0 & 0 \\ \vdots & \vdots & \vdots & \vdots & \vdots & \ddots & \vdots & \vdots \\ 1 & 1 & 0 & 0 & 0 & \dots & 0 & 1 \\ 1 & 0 & 0 & 0 & 0 & \dots & 1 & 0 \end{pmatrix} \tag{50}$$

Denoting $\tilde{G}_{\text{CG}}$ as the associated graph with adjacency matrix $\tilde{A}_{\text{CG}}$, the set of coarse-grained bead triplet indices is defined in a way similar to the heavy-atom triplets:

$$T_{\text{CG}} = \{(i, j, k) | i \in \tilde{G}_{\text{CG}}, j, k \in B_i, j \neq k\} \tag{51}$$

$$B_i = \{j | i, j \in \tilde{G}_{\text{CG}}, \tilde{A}_{\text{CG},ij} = 1\} \tag{52}$$

where $B_i$ denotes the set of all CG bead indices connected to the $i^{\text{th}}$ bead according to $\tilde{A}_{\text{CG}}$.

With these definitions of the node triplets, the value of the cross-product module is assessed with ablation experiments. Refer to Supplementary Appendix E.7 for this analysis.

#### C.5 Post-processing

As shown in eq. (44), the output positions of the neural network $F_\theta$ are combined with the input positions $\mathbf{x}$ to ensure $SE(3)$ equivariance of the denoiser $D_\theta$. These computations are included in the post-processing step of the neural network's output as detailed in algorithm 6.

**Algorithm 5** Cross-product Module

**Require:** $\mathbf{x}$ (atom positions), $\mathbf{h}$ (node features), $T$(set of triplet indices)

  # Concatenate input of $\phi$ module for each triplet.
  **for** $(i, j, k) \in T$ **do**
    $\mathbf{q}_{ijk} \leftarrow [\mathbf{h}_i, \mathbf{h}_j, \mathbf{h}_k, \|\mathbf{x}_j - \mathbf{x}_i\|, \|\mathbf{x}_k - \mathbf{x}_i\|, \|\mathbf{x}_k - \mathbf{x}_j\|]$

  # Update node positions
  **for** $i = 1, \ldots, n$ **do**
    # Compute attention weights
    **for** $(l, j, k) \in T, l = i$ **do**
      $\tilde{w}_{ijk} = \phi_{w,\text{cp}}(\mathbf{q}_{ijk})$
    $W_i \leftarrow \sum_{(l,m,n)\in T, l=i} \exp(\tilde{w}_{lmn})$
    **for** $(l, j, k) \in T, l = i$ **do**
      $w_{ijk} = \frac{\exp(\tilde{w}_{ijk})}{W_i}$
    $\mathbf{x}_i \leftarrow \mathbf{x}_i + \sum_{(l,j,k)\in T, l=i} (\mathbf{x}_j - \mathbf{x}_i) \times (\mathbf{x}_k - \mathbf{x}_i) w_{ijk} \phi_{\text{cp}}(\mathbf{q}_{ijk})$ # SE(3) equivariant update
  **return** $\mathbf{x}$

**Algorithm 6** Denoiser post-processing

**Require:** $\mathbf{x}$ (input positions), $\mathbf{y}$ (networks' output positions), $t$ (diffusion time)
  $\mathbf{c} \leftarrow \frac{1}{n} \sum_{i=1}^{n} \mathbf{x}_i$ # Compute centroid of input structure
  **for** $i = 1, \ldots, n$ **do**
    $\mathbf{z}_i \leftarrow \mathbf{c} + c_{\text{skip}}(t)(\mathbf{x}_i - \mathbf{c}) + c_{\text{out}}(t)\,(\mathbf{y}_i - c_{\text{in}}(t)\mathbf{x}_i)$
  **return** $\mathbf{z}$

#### C.6 Architectures of Feature Modules

In all layers of the models' graph neural network, $\phi_{\cdot}(\cdot)$ modules are used as universal function approximators to evaluate attention weights and latent feature updates. These modules all share a basic architecture based on fully-connected layers, Sigmoid Linear Units (SiLU) activation functions and layer normalization layers (NormLayer), as detailed in algorithm 7. Note that each layer of the graph neural network possesses a unique set of $\phi$ modules, i.e. modules do not share weights.

Supplementary Table 3: **Model Hyperparameters**
The number of parameters of the coarse-structure and fine-structure models depends on the chosen force field and is approximately $\sim 750k$ across all systems. The exact values provided in the table correspond to the number of parameters for the Nup98-12 models.

| Symbol | Description | Coarse-structure Model | Fine-structure Model |
|---|---|---|---|
| $N_{\text{layers}}$ | Number of layers | 10 | 10 |
| $[t_{\min}, t_{\max}]$ | Noise schedule limits (Å) | $[0.1, 50]$ | $[0.002, 5]$ |
| $\mathbf{h}_{\text{in}}$ | Input node features | $[a_i, r_i, s_i]$ | $[a_i, bb_i]$ |
| $\mathbf{e}_{\text{in}}$ | Input edge features | $[{}^0d_{ij}, {}^0d_{ij}^{-1}, spd_{ij}]$ | $[{}^0d_{ij}, {}^0d_{ij}^{-1}, spd_{ij}, b_{ij}, \epsilon_{ij}, \sigma_{ij}, qq_{ij}]$ |
| $d_h$ | Latent node feat. dim. | 64 | 64 |
| $d_e$ | Latent edge feat. dim. | 64 | 64 |
| $d_{\text{cp}}$ | CPM hidden dimension | 64 | 64 |
| | Number of parameters | 756,136 | 754,984 |

**Algorithm 7** $\phi$ modules definition (PyTorch[9])

**Require:** $d_h$ (node feature dimension), $d_e$ (edge feature dimension), $d_{\text{msg}}$ (message dimension), $d_{\text{cp}}$ (hidden dimension of cross-product module)

$$
\begin{aligned}
d_{\text{msg,in}}, d_{x,\text{in}} &\leftarrow 2d_h + d_e + 1\\
d_{e,\text{in}} &\leftarrow d_h + d_e + 1\\
d_{\text{cp,in}} &\leftarrow 3d_h + 3\\
\phi_{\text{msg}} &\leftarrow \text{Sequential}(\ \text{Linear}(d_{\text{msg,in}}, d_{\text{msg}}, \text{bias} = \mathit{False}), \text{SiLU}(),\\
&\qquad \text{NormLayer}(d_{\text{msg}}, \text{bias} = \mathit{False}),\\
&\qquad \text{Linear}(d_{\text{msg}}, d_{\text{msg}}, \text{bias} = \mathit{False}), \text{SiLU}(),\\
&\qquad \text{NormLayer}(d_{\text{msg}}, \text{bias} = \mathit{False}))\\
\phi_{w,\text{msg}} &\leftarrow \text{Sequential}(\ \text{Linear}(d_{\text{msg}}, 1, \text{bias} = \mathit{False}), \text{SiLU}())\\
\phi_h &\leftarrow \text{Sequential}(\ \text{Linear}(d_{\text{h,in}} + d_{\text{msg}}, d_{\text{h}}, \text{bias} = \mathit{False}), \text{SiLU}(),\\
&\qquad \text{NormLayer}(d_{\text{h}}, \text{bias} = \mathit{False}),\\
&\qquad \text{Linear}(d_{\text{h}}, d_{\text{h}}, \text{bias} = \mathit{False}))\\
\phi_e &\leftarrow \text{Sequential}(\ \text{Linear}(d_{e,\text{in}}, d_e, \text{bias} = \mathit{False}), \text{SiLU}(),\\
&\qquad \text{NormLayer}(d_e, \text{bias} = \mathit{False}),\\
&\qquad \text{Linear}(d_e, d_e, \text{bias} = \mathit{False}))\\
\phi_x &\leftarrow \text{Sequential}(\ \text{Linear}(d_{x,\text{in}}, d_{\text{msg}}, \text{bias} = \mathit{False}), \text{SiLU}(),\\
&\qquad \text{NormLayer}(d_{\text{msg}}, \text{bias} = \mathit{False}),\\
&\qquad \text{Linear}(d_{\text{msg}}, d_{\text{msg}}, \text{bias} = \mathit{False}), \text{SiLU}(),\\
&\qquad \text{NormLayer}(d_{\text{msg}}, \text{bias} = \mathit{False}),\\
&\qquad \text{Linear}(d_{\text{msg}}, 1, \text{bias} = \mathit{False}))\\
\phi_{w,x} &\leftarrow \text{Sequential}(\ \text{Linear}(d_{x,\text{in}}, 1, \text{bias} = \mathit{False}), \text{SiLU}())\\
\phi_{\text{cp}} &\leftarrow \text{Sequential}(\ \text{Linear}(d_{\text{cp,in}}, d_{\text{cp}}, \text{bias} = \mathit{False}), \text{SiLU}(),\\
&\qquad \text{NormLayer}(d_{\text{cp}}, \text{bias} = \mathit{False}),\\
&\qquad \text{Linear}(d_{\text{cp}}, d_{\text{cp}}, \text{bias} = \mathit{False}), \text{SiLU}(),\\
&\qquad \text{NormLayer}(d_{\text{cp}}, \text{bias} = \mathit{False}),\\
&\qquad \text{Linear}(d_{\text{cp}}, 1, \text{bias} = \mathit{False}))\\
\phi_{w,\text{cp}} &\leftarrow \text{Sequential}(\ \text{Linear}(d_{\text{cp,in}}, 1, \text{bias} = \mathit{False}), \text{SiLU}())
\end{aligned}
$$

**return** $\phi_{\text{msg}}, \phi_{w,\text{msg}}, \phi_h, \phi_e, \phi_x, \phi_{w,x}, \phi_{\text{cp}}, \phi_{w,\text{cp}}$

### D Datasets

We trained Zephyr on five model systems (Supplementary Table 4). The MD simulations of Nup98-12, Nup98-24, RS peptide and $\texttt{(AAQAA)}_3$ systems were carried out in this study, with the simulation details provided below. Zephyr was also trained on MD trajectories of the protein CLN025 from Lindorff-Larsen et al.,[22] which were provided by D.E. Shaw Research. These simulations were carried out with the CHARMM22* force field.

For all the systems in this study, a requirement was that the system be small enough that the iterative process of training and sampling Zephyr not be too computationally expensive. The systems were chosen to cover a range of size, net charge and structural propensity. The RS peptide is an example of a disordered serine/arginine-rich sequence, which has been studied for its relevance in RNA metabolism.[23] The RS peptide was chosen for this work because it has been extensively characterized in MD simulations and has been used to assess force-field accuracy.[23–25] The $\texttt{(AAQAA)}_3$ peptide was chosen because it has seen significant use as a test system for MD force-field evaluation, as well as serving as a model for studying helix-coil transitions.[26,27] The ten-residue protein CLN025 is one of the smallest proteins that can fold.[28] While the other systems in this study are model IDPs, CLN025 allows us to test whether Zephyr can generate folded structures when trained exclusively on structures from the unfolded state and vice versa (see Supplementary Appendix F.5). The Nup98-12 and Nup98-24 peptides each comprise one (Nup98-12) or two (Nup98-24) repeats of a 12-residue sequence designed by Ng et al.[29] This sequence was designed to recapitulate the behavior of the IDP Nup98, a member of the FG-nucleoporin family of proteins found in the nuclear pore complex.[29] The interactions between FG-nucleoporins and karyopherin proteins in the nuclear pore complex regulate transport of molecular cargo across the nuclear membrane.[30] The Nup98-12 and Nup98-24 peptides were chosen for this work because we expect them to mimic the disordered nature of Nup98 based on prior work.[29,31] Their repetitive sequences allow us to probe the performance of Zephyr on systems of different sizes while keeping the sequence composition the same.

For each system, a fully extended conformation was generated using UCSF ChimeraX[32] with backbone dihedral angles of (−180°,−180°) for each residue. All further processing and simulation steps used GROMACS (version 2023.2)[33,34] and force fields specified in Supplementary Table 6. Hydrogen atoms were added to each structure using GROMACS' `pdb2gmx`, setting the termini of each system according to Supplementary Table 4. All side chains were assigned standard protonation states at neutral pH.

Supplementary Table 4: **Model Systems**
For each of the model systems, the amino acid sequence is provided along with the N- and C-termini in the topology used in the MD simulations.

| System Name | Sequence | N-/C-termini |
|---|---|---|
| Nup98-12 | GGLFGGNTQPAT | $NH_2$/COOH |
| Nup98-24 | (GGLFGGNTQPAT)$_2$ | $NH_2$/COOH |
| CLN025 | YYDPETGTWY | $NH_3^+$/$COO^-$ |
| RS peptide | GAMGPSYGRSRSRSRSRSRSRSRS | $NH_3^+$/$COO^-$ |
| (AAQAA)$_3$ | (AAQAA)$_3$ | $NH_3^+$/$COO^-$ |

The structure was then placed in a rhombic dodecahedral box with periodic boundary conditions, with the box size set such that the minimum distance from the protein to the box edge was 10 Å. The system boxes were solvated; $Na^+$ and $Cl^-$ ions were added to a concentration of 0.15 M such that the system was neutral (Supplementary Table 5).

For simulations run with CHARMM36m and CHARMM22*, the CHARMM-modified TIP3P water model[35] was used. For the Amber14sb simulations, the TIP3P water model[36] was used. In all simulations, the water molecules were constrained using the SETTLE algorithm[37] and bonds involving hydrogen atoms were constrained using the LINCS algorithm.[38] Long-range electrostatics were computed using particle-mesh Ewald summation[39] with fourth-order interpolation and a grid spacing of 1.2 Å. The Verlet scheme was used for neighbor searching. All simulations were run with a timestep of 2 fs. Short-range and van der Waals interactions were computed with a cutoff of 9.5 Å.

Equilibration followed the same procedure for each system, beginning with energy minimization until the maximum force was smaller than 1000.0 kJ/mol/nm or 50,000 steps had

Supplementary Table 5: **MD Box Details**
The systems simulated in this work, including water molecules, ions and box volume. The original CLN025 simulation details are not reported here, as the trajectories were provided by D.E. Shaw Research; see[22] for simulation details. The CLN025 row in this table refers to the simulation box used for the equilibration runs described in Supplementary Appendix E.9, not the original simulation.

| System Name | Force Field | Water molecules | $Na^+$ ions / $Cl^-$ ions | Volume [$nm^3$] |
|---|---|---|---|---|
| Nup98-12 | CHARMM36m | 6,249 | 18 / 18 | 186.77 |
| Nup98-24 | CHARMM36m | 29,429 | 82 / 82 | 884.45 |
| CLN025 | CHARMM22* | 4,089 | 14 / 12 | 122.68 |
| RS peptide | CHARMM36m | 13,146 | 37 / 45 | 394.06 |
| `(AAQAA)`$_3$ | Amber14sb | 9,562 | 27 / 27 | 294.74 |

been completed. For each of the following equilibration steps, the v-rescale thermostat[40] was used. Energy minimization was followed by 10 ns NPT equilibration using the Berendsen barostat[41] and a further 10 ns NPT equilibration using the Parrinello-Rahman barostat.[42] The final structure from the NPT equilibration step was run in a high-temperature (450 K) NVT simulation for 12 ns. From the high-temperature NVT simulation, 12 structures were extracted spaced 1000 frames (1 ns) apart using GROMACS' `trjconv` command. Each of these 12 structures was used to initialize an independent MD trajectory; for the Nup98-24, RS peptide and `(AAQAA)`$_3$ systems each production run was 2 µs long, and for the Nup98-12 system they were 4 µs long. All production runs were carried out in the NPT ensemble using the Parrinello-Rahman barostat[42] and the v-rescale thermostat.[40] All simulations were set to a pressure of 1 bar and a temperature of 298 K (except for the high-temperature NVT run at 450 K).

MD simulations of the Nup98-12 system with acetyl and N-methylamide capping groups were carried out as well using the same procedure as for all other simulations. These are discussed in detail in Supplementary Appendix E.8.

After gathering the molecular dynamics trajectories for the five systems mentioned above, the atomic coordinates associated with the protein structure were extracted at intervals of 1 ns to minimize temporal correlation between the structures. The train/validation/test splits of the datasets were formed by selecting all downsampled structures from a random

Supplementary Table 6: **System Topologies**
For each system, the following information is provided: the force field used in the simulations, the number of residues, the number of atoms, the number of bonds, the number of bond angles and the number of dihedral angles in the topology. The numbers in parentheses indicate the number of geometric features that exclude hydrogen atoms, i.e. the heavy-atom features.

| System | Force Field | Res. | Atoms | Bonds | Bond Angles | Dih. Angles |
|---|---|---|---|---|---|---|
| Nup98-12 | CHARMM36m | 12 | 153(79) | 154(80) | 273(108) | 315(118) |
| Nup98-24 | CHARMM36m | 24 | 303(157) | 306(160) | 545(217) | 633(242) |
| CLN025 | CHARMM22* | 10 | 166(93) | 171(98) | 299(136) | 390(162) |
| RS peptide | CHARMM36m | 24 | 377(187) | 378(188) | 662(245) | 776(264) |
| $(\texttt{AAQAA})_3$ | Amber14sb | 15 | 174(88) | 173(87) | 309(119) | 351(113) |

subset of trajectories. In other words, datasets are randomly split at the trajectory level, not the structure level. Details of the datasets and their splits are given in Supplementary Table 7.

Supplementary Table 7: **Dataset Details**
The total simulated time across trajectories, number of structures and training/validation/test proportion are given for the main datasets and minimal datasets constructed for each system. All trajectories for a given system have the same length. The minimal datasets are used to test the ability of Zephyr to train on scarce data (Supplementary Appendix F.3). The CLN025 dataset was provided by D.E. Shaw Research[22] as 53 trajectories of 2 µs each, which originated from a single trajectory of 106 µs. The number of samples in the model dataset is equal to the number of samples in the training dataset for each system.

| | time (µs) | structures | train/valid/test | | train/valid/test (minimal) | |
|---|---|---|---|---|---|---|
| | | | trajectories | % | trajectories | % |
| Nup98-12 | 48 | 48,000 | 6/1/5 | 50.0/8.3/41.7 | 1/1/10 | 8.3/8.3/83.3 |
| Nup98-24 | 24 | 24,000 | 6/1/5 | 50.0/8.3/41.7 | 1/1/10 | 8.3/8.3/83.3 |
| CLN025 | 106 | 105,947 | 24/5/24 | 45.3/9.4/45.3 | 5/5/43 | 9.4/9.4/81.1 |
| RS peptide | 24 | 24,000 | 6/1/5 | 50.0/8.3/41.7 | 1/1/10 | 8.3/8.3/83.3 |
| $(\texttt{AAQAA})_3$ | 24 | 24,000 | 6/1/5 | 50.0/8.3/41.7 | 1/1/10 | 8.3/8.3/83.3 |

Supplementary Table 8: **Model Dataset Size Details**
The reported numbers correspond to the total number of structures generated by the model, prior to pruning.

| System | Structures (Main) | Structures (Minimal) |
|---|---|---|
| Nup98-12 | 24,000 | 40,000 |
| Nup98-24 | 12,000 | 20,000 |
| CLN025 | 47,976 | 85,957 |
| RS peptide | 12,000 | 20,000 |
| $\texttt{(AAQAA)}_3$ | 12,000 | 20,000 |

| | |
|---|---|
| CLN025 (thresholded) | 25,000 |
| Nup98-12 ($\psi$ scans) | 24,000 |

### E Assessment of Ensemble Accuracy

#### E.1 Bond Length and Bond Angle Z-score

Each tested system includes many bond lengths and bond angles (Supplementary Table 6), which complicates the comparison of the generated ensembles and MD ensembles with respect to these features. As all bond length and bond angle distributions are unimodal, the Z-score of each feature, computed relative to the training set statistics, is used to pool them into a single standardized distribution. Specifically, let $\{\mathbf{x}^j\}$ denote the set of atomic positions in a given structure $j$ and let $f_k(\{\mathbf{x}^j\})$ denote the $k^{\text{th}}$ feature (bond length or bond angle) of the $j^{\text{th}}$ structure in a given dataset. The corresponding Z-score $z_k^j$ is defined as:

$$z_k^j = \frac{f_k(\{\mathbf{x}^j\}) - \mu_k}{\sigma_k} \tag{53}$$

$$\mu_k = \frac{1}{|J_{\text{train}}|} \sum_{j \in J_{\text{train}}} f_k(\{\mathbf{x}^j\}), \qquad \sigma_k = \sqrt{\frac{1}{|J_{\text{train}}| - 1} \sum_{j \in J_{\text{train}}} (f_k(\{\mathbf{x}^j\}) - \mu_k)^2} \tag{54}$$

where $J_{\text{train}}$ denotes the set of indices corresponding to structures in the training split of the dataset.

#### E.2 Energy Metrics

In addition to geometric features, we compare the generated and MD ensembles using energy metrics based on the potential energy terms defined in the empirical force fields used in MD simulations. While structural metrics are used to evaluate the accuracy of generated structures across different length scales, energy metrics provide complementary information because they are nonlinear functions of the underlying geometric features (e.g. pairwise distances, bond angles, dihedral angles). Consequently, they provide a sensitive measure of the variability of the generated ensembles. Only intramolecular (protein-only) energy terms are considered. Unlike the potential energies computed during MD simulations, the energy terms used in our assessment exclude protein-solvent and solvent-solvent interactions, allowing the structural ensembles to be evaluated irrespective of solvent.

The various energies that are assessed are listed below:

1. Bond Energy

   For a given bond between two atoms positioned at $\mathbf{x}_i$ and $\mathbf{x}_j$, the bond energy is calculated as:

$$V_{\mathrm{B}}(\mathbf{x}_i, \mathbf{x}_j) = \frac{k_{ij}}{2} \left( \|\mathbf{x}_i - \mathbf{x}_j\| - l_{ij}^0 \right)^2 \tag{55}$$

   where $k_{ij}, l_{ij}^0$ are parameters defined in the given force field and depend on the atom types of the two atoms forming the bond.

2. Bond Angle Energy

   Given a triplet of atoms positioned at $\mathbf{x}_i, \mathbf{x}_j$ and $\mathbf{x}_k$, let $\theta_{ijk}$ denote the angle between the vectors $\mathbf{d}_{ij} = \mathbf{x}_i - \mathbf{x}_j$ and $\mathbf{d}_{jk} = \mathbf{x}_k - \mathbf{x}_j$, i.e. $\theta_{ijk} = \arccos\left(\frac{\mathbf{d}_{ij} \cdot \mathbf{d}_{jk}}{\|\mathbf{d}_{ij}\| \|\mathbf{d}_{jk}\|}\right)$. In general, the bond angle energy depends on the force field used. For systems simulated with an

Amber force field, the bond angle energy is a simple harmonic potential:

$$V_{\rm BA}(\mathbf{x}_i, \mathbf{x}_j, \mathbf{x}_k) = \frac{k_{ijk}}{2}\left(\theta_{ijk} - \theta^0_{ijk}\right)^2 \tag{56}$$

where $k_{ijk}, \theta^0_{ijk}$ are parameters defined in the force field that depend on the types of atoms that define the bond angle.

For the CHARMM force fields used here, the bond angle energy is given by a Urey-Bradley potential:

$$V_{\rm BA}(\mathbf{x}_i, \mathbf{x}_j, \mathbf{x}_k) = \frac{k^\theta_{ijk}}{2}\left(\theta_{ijk} - \theta^0_{ijk}\right)^2 + \frac{k^r_{ijk}}{2}\left(\|\mathbf{x}_i - \mathbf{x}_k\| - d^0_{ik}\right)^2 \tag{57}$$

where $k^\theta_{ijk}, \theta^0_{ijk}, k^r_{ijk}, d^0_{ik}$ are parameters defined in the force field that depend on the types of atoms that define the bond angle.

3. Dihedral Angle Energy

Given a quartet of atoms positioned at $\mathbf{x}_i, \mathbf{x}_j, \mathbf{x}_k$ and $\mathbf{x}_l$, let $\phi_{ijkl}$ denote the dihedral angle between the two planes formed by $\mathbf{x}_i, \mathbf{x}_j, \mathbf{x}_k$ and $\mathbf{x}_j, \mathbf{x}_k, \mathbf{x}_l$. The dihedral angle $\phi_{ijkl}$ is defined as follows:

$$\mathbf{d}_{ij} = \mathbf{x_j} - \mathbf{x}_i, \qquad \mathbf{d}_{jk} = \mathbf{x_k} - \mathbf{x}_j, \qquad \mathbf{d}_{kl} = \mathbf{x_l} - \mathbf{x}_k \tag{58}$$

$$\mathbf{u}_{ijk} = \mathbf{d}_{ij} \times \mathbf{d}_{jk}, \qquad \mathbf{u}_{jkl} = \mathbf{d}_{jk} \times \mathbf{d}_{kl} \tag{59}$$

$$x = \frac{\mathbf{u}_{ijk} \cdot \mathbf{u}_{jkl}}{\|\mathbf{u}_{ijk}\|\|\mathbf{u}_{jkl}\|}, \qquad y = \frac{\mathbf{d}_{jk} \cdot (\mathbf{u}_{ijk} \times \mathbf{u}_{jkl})}{\|\mathbf{d}_{jk}\|\|\mathbf{u}_{ijk}\|\|\mathbf{u}_{jkl}\|} \tag{60}$$

$$\phi_{ijkl} = \arctan2(y, x), \qquad -\pi < \phi_{ijkl} \leq \pi \tag{61}$$

where $\arctan2(y, x)$ is the two-argument arctangent function, which corresponds to the argument of the complex number $x + iy$. There are two types of dihedral angle energy calculated for the systems that we studied. First, the periodic dihedral angle energy is calculated as follows:

$$V^{\text{per}}_{\text{dihedral angle}}(\mathbf{x}_i, \mathbf{x}_j, \mathbf{x}_k, \mathbf{x}_l) = k_\phi(1 + \cos(n\phi_{ijkl} - \phi_0)) \tag{62}$$

where $k_\phi, n, \phi_0$ are parameters determined by the given force field. The second type of dihedral angle energy is a harmonic potential:

$$V^{\text{harm}}_{\text{dihedral angle}}(\mathbf{x}_i, \mathbf{x}_j, \mathbf{x}_k, \mathbf{x}_l) = \frac{k_\phi}{2}(\phi_{ijkl} - \phi_0)^2 \tag{63}$$

where $k_\phi, \phi_0$ are force-field-dependent parameters. Finally, the total dihedral angle energy $V_{\text{Dih.}}$ is defined as the sum of all applicable proper and improper dihedral-angle energy terms.

4. Lennard-Jones Energy

The Lennard-Jones energy between two atoms located at $\mathbf{x}_i$ and $\mathbf{x}_j$ is calculated as follows:

$$V_{\text{LJ}}(\mathbf{x}_i, \mathbf{x}_j) = 4\epsilon_{ij}\left(\left(\frac{\sigma_{ij}}{d_{ij}}\right)^{12} - \left(\frac{\sigma_{ij}}{d_{ij}}\right)^{6}\right), \quad d_{ij} = \|\mathbf{x}_i - \mathbf{x}_j\| \tag{64}$$

where $\sigma_{ij}, \epsilon_{ij}$ are force-field-dependent parameters that depend on the types of atoms participating in the interaction. Certain pairs of atoms are treated differently in a manner consistent with the force field used and the topology of the given system. For example, pairs of atoms that are separated by one (1-2) or two bonds (1-3) are excluded as an LJ pair.

5. Coulomb Energy

Similar to the Lennard-Jones energy, the Coulomb energy for a pair of atoms positioned at $\mathbf{x}_i$ and $\mathbf{x}_j$ is calculated as follows:

$$V_{\mathrm{Coul}}(\mathbf{x}_i, \mathbf{x}_j) = \frac{q_i q_j}{4\pi\epsilon_0 d_{ij}}, \quad d_{ij} = \|\mathbf{x}_i - \mathbf{x}_j\| \tag{65}$$

where $q_i$ and $q_j$ are the partial charges of atoms $i$ and $j$, respectively, and $\epsilon_0$ is the vacuum permittivity.

6. Correction-Map (CMAP) Energy

A CMAP is typically defined for CHARMM force fields and is used to bias the distribution of backbone dihedral angles. A CMAP defines a 2D energy surface for a set of Ramachandran backbone dihedral angles of a given residue type. More precisely, for a given residue at position $r$ in the protein sequence, the CMAP energy associated with this residue depends on the two backbone dihedral angles formed by the quintet of backbone atoms $(C^{r-1}, N^r, C_\alpha^r, C^r, N^{r+1})$. Let $\mathbf{x}_i, \mathbf{x}_j, \mathbf{x}_k, \mathbf{x}_l, \mathbf{x}_m$ denote the positions of this given quintet of backbone atoms indexed as $(i, j, k, l, m)$. The CMAP energy associated with this residue depends on the dihedral angles $\phi_{ijkl}$ and $\phi_{jklm}$. It is calculated by interpolating the fixed 2D grid of energies defined by the force field with a bicubic polynomial:

$$V_{\mathrm{CMAP}}(\mathbf{x}_i, \mathbf{x}_j, \mathbf{x}_k, \mathbf{x}_l, \mathbf{x}_m) = I_{ijklm}(\phi_{ijkl}, \phi_{jklm}) \tag{66}$$

where $I_{ijklm}$ is the 2D grid interpolator for the specific quintet defined by atoms $i, j, k, l, m$. For the CHARMM force fields used in this work, the raw correction map is a $24 \times 24$ grid where each axis spans the range $[-180°, 180°]$, i.e. each bin has a width of $15°$ along each axis.

#### E.3 Ramachandran Potential of Mean Force

To characterize the sampling of backbone conformations of the studied systems, the 2D distributions of Ramachandran backbone dihedral angles $(\phi, \psi)$ are assessed using the Ramachandran potential of mean force (PMF). Specifically, the Ramachandran PMF $E_{\text{Rama}}$ is defined as the negative logarithm of the probability density $P(\phi, \psi)$:

$$E_{\text{Rama}}(\phi, \psi) = -\log(P(\phi, \psi)) \tag{67}$$

where log() denotes the natural logarithm and $P(\phi, \psi)$ is estimated with the 2D histogram of the backbone dihedral angles sampled in a given dataset. Bins with zero probability density ($P(\phi, \psi) = 0$) are assigned a white color in all PMF maps that are shown. The PMF of each system is shifted such that its global minimum, which is the most probable $(\phi, \psi)$ state, corresponds to $E_{\text{Rama}}(\phi, \psi) = 0$.

#### E.4 Hydrogen-bond Contact Map

In addition to the Ramachandran PMF, the structural properties of the generated ensembles are assessed with hydrogen-bond contact maps. For a protein containing $n$ residues, the hydrogen-bond contact map is defined as an $(n, n)$ matrix whose $(i, j)$ entry corresponds to the probability of residue $i$ donating at least one hydrogen to residue $j$. Probabilities are estimated as the fraction of structures in which a given residue pair forms at least one hydrogen bond.

A residue pair is considered as forming a hydrogen bond if at least one donor–hydrogen–

acceptor (D–H–A) triplet satisfies the following geometric criteria:

$$\|\vec{x}_\mathrm{D} - \vec{x}_\mathrm{H}\| < 1.2\,\text{Å}$$
$$\|\vec{x}_\mathrm{D} - \vec{x}_\mathrm{A}\| < 3.0\,\text{Å}$$
$$\arccos\left(\frac{(\vec{x}_\mathrm{A} - \vec{x}_\mathrm{H}) \cdot (\vec{x}_\mathrm{D} - \vec{x}_\mathrm{H})}{\|\vec{x}_\mathrm{A} - \vec{x}_\mathrm{H}\|\|\vec{x}_\mathrm{D} - \vec{x}_\mathrm{H}\|}\right) \geq 150^\circ \tag{68}$$

where $\vec{x}_\mathrm{D}, \vec{x}_\mathrm{H}, \vec{x}_\mathrm{A}$ denote the positions of the donor, hydrogen and acceptor atom, respectively.

#### E.5 UMAP Energy Surfaces

To further assess high-dimensional features of generated structural ensembles, energy surfaces of two-dimensional UMAP[18] embeddings are compared (Supplementary Fig. 8). Two types of properties are embedded: heavy-atom positions (aligned to principal axes, as detailed below) and dihedral angles of heavy-atom structures. For the case of dihedral angles $\theta$, $(\sin(\theta), \cos(\theta))$ are embedded to properly account for their periodicity. The energy surface $E(u_1, u_2)$ of a 2D UMAP embedding $(u_1, u_2)$ is calculated similarly to the Ramachandran PMF (Supplementary Appendix E.3):

$$E(u_1, u_2) = -\log(P(u_1, u_2)) \tag{69}$$

where log() denotes the natural logarithm and $P(u_1, u_2)$ denotes the probability density of the embedding, which is estimated from the corresponding 2D histogram. Similarly to the Ramachandran PMF analysis, bins that have zero probability density ($P(u_1, u_2) = 0$) are assigned a white color and the scale of $E(u_1, u_2)$ is shifted such that its global minimum corresponds to $E(u_1, u_2) = 0$.

UMAP models are fitted to the data using the UMAP Python package[18] with the default parameters, i.e., `n_neighbors=15`, `min_dist=0.1`. The values of the properties of each structure are concatenated into a single vector with a fixed ordering. For example, the dihedral angles of a structure are ordered consistently with the structure's topology file (.top file in GROMACS). This procedure yields a data matrix of shape $(N, d)$ where $N$ denotes the total number of structures in the dataset and $d$ is the dimension of the concatenated property vector. Finally, prior to fitting the given properties with UMAP, data matrices are standardized such that they have a mean of 0 and a standard deviation of 1 along each dimension. The output of the UMAP models is similarly standardized to facilitate consistent visualization across the systems studied.

When atom positions are embedded, the raw atom positions of each structure are first aligned to their principal axes to remove rotational and translational degrees of freedom. The principal component (PC) axes are found by diagonalizing the covariance matrix of 3D positions using principal component analysis (PCA). Specifically, given a structure with raw atom positions $\tilde{\mathbf{x}}_i$ where $i$ indexes atoms in the structure, the PC-aligned atom positions $\mathbf{x}'_i$ are calculated as follows:

$$\tilde{\mathbf{x}}_{\text{centroid}} = \frac{1}{n_{\text{atoms}}} \sum_{i=1}^{n_{\text{atoms}}} \tilde{\mathbf{x}}_i \qquad \text{(Calculate centroid of structure)} \tag{70}$$

$$\tilde{\mathbf{y}}_i = \tilde{\mathbf{x}}_i - \tilde{\mathbf{x}}_{\text{centroid}} \qquad \text{(Remove translational degrees of freedom)} \tag{71}$$

$$C = \frac{1}{n_{\text{atoms}}} \sum_{i=1}^{n_{\text{atoms}}} \tilde{\mathbf{y}}_i^T \tilde{\mathbf{y}}_i \qquad \text{(Calculate covariance matrix. } \tilde{\mathbf{y}}_i \text{ is a row vector)} \tag{72}$$

$$VSV^T = C \qquad \text{(Diagonalize } C \text{ to find principal components)} \tag{73}$$

$$V' = V \begin{pmatrix} 1 & 0 & 0 \\ 0 & 1 & 0 \\ 0 & 0 & \det(V) \end{pmatrix} \qquad \text{(Ensure } V' \text{ does not have reflections.)} \tag{74}$$

$$\mathbf{x}'_i = \tilde{\mathbf{y}}_i V' \tag{75}$$

where the last manipulation ensures that $V'$ strictly belongs to the $SO(3)$ subgroup of $O(3)$. After obtaining the PC-aligned atom coordinates $\mathbf{x}'$, there remain degrees of freedom in determining the orientation of the principal axes because the eigenvectors of $C$ are unique only up to reflections. Specifically, for any set of principal axes $[\vec{u}_x, \vec{u}_y, \vec{u}_z]$, one can define another equivalent set by reflecting any two axis vectors, e.g. $[-\vec{u}_x, -\vec{u}_y, \vec{u}_z]$. Therefore, when diagonalizing $C$, some principal axes may be flipped by chance depending on the diagonalization method. Since UMAP models are sensitive to the sign of embedded values, this degree of freedom must be removed to avoid introducing additional variance in the embedded space. To do so, the orientation of the principal axes is determined by enforcing the condition that the majority of the structure's positions are on the positive side of the first two axes. Specifically, $\mathbf{x'}_i$ is further transformed using a structure-specific matrix $R$ determined as follows:

$$\mathbf{s} = \mathrm{sign}\left(\sum_{i=1}^{n_{\mathrm{atoms}}} \mathrm{sign}(\mathbf{x'}_i)\right) \quad \text{(Find sign majority along each axis of the structure)} \tag{76}$$

$$R = \begin{pmatrix} s_x & 0 & 0 \\ 0 & s_y & 0 \\ 0 & 0 & s_x s_y \end{pmatrix} \quad \text{(Flip axes with sign, but do not introduce reflections)} \tag{77}$$

$$\mathbf{x}_i = \mathbf{x'}_i R \tag{78}$$

where it is assumed that $\mathrm{sign}(\cdot)$ is applied element-wise on a vector input and, for this specific calculation, $\mathrm{sign}(0) = 1$. The aligned atom positions $\mathbf{x}_i$ are given as input to the UMAP model.

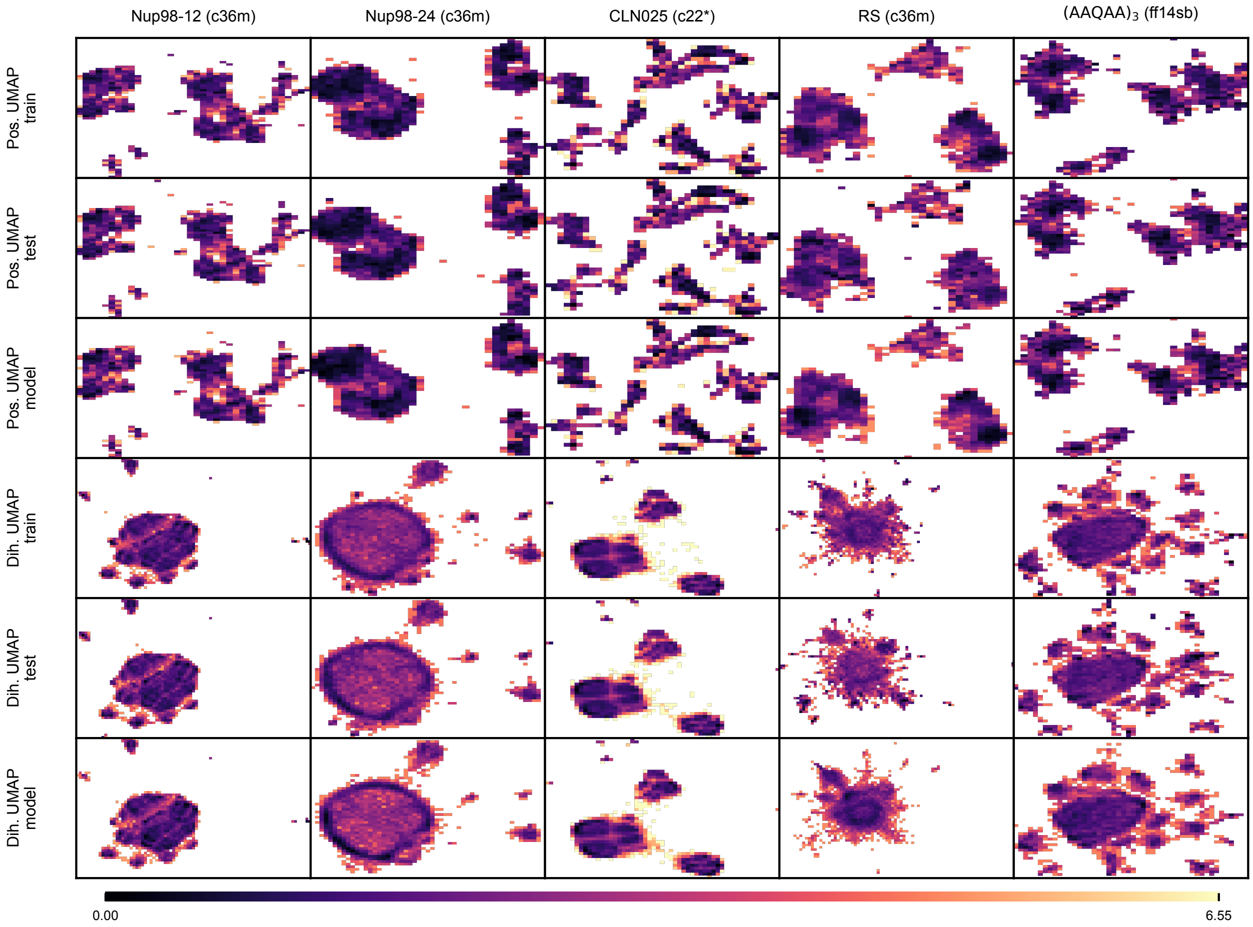

Supplementary Figure 8: **UMAP Energy Surfaces**
2D energy surfaces of UMAP-embedded properties are compared for each studied system. Energies are calculated as $-\log(P(u_1, u_2))$, where the probability density $P(u_1, u_2)$ is calculated from histograms of the UMAP vectors. For MD datasets, probability densities are averaged over trajectories contained in the dataset. In addition, for each embedded property of each system, the color scale is shifted such that zero corresponds to the global energy minimum across all surfaces (train, test and model).

#### E.6 Assessment of Model Overfitting

Although several measures were taken to reduce the chance of overfitting, such as reducing the number of bias parameters, tracking validation loss and using a small batch size, some overfitting on the training dataset may still occur. To quantify overfitting to the training dataset, the similarity of structures of a given ensemble was compared to structures in the training dataset using the root mean squared deviation (RMSD). Specifically, for a given structure $\mathbf{y}$, its minimum train RMSD is calculated as follows:

$$\mathrm{RMSD}_{\text{train, min}}(\mathbf{y}) = \min_{j \in J_{\text{train}}} \min_{\substack{R \in SO(3) \\ t \in \mathbb{R}^3}} \sqrt{\frac{1}{n_{\text{atoms}}} \sum_{i=1}^{n_{\text{atoms}}} \|R\mathbf{y}_i + t - \mathbf{x}_i^j\|^2} \tag{79}$$

where $i$ indexes atoms in the structure, $j$ indexes structures in the training index set $J_{\text{train}}$ and the inner minimum indicates that the given structure is rotated and translated to best align it with the training structure before the RMSD is calculated. These transformations are found using the Kabsch algorithm.[43]

Distributions of $\mathrm{RMSD}_{\text{train, min}}$ for model datasets and their respective test sets are compared in Supplementary Fig. 9. In this comparison, a leftward shift (smaller $\mathrm{RMSD}_{\text{train, min}}$) of the model distributions compared to the test distributions would indicate overfitting. As shown, the model datasets' $\mathrm{RMSD}_{\text{train, min}}$ are distributed similarly to the corresponding test sets. Based on these comparisons, the models did not overfit onto their training datasets.

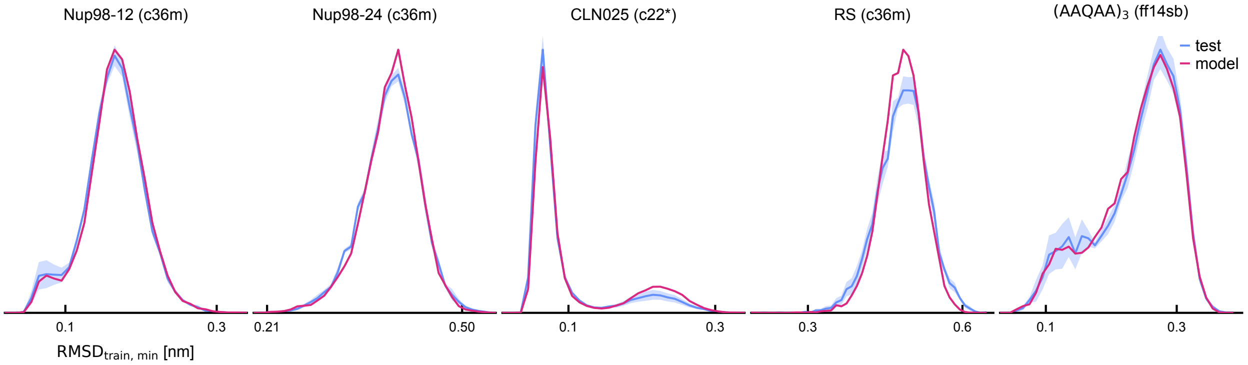

Supplementary Figure 9: **Overfitting assessment**
Distributions of the $\mathrm{RMSD}_{\mathrm{train,\ min}}$ are compared for all studied systems. The $\mathrm{RMSD}_{\mathrm{train,\ min}}$ of a given structure is the minimum RMSD found after comparing a given structure to all structures in the respective training set (refer to eq. (79)). Shaded regions of the test set distributions indicate the mean ± the standard error of the mean over trajectories.

#### E.7 Effect of Cross-Product Module on Dihedral Angles

To assess the value of cross-product modules for sampling accurate structural ensembles, these modules were ablated from the coarse-structure and fine-structure models and the resulting models were retrained on the Nup98-12 system. The sampling methodology remained the same as that of the base model. To compensate for the loss of expressibility, the dimensions of the latent node feature space and edge feature space were increased from 64 to 75. This changed the total number of parameters of each type of model to $\sim$ 800k, which is comparable to the number of parameters for models with cross-product modules ($\sim$ 750k).

The dihedral angles of structures sampled with models lacking a cross-product module exhibit a spurious chiral symmetry that is not exhibited in the MD ensemble or the base model's ensemble (Supplementary Fig. 10). Due to the lack of chirality-aware modules, the model lacking the cross-product module does not modify the chirality of these features during denoising. Therefore, the chirality is set by the initial structure, which has a 50% chance of sampling each chirality since it originates from pure noise.

#### E.8 Effect of Force Field on Ensemble Learning

To assess the effect of the force field on the model's ability to learn the conformational ensemble of a given protein, models were trained on trajectories generated with two different force fields: CHARMM36m and Amber14sb. In order to carry out simulations with the Amber14sb force field, it was necessary to set up the system with neutral termini, with acetyl and N-methylamide capping groups for the N- and C-termini, respectively. The system used for this test, denoted Nup98-12-capped, has the same sequence as the Nup98-12 system, but its topology differs due to the addition of N/C terminal caps. To avoid introducing confounding factors in assessing the effect of force field on ensemble learning, a set of Nup98-12-capped trajectories was also generated with CHARMM36m. Apart from using capped termini and using the Amber14sb force field for one of the datasets, both Nup98-12-capped datasets were generated, treated and analyzed in the same manner as the main Nup98-

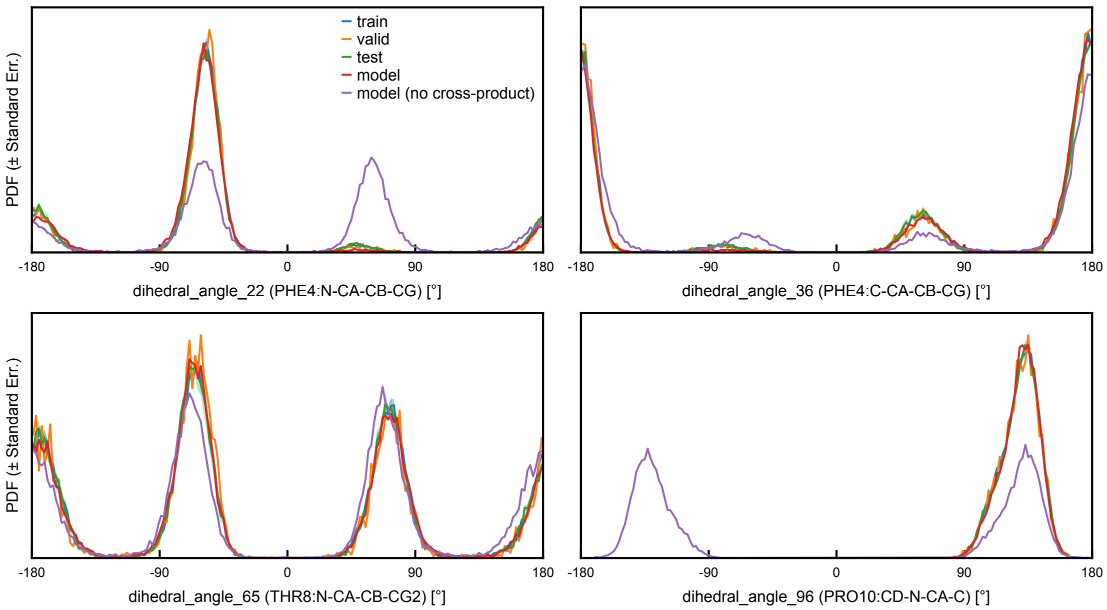

Supplementary Figure 10: **Effect of Cross-Product Modules**
Four examples of dihedral angle distributions are compared for the Nup98-12 system. The train, validation and test datasets for this system are the same as those presented in the main text Figures 3 to 5. Two model datasets are compared where one (red) is generated with the base model, which uses the cross-product module, while the other (purple) is generated with a model that does not have a cross-product module. The label below each plot identifies the residue name, residue index and atom names of the quartet of atoms that form the given dihedral angle; the angle number and atom names are those used in the GROMACS topology file of the system.

12 dataset (see Supplementary Appendix D for details). With these two Nup98-12-capped datasets, and their respective model ensembles, an ensemble comparison identical to that presented for the five main systems (Figures 3 and 4) was performed.

In the fine structure assessment (Supplementary Fig. 11a), the model achieves similar accuracy for both force fields on the bond length and bond angle metrics, indicating that learning of the smallest length scales is unaffected by the underlying force field. Comparison of the dihedral angle energy distributions, however, indicates that the model distribution is slightly right-shifted relative to the MD ensemble for the Amber14sb force field, while the model and MD ensembles are consistent for the CHARMM36m force field. The CHARMM family of force fields applies a grid-based energy correction (CMAP) to the backbone dihedral angles, which we hypothesized might make the Ramachandran map more difficult for the model to learn. The Amber14sb maps are smoother than those of CHARMM36m for this particular system (Supplementary Fig. 11a, bottom three rows); notably, this greater smoothness does not translate into easier learning, as the quantitative comparison below shows.

To test our hypothesis about the learning difficulty, we compare the Ramachandran maps quantitatively using the Jensen–Shannon (JS) distance. The JS distance between the Ramachandran maps of the model and test ensembles is smaller for the CHARMM36m ensembles (0.34) than for the Amber14sb ensembles (0.37). These JS distances are similar to the train-test JS distances for both force fields (0.32 and 0.35 for CHARMM36m and Amber14sb, respectively). Together, these comparisons suggest that the force field does not measurably affect the model's ability to learn the Ramachandran map, contrary to our hypothesis that the CMAP of CHARMM force fields increases the learning difficulty.

While the choice of force field does not measurably affect how accurately the model reproduces the Ramachandran map, it strongly shapes the map itself. Force-field choice is known to introduce noticeable biases when sampling ensembles of IDPs.[24] Comparing maps across force fields, the JS distance between the CHARMM36m and Amber14sb maps is 0.47 for the test sets and 0.45 for the training sets, which are larger than either of the model-test JS distances reported above (0.34 and 0.37). This indicates that changing the underlying force field results in a larger change in the Ramachandran map than replacing MD sampling with sampling by the model. In other words, although the model samples more broadly than MD based on comparisons of the Ramachandran maps (Supplementary Fig. 11a and Figure 3), it retains bias from the force field.

For the coarse structure assessment (Supplementary Fig. 11b), the model energy distributions are generally more right-shifted relative to the MD test set distributions when trained on the Amber14sb ensemble compared to the CHARMM36m ensemble. Comparing the sampling of non-bonded interactions reported by the Lennard-Jones and Coulomb energy distributions, the model exhibits a larger right shift relative to the MD distributions when trained on Amber14sb structures. The radius of gyration is the only metric that shows agreement between the model and test set for the Amber14sb force field. Comparison of the total energy distributions also demonstrates that the model more accurately recapitulates MD sampling when trained on the CHARMM36m force field. Consistent with what was observed for the hydrogen-bond contact maps for the other systems (Figure 4), the model ensembles exhibit lower probabilities of hydrogen-bonded contacts in general compared to the corresponding MD ensembles, irrespective of force field.

In summary, the fine structure assessment shows that the force field has only minor effects on the fine structure learning difficulty. The coarse structure assessment, however, indicates that the model has greater difficulty in learning the Nup98-12-capped ensemble generated with Amber14sb than the one generated with CHARMM36m.

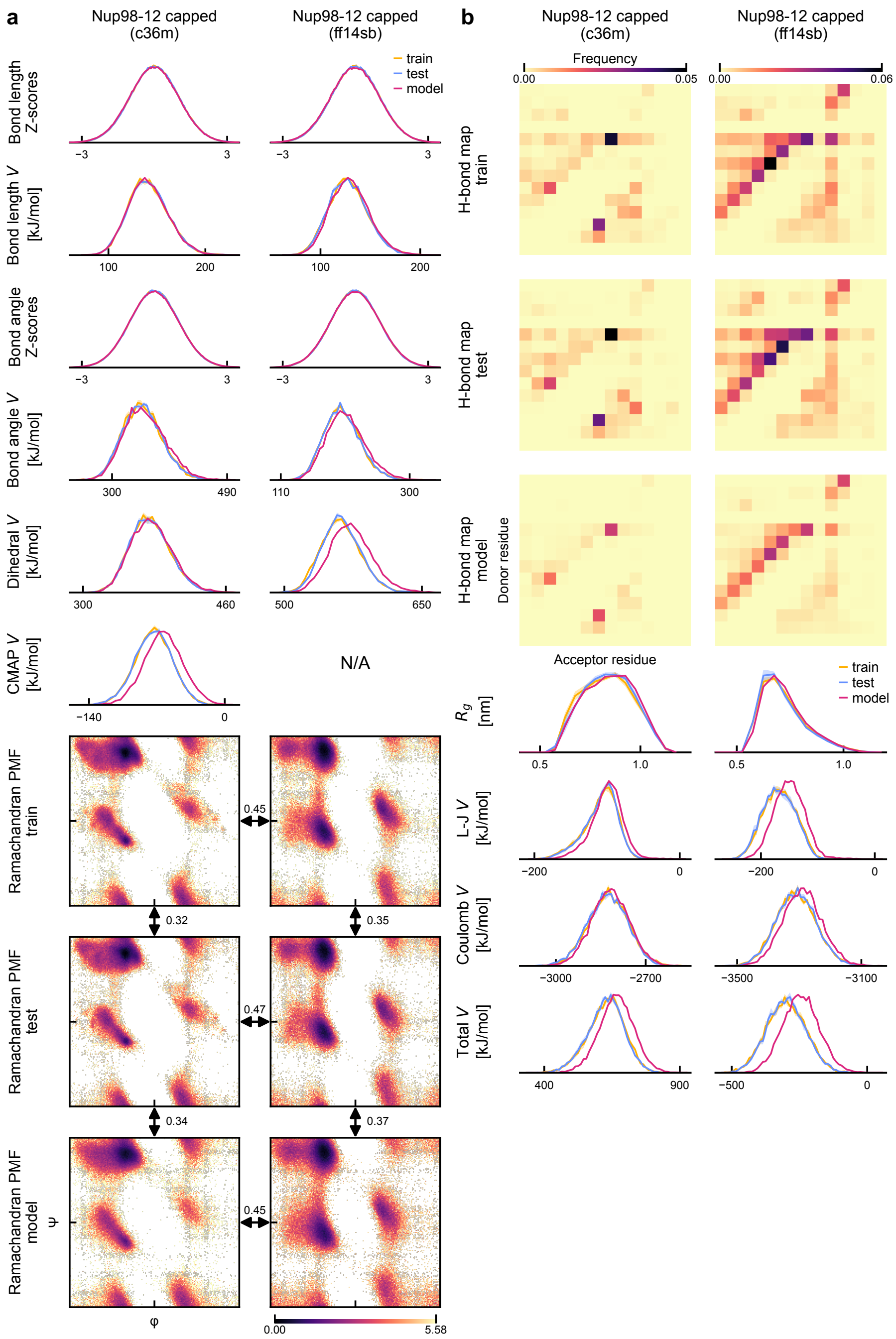

a
Nup98-12 capped (c36m)
Nup98-12 capped (ff14sb)
train
test
model
Bond length Z-scores
Bond length V [kJ/mol]
Bond angle Z-scores
Bond angle V [kJ/mol]
Dihedral V [kJ/mol]
CMAP V [kJ/mol]
N/A
Ramachandran PMF train
Ramachandran PMF test
Ramachandran PMF model
0.45
0.32
0.35
0.47
0.34
0.37
0.45
ψ
φ
0.00
5.58
b
Nup98-12 capped (c36m)
Nup98-12 capped (ff14sb)
Frequency
0.00
0.05
0.06
H-bond map train
H-bond map test
H-bond map model
Donor residue
Acceptor residue
$R_g$ [nm]
L-J V [kJ/mol]
Coulomb V [kJ/mol]
Total V [kJ/mol]

Supplementary Figure 11: **Force Field Effects.** Fine-structure (**a**) and coarse-structure (**b**) metrics are compared for the same Nup98-12-capped system generated with two force fields: CHARMM36m and Amber14sb. (**a**) Probability densities of several metrics (rows), including Z-scores and potential energies, are shown for both force fields (columns). Supplementary Appendix E details how these metrics were computed. The distributions for the model, train and test sets are shown in each case. The train and test distributions are averaged over independent MD trajectories, with the shaded area indicating the standard error of the mean. Note that Amber14sb has no CMAP correction. The last three rows show the Ramachandran potential of mean force (PMF), $E_{\mathrm{Rama}}(\phi, \psi) = -\log P(\phi, \psi)$, shifted such that the global minimum is zero for the model, train and test sets. Backbone dihedral angles $\phi, \psi$ span the range $[-180°, 180°]$. Pairwise comparisons between Ramachandran maps are provided, with labels indicating JS distance values. (**b**) Hydrogen-bond contact maps are shown for the model, train and test sets of both force fields (upper three rows). The color of each square in these maps indicates the probability of a hydrogen bond between a pair of residues. In the lower four rows, one-dimensional distributions compare model, train and test set distributions as indicated in the legend. Distributions for the train and test sets are averaged over independent MD trajectories, and the standard error of the mean is shown (shaded area).

#### E.9 Effect of MD Equilibration of Model Datasets

Energy minimization has been used in conjunction with generative models of protein structures to relax high-energy interactions within the structure.[44] Here, we test the effect of MD equilibration on model datasets to determine the extent to which this would reduce bias in model ensembles. These tests also serve as an evaluation of the proximity (in terms of equilibration time) of the model ensemble to the MD ensemble.

Structures from the model ensembles are equilibrated following a procedure similar to that used in the construction of the MD datasets (Supplementary Appendix D). First, each generated structure is solvated in a box of water whose dimensions match the dimensions of the original MD simulation box. Then, water molecules that are within 1.5 Å of any of the protein's atoms are removed to prevent atom clashes. Finally, additional water molecules are removed and $Na^+$ and $Cl^-$ ions are added to exactly match the number of water molecules and ions in the original MD box. Subsequently, the system's energy is minimized until the maximum force is smaller than 1000.0 kJ/mol/nm or 50,000 optimization steps are performed. Following these preprocessing steps, 100 ps of molecular dynamics in the NVT ensemble is run with a thermostat that uses stochastic velocity-rescaling[40] and constraints on bonds including hydrogen atoms are enforced with the LINCS algorithm.[38] The temperature of the NVT ensemble is set to match the temperature of the original simulation, i.e. 340 K for the CLN025 system and 298 K for all other systems.

Once all equilibration runs terminate, the equilibrated dataset is built by extracting the final heavy-atom structures of these runs. Hydrogen atoms are then added using GROMACS' `pdb2gmx`, consistent with the method used for the MD and model datasets. An equilibrated dataset is constructed for each of the five systems studied in this work. For the case of CLN025, we created the simulation box of the underlying simulation using the information available in Lindorff-Larsen et al.[22] Details of the simulation box were not provided in that work, so this is an approximation.

The fine and coarse structure of the equilibrated datasets is assessed in a manner similar to that presented in the main text (Supplementary Figs. 12 and 13). Compared with the original model datasets, the equilibrated datasets generally exhibit less variability and reduced discrepancy with the MD potential energy distributions. For the Nup98-12 and $\texttt{(AAQAA)}_3$ systems, the 100 ps equilibration is sufficient to recover the features of the MD ensemble. However, for the Nup98-24 and RS peptide systems, equilibration helps reduce rightward biases in the potential energies, but not completely; these longer peptides (24 residues) may require longer equilibration compared to the Nup98-12 and $\texttt{(AAQAA)}_3$ systems (12 and 15 residues, respectively).

### F Sampling Efficiency

#### F.1 Measurement of Ensemble Generation Time

To assess the efficiency of our proposed method for constructing a given structural ensemble, the time required to complete each of the three main phases of the method for each studied system was measured. In all cases, measurements were performed on the same workstation equipped with 8 CPU cores and 1 GPU (Supplementary Appendix B.7).

First, the time to generate MD trajectories was estimated using GROMACS' `mdrun`. The timed runs consisted of a 1 ns trajectory using the same MD simulation parameters as the production runs that were harvested to build the datasets (Supplementary Appendix D). To use the GPU in MD runs, GROMACS was compiled with GPU-aware MPI, and particle-particle and particle-mesh Ewald tasks were performed on the GPU. A summary of the sampling performance of MD is given in the first column of Supplementary Table 9. For further details on the GROMACS configuration, we refer the reader to the benchmarking log files provided in the data repository. Note that initialization steps, including system preparation, energy minimization and equilibration, are not accounted for in the MD performance estimates. As these are one-time setup steps per system, their compute cost is amortized, becoming negligible for the long sampling times of interest here.

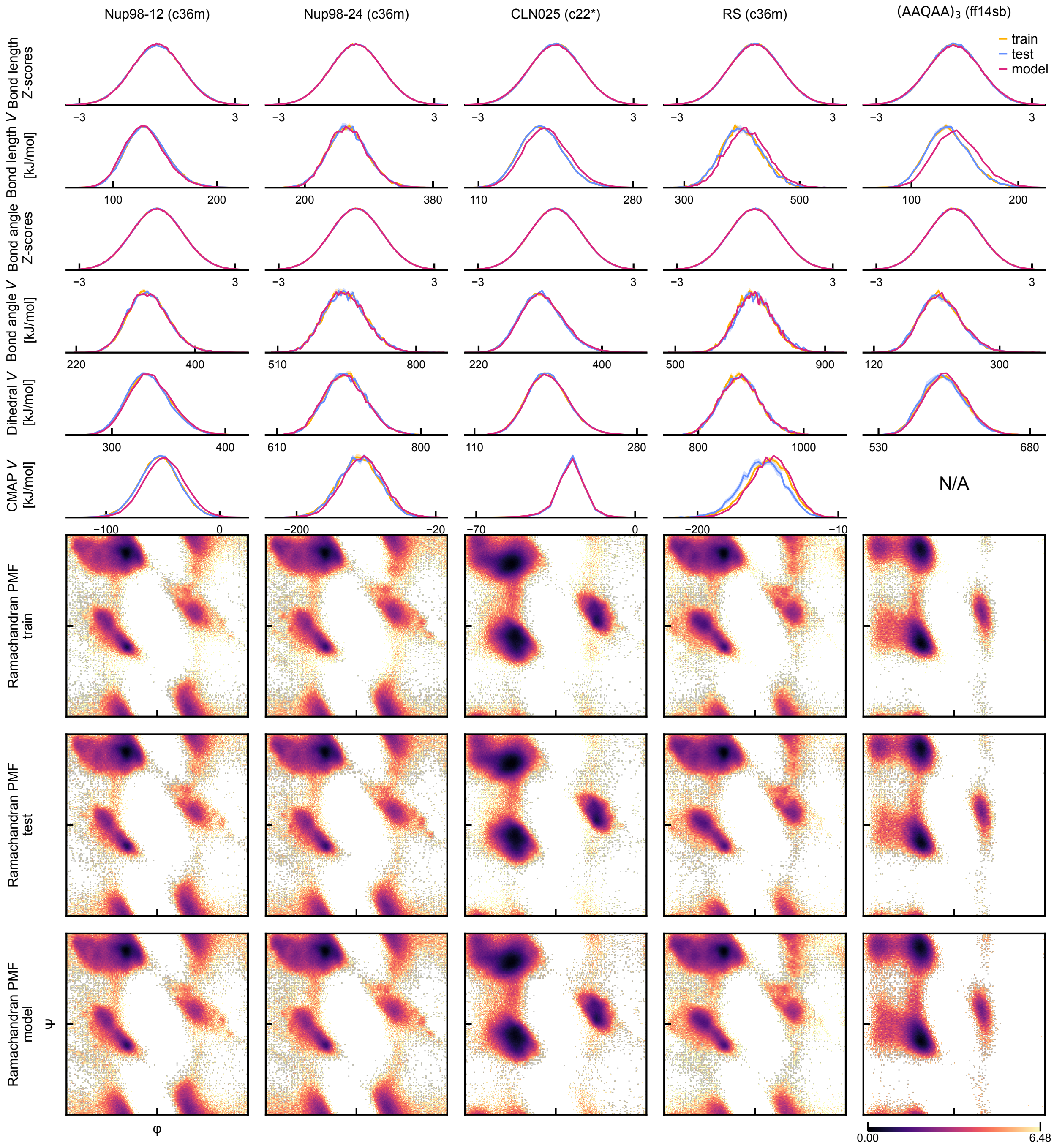

Supplementary Figure 12: **Fine Structure Assessment of Equilibrated Model Datasets.**
Probability densities of several metrics (rows), including Z-scores and potential energies, are shown for all studied systems (columns). Supplementary Appendix E details how these metrics were computed. The distributions for the model, train and test sets are shown in each case. Distributions for the train and test sets are averaged over independent MD trajectories, and the standard error of the mean is shown (shaded area). Note that Amber14sb has no CMAP correction. The last three rows show the Ramachandran potential of mean force (PMF), $E_{\text{Rama}}(\phi, \psi) = -\log P(\phi, \psi)$, shifted such that the global minimum is zero for the model, train and test sets. Backbone dihedral angles $\phi, \psi$ span the range $[-180°, 180°]$.

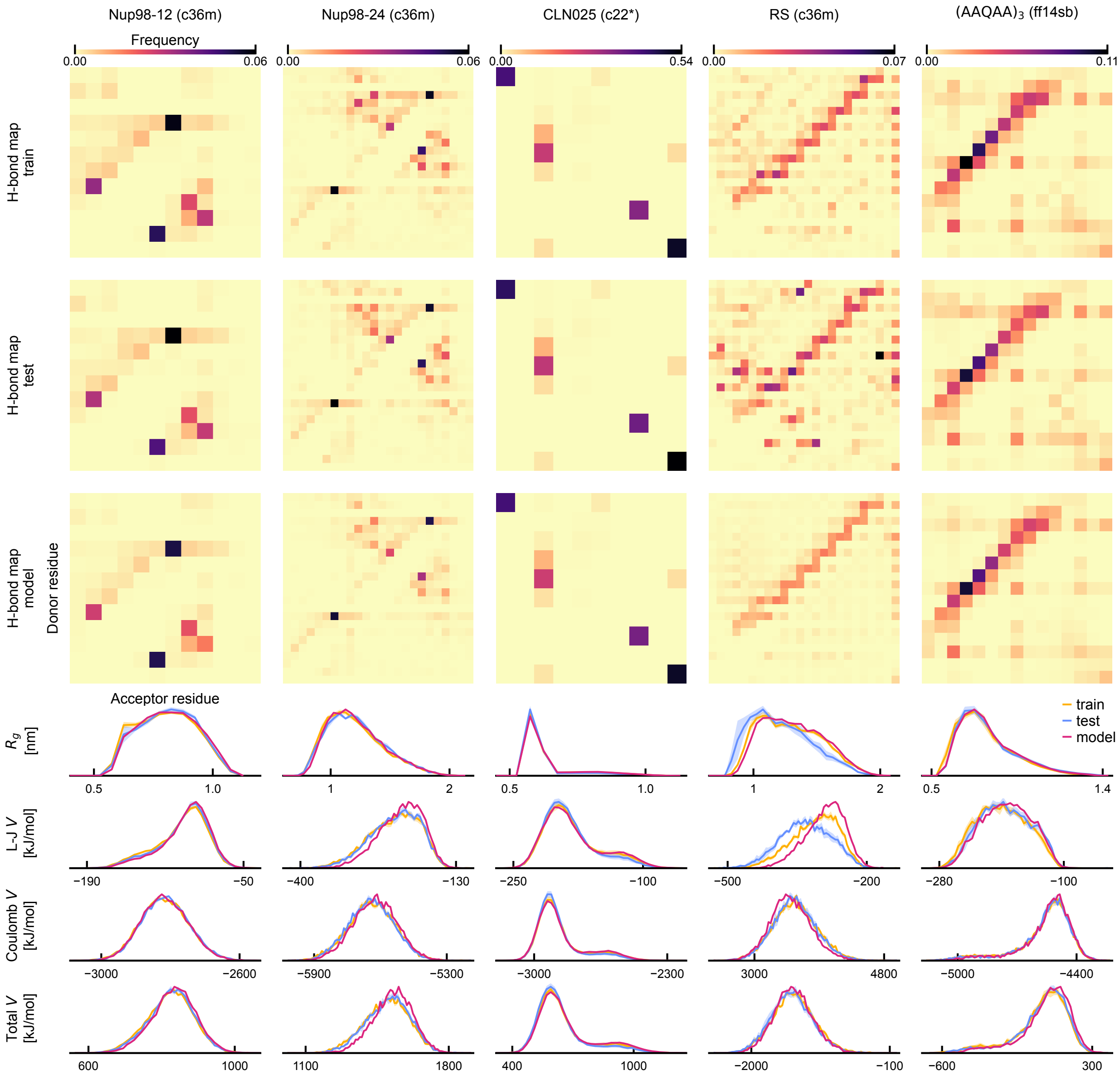

Supplementary Figure 13: **Coarse Structure Assessment of Equilibrated Model Datasets**
Hydrogen-bond contact maps are shown for the model, train and test sets of all systems (upper three rows). The color of each square in these maps indicates the probability of a hydrogen bond between a pair of residues. In the lower four rows, one-dimensional distributions compare model, training and test set distributions as indicated in the legend. Distributions for the train and test sets are averaged over independent MD trajectories, and the standard error of the mean is shown (shaded area).

Second, the time to train the coarse-structure and fine-structure models was estimated for each given system (Supplementary Table 9, columns 2 and 3). The estimated training time accounts for all steps performed in a single training epoch, including train and validation loss evaluation, backward propagation, optimizer steps, generation of the 100-sample monitoring dataset and calculation and recording of monitored values (losses and metrics). Note that the estimated time does not account for data loading since all model input data was loaded onto the GPU prior to starting parameter optimization. This was possible because the size of the studied systems was sufficiently small. Generally, this is not possible, and loading data from the hard drive to the GPU memory takes a non-negligible amount of time. However, this additional compute time can usually be amortized with parallelized data loaders. Lastly, the initial steps of model training, such as model initialization and input feature calculation, were not taken into account. These steps generally took less than one minute to complete and were performed once.

Finally, the time to generate the coarse and fine structures of 10,000 samples was estimated (Supplementary Table 9, column 4). The sampling time accounted for the calculation of the input model features, loading of the model input data onto the GPU, calculation of the denoising step with the trained model, and calculation of the conditioners' gradient. The majority of the sampling time was spent denoising the heavy-atom structure. Although the training time and sampling time generally increase with system size, there are some variations. For example, although the number of atoms in CLN025 is smaller than $\texttt{(AAQAA)}_3$, the compactness of the CLN025 structures increases the computational complexity of the message-passing layers (Supplementary Appendix A.2), which increases both training and sampling time.

#### F.2 Assessment of Model Ensemble Convergence

To assess the convergence rate of the model in constructing a given conformational ensemble, the difference between a given ensemble and the associated test ensemble is quantified as a

Supplementary Table 9: **Compute Times of Sampling Methodology**
The sampling rate of MD is compared against the training time of the coarse-structure (CG) and fine-structure (HA) models and the sampling of 10,000 structures with the models. The second MD column is calculated from the first column assuming samples are collected every 1 ns of simulated time. Reported times are estimated based on one MD trajectory or one instance of training and sampling. The last column provides the sampling rate of the model (col. 5) in units of the MD sampling rate (col. 2).

| | MD (ns/day) | MD (1 ns inter.) (h/10k samp.) | Training (h) | | Sampling (h/10k samp.) | Sampling Acceleration |
|---|---|---|---|---|---|---|
| | | | CG | HA | | |
| Nup98-12 | 1962.368 | 122.301 | 2.202 | 3.119 | 0.978 | 125 |
| Nup98-24 | 614.082 | 390.827 | 3.550 | 4.410 | 2.564 | 152 |
| CLN025 | 2117.090 | 113.363 | 3.481 | 8.293 | 1.735 | 65 |
| RS peptide | 1062.156 | 225.956 | 3.533 | 7.337 | 3.587 | 63 |
| `(AAQAA)`$_3$ | 1125.517 | 213.235 | 1.752 | 2.793 | 1.291 | 165 |

function of the number of samples $N$ in the given ensemble. As $N$ increases, the difference is expected to decrease, and the rate of decrease quantifies the value of additional structures in converging to the test ensemble of the given system. Two types of ensembles are compared to the test ensemble: train and model ensembles.

The difference between a given ensemble and the test ensemble is quantified by comparing the distributions of several geometric features that assess the coarse and fine structures of the system. The fine structure is assessed using the bond length and bond angle distributions while the coarse structure is assessed using the Ramachandran PMF and the radius of gyration. For a given feature, the difference between the two respective distributions is quantified with the JS distance (JSD). Note that, in this work, JSD refers to the Jensen-Shannon *distance*, and not the Jensen-Shannon divergence. More precisely, given two distributions $P(\vec{m}), Q(\vec{m})$ of a given metric $\vec{m}$ with dimension $d$, the JSD between them is calculated as follows:

$$JSD(\vec{m}) = \sqrt{\frac{KL(Q||M) + KL(P||M)}{2}}, \quad M = \frac{P+Q}{2} \tag{80}$$

where $KL(P||Q)$ denotes the Kullback-Leibler (KL) divergence of a probability density $Q$ relative to a reference probability density $P$ estimated using histograms of the metric $\vec{m}$.

For metrics with a large number of dimensions (such as bond lengths and bond angles), estimating $JSD$ with $d$-dimensional histograms is inaccurate due to the prevalence of low-count bins caused by the curse of dimensionality. Instead, when $d > 2$, the average $JSD$ across all 1-dimensional marginal distributions is used as an estimate of $JSD$:

$$\text{For } d > 2, \quad JSD(\vec{m}) = \frac{1}{d}\sum_{i=1}^{d} JSD(\vec{m}_i) \tag{81}$$

where $JSD(\vec{m}_i)$ denotes the JSD of the $i^{\text{th}}$ component of $\vec{m}$.

To remove the arbitrariness of selecting the samples that constitute a given model subset of size $N$, 100 random subsets of the model dataset are constructed by sampling without replacement. For each value of subset size $N$, the 100 values of $JSD$ are used to calculate the average and standard error of $JSD$. Finally, these statistics are used to construct the $JSD(N)$ curves.

To calculate the statistics of $JSD(N)$ for the training datasets, structures are not randomly selected across the entire dataset, as this would break the temporal ordering of MD. Instead, to preserve the ordering inherent to the MD sampling process, random sampling is performed at the level of trajectories. Specifically, for a given system, the trajectories in the training datasets are concatenated in a random order along the time dimension to form one trajectory. Then, the $JSD$ of the set of all structures that appear before a given time $t$ in this concatenated trajectory is calculated for some predetermined values of $t$. Since the subset size $N$ increases monotonically with $t$, higher values of $t$ correspond to larger subsets. This procedure is meant to reproduce the ensemble building process of MD whereby additional samples are generated as the simulation progresses. Finally, similar to the treatment of model datasets, the bootstrap estimate of the average and standard error of $JSD(N)$ is calculated across 100 random orderings of the trajectories.

#### F.3 Assessment of Models Trained on Minimal Datasets

The ability of the model to learn a given conformational ensemble is further assessed by retraining on a minimal amount of data. This test is meant to push the model to its limit and assess its learning ability in scenarios where training data is scarce. With the new models trained on minimal datasets, the sampling and evaluation methodology is repeated to assess the fine and coarse structure of the model datasets.

Details of the minimal training datasets are given in the last column of Supplementary Table 7. These datasets contain at most 10% of the total number of structures in a given ensemble, which is at most 4000 structures for all systems except CLN025. Given the limited structural variability in the minimal datasets, all models overfitted their training data in the last stage of training, as evidenced by increasing validation loss curves (Supplementary Fig. 14). To be consistent with the training strategy used for the main datasets, the state of the fine-structure and coarse-structure models' parameters halfway through training (epoch $\approx$ 2000) was used to generate ensembles. This is also close to the onset of overfitting for some systems (Nup98-24, RS peptide and $\texttt{(AAQAA)}_3$).

The fine structure assessment of the minimally trained models is presented in Supplementary Fig. 15. In general, bond lengths and bond angles are well-sampled by the model, but the bond energies of the Nup98-24, RS peptide and $\texttt{(AAQAA)}_3$ systems show evidence that the model is sampling bond lengths closer to the equilibrium bond lengths defined by the force field energy function. The dihedral angle energy is also well-sampled, except for the CLN025 system where the model distribution is slightly right-shifted. For backbone dihedral angles, model distributions are generally more dispersed compared to MD, which was also observed in the Ramachandran maps in the assessment of the main datasets (Figure 3).

The coarse structure assessment of the minimally trained models is presented in Supplementary Fig. 16. Similar to the main assessment (Figure 4), the global structure of hydrogen-bond contact maps is preserved by the model, but frequencies are underestimated relative to the MD maps. The radius of gyration is accurately sampled for the Nup98-24,

CLN025 and (AAQAA)$_3$ systems, but the model exhibits a bias towards higher values for the Nup98-12 and RS peptide systems. In all systems, the Lennard-Jones energy distribution of the model dataset is right-shifted compared to the MD distribution. The Coulomb energies are accurately sampled for the Nup98-12, CLN025 and RS peptide systems, but discrepancies are observed for the Nup98-24 and (AAQAA)$_3$ systems. Finally, the model distributions of coarse structure metrics are sometimes right-shifted compared to the corresponding MD distributions. In summary, the model retains the essential fine and coarse structure features when trained on less than 10% of the original datasets, with accuracy and variability comparable to models trained on the complete datasets, though with additional discrepancies as noted above.

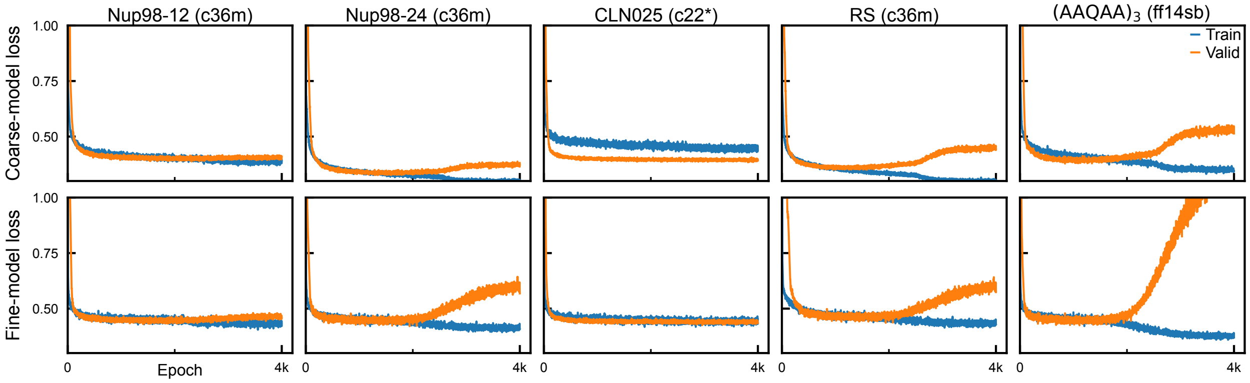

Supplementary Figure 14: **Training Losses on Minimal Datasets.**
Training and validation losses for models trained on the minimal datasets. Both the coarse-structure and fine-structure models' losses are shown.

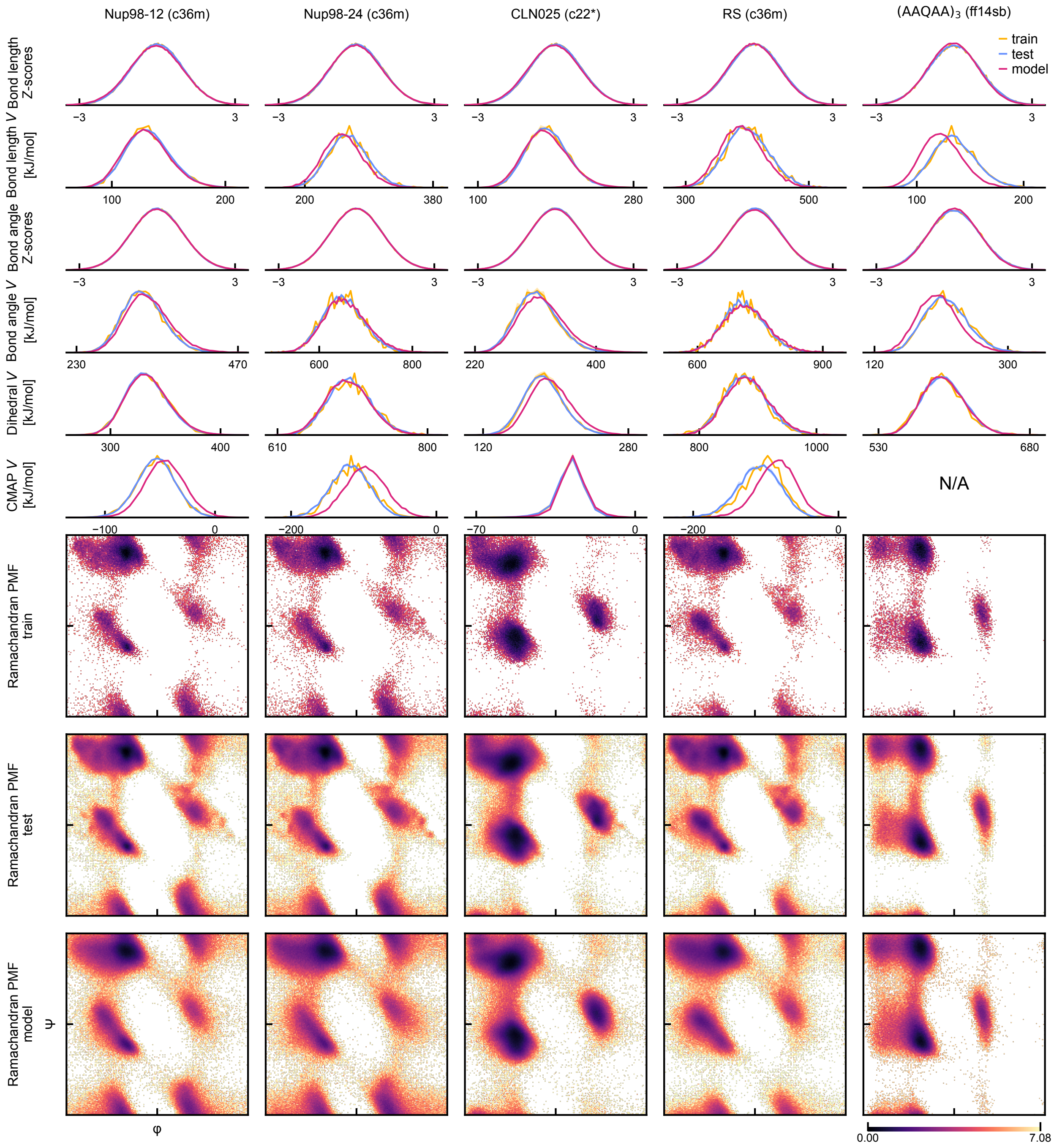

Supplementary Figure 15: **Fine Structure Assessment of Models Trained On Minimal Datasets**

Probability densities of several metrics (rows), including Z-scores and potential energies, are shown for all studied systems (columns). Supplementary Appendix E details how these metrics were computed. The distributions for the model, train and test sets are shown in each case. The train and test distributions are averaged over independent MD trajectories, with the shaded area indicating the standard error of the mean. Note that Amber14sb has no CMAP correction. The last three rows show the Ramachandran potential of mean force (PMF), $E_{\mathrm{Rama}}(\phi, \psi) = -\log P(\phi, \psi)$, shifted such that the global minimum is zero for the model, train and test sets. Backbone dihedral angles $\phi, \psi$ span the range $[-180°, 180°]$.

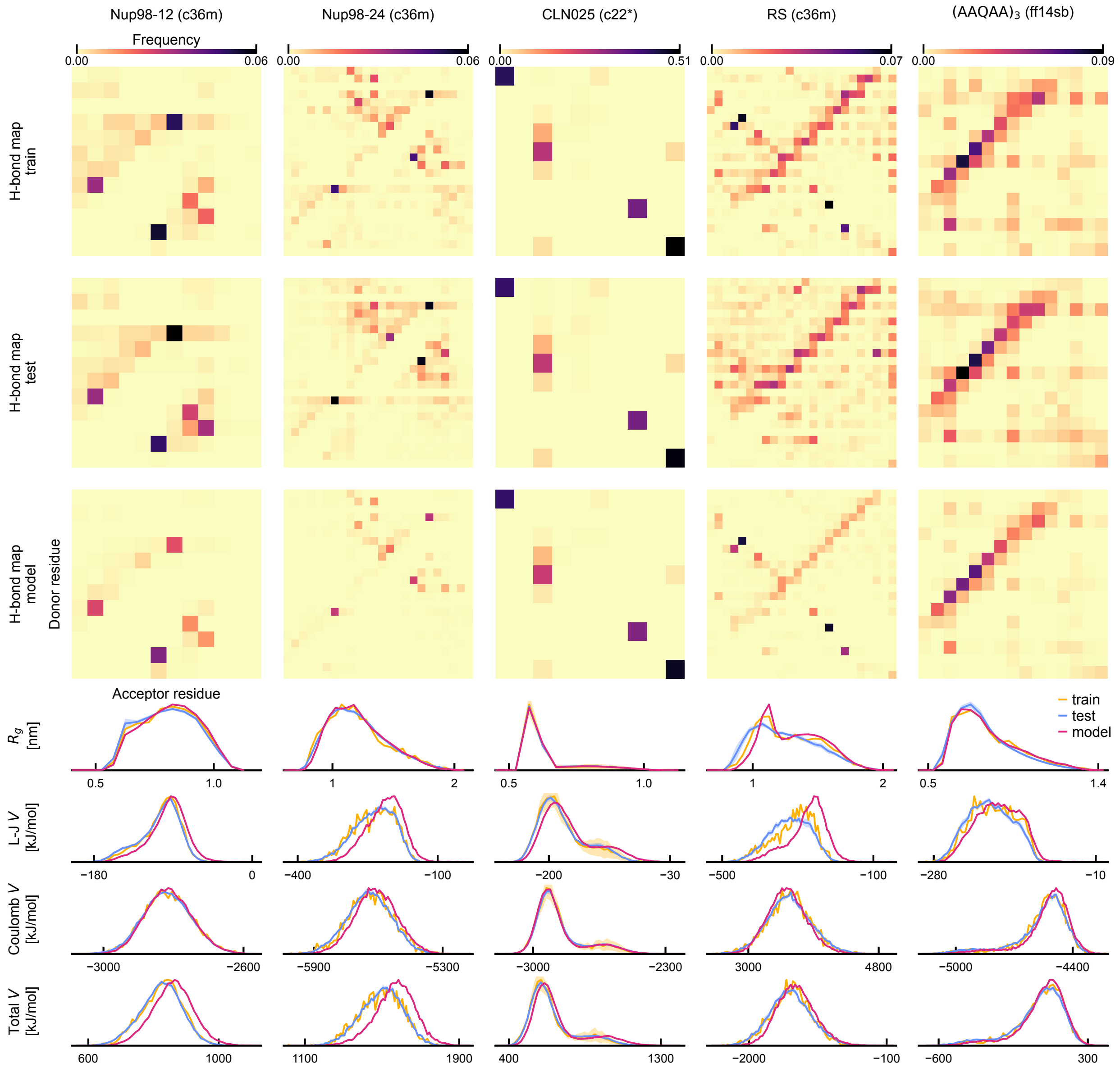

Supplementary Figure 16: **Coarse Structure Assessment of Models Trained On Minimal Datasets**
Hydrogen-bond contact maps are shown for the model, train and test sets of all systems (upper three rows). The color of each square in these maps indicates the probability of a hydrogen bond between a pair of residues. In the lower four rows, one-dimensional distributions compare model, training and test set distributions as indicated in the legend. Distributions for the train and test sets are averaged over independent MD trajectories, and the standard error of the mean is shown (shaded area).

#### F.4 Discovery Rate of Backbone Dihedral States

As a measure of sampling efficiency, the rate of discovering unique conformational states is compared between the model and MD datasets. For this analysis, conformational states are defined using backbone dihedral angles. Specifically, the backbone dihedral state of a given residue is an integer value that identifies a predefined Ramachandran map region where its pair of backbone dihedral angles $(\phi, \psi)$ is located. In this analysis, the Ramachandran map is divided into four regions (Supplementary Fig. 17). These boundaries ensure that secondary structure basins are not split into separate states. For example, the periodic continuity of the $\beta$-sheet basin across the $\psi$ periodic boundary ($\psi = \pm 180°$) is preserved. We then define the backbone dihedral state of a protein with $N_r$ residues as the ordered sequence of the $N_r - 2$ per-residue states. The two terminal residues, whose $\phi$ or $\psi$ is undefined, are excluded.

Once the state of each structure in an ensemble is determined, the number of unique states $n_{\text{states}}(N)$ in a subset of $N$ structures is estimated for both MD and model ensembles. Similar to the estimation of the JS distance (Supplementary Appendix F.2), estimates of $n_{\text{states}}(N)$ are calculated with 100 random subsets for each value of $N$. For MD datasets, the dataset is randomized at the level of trajectories to preserve the temporal ordering of the structures, as was done in the JS distance calculation (see Supplementary Appendix F.2). Then, for a random concatenation of the trajectories in a given MD dataset, subsets of size $N$ are formed by taking the first $N$ structures in this concatenated trajectory. For model datasets, subsets of size $N$ are created by randomly reordering the entire dataset and taking the first $N$ structures in this new ordering.

#### F.5 Discovery of CLN025 Conformational States

To further evaluate Zephyr's ability to generalize beyond its training data, we tested whether it could sample conformational states absent from the training dataset. Specifically, using MD and generated datasets of CLN025, which is known to exhibit both folded and unfolded states,[22] we assessed the ability of the model to generate states unseen during training.

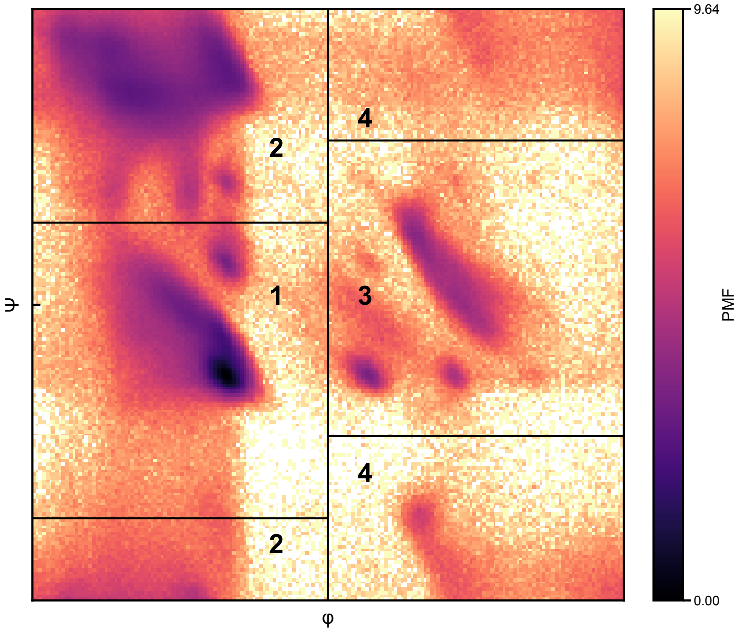

Supplementary Figure 17: **Dihedral state boundaries**
Ramachandran PMF aggregating the backbone dihedral angles of the Top2018 dataset of protein structures.[1] State boundaries are shown as black lines and the delimited regions identify the four possible states, labeled 1–4.

To classify a CLN025 structure as folded or unfolded, we compared it to an NMR ensemble of folded CLN025 structures (PDB 2RVD[28]). Since the topology of the MD CLN025 structures differs from the topology of the PDB 2RVD conformers, missing atoms in the PDB 2RVD termini were added using CHARMM-GUI[45–48] to enable a one-to-one comparison. Using the modified PDB 2RVD conformers, we calculated the heavy-atom RMSD of a given structure to each of the conformers in the PDB 2RVD ensemble. Among all these values, we denote the minimum calculated value as $\mathrm{RMSD_{min}}$. When all frames of the CLN025 MD dataset[22] are analyzed, the resulting distribution of $\mathrm{RMSD_{min}}$ indicates the presence of folded (low $\mathrm{RMSD_{min}}$ values) and unfolded (high $\mathrm{RMSD_{min}}$ values) structures in the dataset (Supplementary Fig. 18a). We define a structure as folded when $\mathrm{RMSD_{min}} \leq 3.0\,\text{Å}$ and unfolded when $\mathrm{RMSD_{min}} > 3.0\,\text{Å}$. Using this definition, the population of the MD ensemble that is folded is 77%.

When Zephyr is trained on the main CLN025 dataset, which includes both folded and unfolded structures, the resulting generated ensemble exhibits a distribution of $\mathrm{RMSD_{min}}$ consistent with the training dataset with similar populations of folded and unfolded states (Supplementary Fig. 18b). The folded fraction is $76.1 \pm 0.2\%$ for the model dataset, $76.2 \pm 4.4\%$[1] for the training dataset and $81.6 \pm 4.4\%$ for the test dataset.

In order to test the ability of the model to produce novel conformational states, we trained Zephyr on datasets consisting of only folded or unfolded structures and assessed its generalizability by counting the number of structures belonging to the state it was not trained on (Supplementary Fig. 18c). In total, five tests were carried out where the training dataset was constructed by filtering structures based on their value of $\mathrm{RMSD_{min}}$.

In the first two tests, Zephyr was trained only on MD structures with $\mathrm{RMSD_{min}} \leq 2.5\,\text{Å}$ or $\mathrm{RMSD_{min}} \leq 3.0\,\text{Å}$. When trained on structures with a $2.5\,\text{Å}$ $\mathrm{RMSD_{min}}$ cutoff, the model was unable to generate unfolded structures. However, with the $3.0\,\text{Å}$ cutoff training dataset, the model generated a non-zero amount of unfolded structures. In this generated ensemble,

[1] The error is estimated by bootstrapping the splitting process of MD datasets with 10000 resamples with replacement.

the structure furthest from the NMR ensemble has a $\mathrm{RMSD_{min}}$ value of 3.8 Å. This structure is visualized with the closest match of the PDB 2RVD ensemble in Supplementary Fig. 18d.

In the other three tests, we assessed the model's ability to generate folded structures when trained only on unfolded structures. Specifically, we trained the model on structures with $\mathrm{RMSD_{min}} \geq 4.0$ Å, $\mathrm{RMSD_{min}} \geq 5.0$ Å or $\mathrm{RMSD_{min}} \geq 6.0$ Å. With both the 4.0 Å and 5.0 Å thresholds, the model was able to generate structures belonging to the folded state. With the 6.0 Å threshold, no structures in the folded state were generated. For the ensemble generated with the model trained using a 4.0 Å $\mathrm{RMSD_{min}}$ cutoff, the structure closest to the NMR ensemble has a $\mathrm{RMSD_{min}}$ value of 1.93 Å and its structure is visualized with the closest PDB 2RVD structure in Supplementary Fig. 18e.

In all five tests, the model exhibits signs of generalizability as quantified by a wider $\mathrm{RMSD_{min}}$ distribution of the model ensembles in comparison with the respective training dataset distribution. Importantly, in some of the tests, some generated structures belong to the state not represented in training. When trained on the unfolded state, the extension of the model $\mathrm{RMSD_{min}}$ distribution is more pronounced than model ensembles trained on the folded state. One reason that may explain this is that the unfolded structures may contain local conformations and contacts that are also present in the folded state. For other proteins, it has been shown that there are similarities between the mean structure of the unfolded state and the folded state[49] and the unfolded state may contain significant native-like structure.[50] Therefore, when trained on unfolded structures, the model is able to learn atomic interactions that are relevant in stabilizing the folded state.

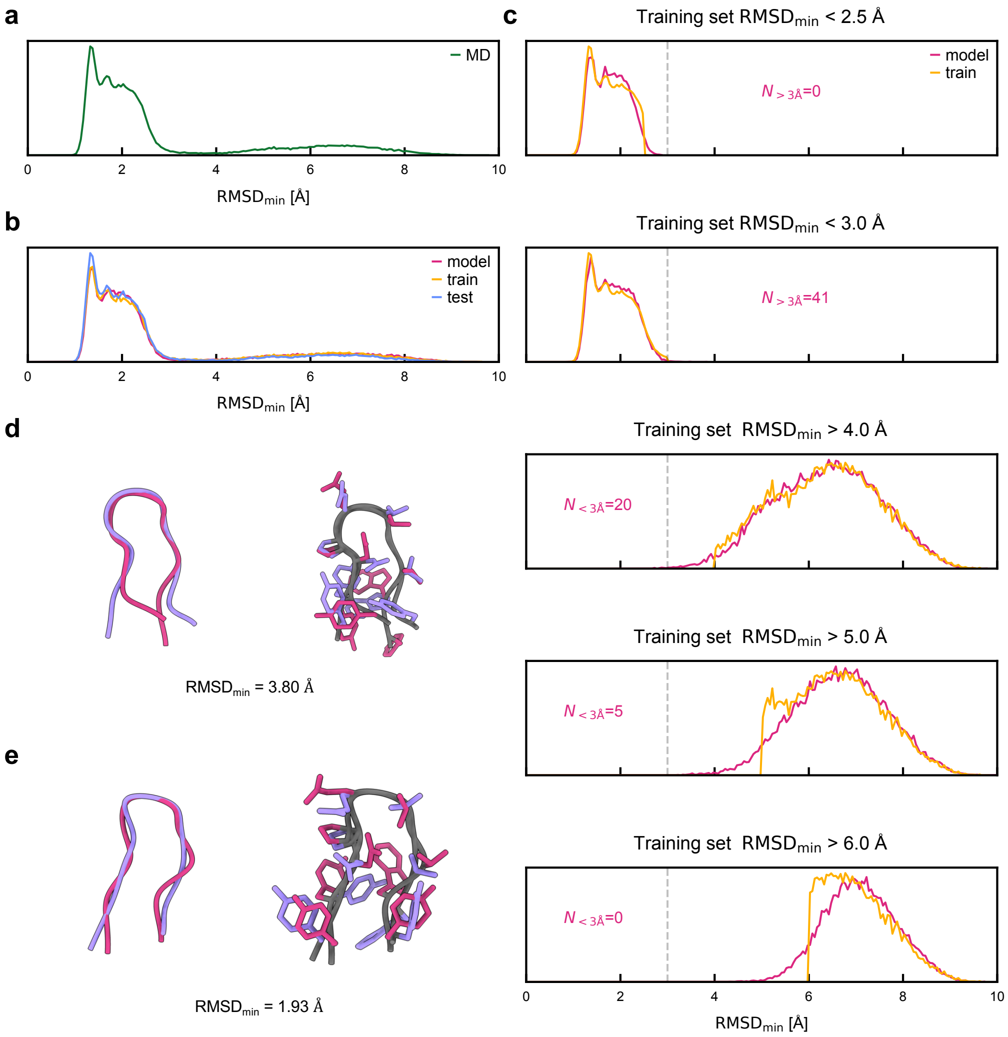

Supplementary Figure 18: **CLN025 experiments**
(**a**) Distribution of $\mathrm{RMSD_{min}}$ of all frames of the CLN025 dataset compared to the 20 conformers of PDB 2RVD. (**b**) Distributions of $\mathrm{RMSD_{min}}$ for the splits of the complete CLN025 dataset. (**c**) Results of training Zephyr on five different subsets of the full CLN025 dataset. The training set used in each case is determined using a threshold value of $\mathrm{RMSD_{min}}$, which is indicated in each subplot title. The model set contains 25,000 structures in each case. The top two plots show examples of training on subsets of the folded state and the lower three plots show examples of training on subsets of the unfolded state. The distribution of $\mathrm{RMSD_{min}}$ of the training set (yellow) and the model set (pink) are shown. The number of generated structures belonging to the state not included in training (either folded or unfolded) is indicated in pink. **(d,e)** Visualizations of generated structures (pink) and their closest conformer in the PDB 2RVD ensemble (purple). Backbone-only (left) and backbone and sidechain (right) structures are visualized using UCSF ChimeraX.[32] **(d)** shows the generated structure with the maximum $\mathrm{RMSD_{min}}$ when the model was trained on only folded structures (**c**, 2nd row). **(e)** shows the generated structure with the minimum $\mathrm{RMSD_{min}}$ when the model was trained on only unfolded structures (**c**, 3rd row).

### G Supplementary References

(1) Williams, C. J.; Richardson, D. C.; Richardson, J. S. The Importance of Residue-level Filtering and the Top2018 Best-parts Dataset of High-quality Protein Residues. *Protein Science* **2022**, *31*, 290–300, DOI: 10.1002/pro.4239.

(2) Gaines, J. C.; Clark, A. H.; Regan, L.; O'Hern, C. S. Packing in Protein Cores. *Journal of Physics: Condensed Matter* **2017**, *29*, 293001, DOI: 10.1088/1361-648X/aa75c2.

(3) Bagci, Z.; Jernigan, R. L.; Bahar, I. Residue Coordination in Proteins Conforms to the Closest Packing of Spheres. *Polymer* **2002**, *43*, 451–459, DOI: 10.1016/S0032-3861(01)00427-X.

(4) Karras, T.; Aittala, M.; Aila, T.; Laine, S. Elucidating the Design Space of Diffusion-Based Generative Models. Advances in Neural Information Processing Systems. 2022; Vol. 35; pp 26565–26577, DOI: 10.52202/068431-1926.

(5) Kingma, D. P.; Ba, J. Adam: A Method for Stochastic Optimization. International Conference on Learning Representations (ICLR). 2015; DOI: 10.48550/arXiv.1412.6980.

(6) Maruyama, G. Continuous Markov Processes and Stochastic Equations. *Rendiconti del Circolo Matematico di Palermo* **1955**, *4*, 48–90, DOI: 10.1007/BF02846028.

(7) Ingraham, J. B.; Baranov, M.; Costello, Z.; Barber, K. W.; Wang, W.; Ismail, A.; Frappier, V.; Lord, D. M.; Ng-Thow-Hing, C.; Van Vlack, E. R.; Tie, S.; Xue, V.; Cowles, S. C.; Leung, A.; Rodrigues, J. V.; Morales-Perez, C. L.; Ayoub, A. M.; Green, R.; Puentes, K.; Oplinger, F.; Panwar, N. V.; Obermeyer, F.; Root, A. R.; Beam, A. L.; Poelwijk, F. J.; Grigoryan, G. Illuminating Protein Space with a Programmable Generative Model. *Nature* **2023**, *623*, 1070–1078, DOI: 10.1038/s41586-023-06728-8.

(8) Petersen, K. B.; Pedersen, M. S. *The Matrix Cookbook*; Technical University of Denmark, 2008.

(9) Paszke, A.; Gross, S.; Massa, F.; Lerer, A.; Bradbury, J.; Chanan, G.; Killeen, T.; Lin, Z.; Gimelshein, N.; Antiga, L.; Desmaison, A.; Kopf, A.; Yang, E.; DeVito, Z.; Raison, M.; Tejani, A.; Chilamkurthy, S.; Steiner, B.; Fang, L.; Bai, J.; Chintala, S. PyTorch: An Imperative Style, High-Performance Deep Learning Library. Advances in Neural Information Processing Systems. 2019; Vol. 32.

(10) Hunter, J. D. Matplotlib: A 2D Graphics Environment. *Computing in Science & Engineering* **2007**, *9*, 90–95, DOI: 10.1109/MCSE.2007.55.

(11) Michaud-Agrawal, N.; Denning, E. J.; Woolf, T. B.; Beckstein, O. MDAnalysis: A Toolkit for the Analysis of Molecular Dynamics Simulations. *Journal of Computational Chemistry* **2011**, *32*, 2319–2327, DOI: 10.1002/jcc.21787.

(12) Gowers, R. J.; Linke, M.; Barnoud, J.; Reddy, T. J. E.; Melo, M. N.; Seyler, S. L.; Domański, J.; Dotson, D. L.; Buchoux, S.; Kenney, I. M.; Beckstein, O. MDAnalysis: A Python Package for the Rapid Analysis of Molecular Dynamics Simulations. *SciPy 2016* **2016**, DOI: 10.25080/Majora-629e541a-00e.

(13) Hagberg, A. A.; Schult, D. A.; Swart, P. J. Exploring Network Structure, Dynamics, and Function Using NetworkX. Proceedings of the 7th Python in Science Conference. 2008; pp 11–15.

(14) Harris, C. R.; Millman, K. J.; van der Walt, S. J.; Gommers, R.; Virtanen, P.; Cournapeau, D.; Wieser, E.; Taylor, J.; Berg, S.; Smith, N. J.; Kern, R.; Picus, M.; Hoyer, S.; van Kerkwijk, M. H.; Brett, M.; Haldane, A.; del Río, J. F.; Wiebe, M.; Peterson, P.; Gérard-Marchant, P.; Sheppard, K.; Reddy, T.; Weckesser, W.; Abbasi, H.; Gohlke, C.; Oliphant, T. E. Array Programming with NumPy. *Nature* **2020**, *585*, 357–362, DOI: 10.1038/s41586-020-2649-2.

(15) Lam, S. K.; Pitrou, A.; Seibert, S. Numba: A LLVM-Based Python JIT Compiler. Proceedings of the Second Workshop on the LLVM Compiler Infrastructure in HPC. 2015; DOI: 10.1145/2833157.2833162.

(16) Eastman, P.; Galvelis, R.; Peláez, R. P.; Abreu, C. R. A.; Farr, S. E.; Gallicchio, E.; Gorenko, A.; Henry, M. M.; Hu, F.; Huang, J.; Krämer, A.; Michel, J.; Mitchell, J. A.; Pande, V. S.; Rodrigues, J. P.; Rodriguez-Guerra, J.; Simmonett, A. C.; Singh, S.; Swails, J.; Turner, P.; Wang, Y.; Zhang, I.; Chodera, J. D.; De Fabritiis, G.; Markland, T. E. OpenMM 8: Molecular Dynamics Simulation with Machine Learning Potentials. *The Journal of Physical Chemistry B* **2024**, *128*, 109–116, DOI: 10.1021/acs.jpcb.3c06662.

(17) Virtanen, P.; Gommers, R.; Oliphant, T. E.; Haberland, M.; Reddy, T.; Cournapeau, D.; Burovski, E.; Peterson, P.; Weckesser, W.; Bright, J.; van der Walt, S. J.; Brett, M.; Wilson, J.; Millman, K. J.; Mayorov, N.; Nelson, A. R. J.; Jones, E.; Kern, R.; Larson, E.; Carey, C. J.; Polat, İ.; Feng, Y.; Moore, E. W.; VanderPlas, J.; Laxalde, D.; Perktold, J.; Cimrman, R.; Henriksen, I.; Quintero, E. A.; Harris, C. R.; Archibald, A. M.; Ribeiro, A. H.; Pedregosa, F.; van Mulbregt, P.; SciPy 1.0 Contributors SciPy 1.0: Fundamental Algorithms for Scientific Computing in Python. *Nature Methods* **2020**, *17*, 261–272, DOI: 10.1038/s41592-019-0686-2.

(18) McInnes, L.; Healy, J.; Saul, N.; Großberger, L. UMAP: Uniform Manifold Approximation and Projection. *Journal of Open Source Software* **2018**, *3*, 861, DOI: 10.21105/joss.00861.

(19) Abramson, J.; Adler, J.; Dunger, J.; Evans, R.; Green, T.; Pritzel, A.; Ronneberger, O.; Willmore, L.; Ballard, A. J.; Bambrick, J.; Bodenstein, S. W.; Evans, D. A.; Hung, C.-C.; O'Neill, M.; Reiman, D.; Tunyasuvunakool, K.; Wu, Z.; Žemgulytė, A.; Arvaniti, E.; Beattie, C.; Bertolli, O.; Bridgland, A.; Cherepanov, A.; Congreve, M.; Cowen-Rivers, A. I.; Cowie, A.; Figurnov, M.; Fuchs, F. B.; Gladman, H.; Jain, R.; Khan, Y. A.;

Low, C. M. R.; Perlin, K.; Potapenko, A.; Savy, P.; Singh, S.; Stecula, A.; Thillaisundaram, A.; Tong, C.; Yakneen, S.; Zhong, E. D.; Zielinski, M.; Žídek, A.; Bapst, V.; Kohli, P.; Jaderberg, M.; Hassabis, D.; Jumper, J. M. Accurate Structure Prediction of Biomolecular Interactions with AlphaFold 3. *Nature* **2024**, *630*, 493–500, DOI: 10.1038/s41586-024-07487-w.

(20) Thomas, N.; Smidt, T.; Kearnes, S.; Yang, L.; Li, L.; Kohlhoff, K.; Riley, P. Tensor Field Networks: Rotation- and Translation-Equivariant Neural Networks for 3D Point Clouds. arXiv, 2018; DOI: 10.48550/arXiv.1802.08219.

(21) Kondor, R.; Lin, Z.; Trivedi, S. Clebsch–Gordan Nets: A Fully Fourier Space Spherical Convolutional Neural Network. Advances in Neural Information Processing Systems. 2018; Vol. 31; DOI: 10.48550/arXiv.1806.09231.

(22) Lindorff-Larsen, K.; Piana, S.; Dror, R. O.; Shaw, D. E. How Fast-Folding Proteins Fold. *Science* **2011**, *334*, 517–520, DOI: 10.1126/science.1208351.

(23) Xiang, S.; Gapsys, V.; Kim, H.-Y.; Bessonov, S.; Hsiao, H.-H.; Möhlmann, S.; Klaukien, V.; Ficner, R.; Becker, S.; Urlaub, H.; Lührmann, R.; de Groot, B.; Zweckstetter, M. Phosphorylation Drives a Dynamic Switch in Serine/Arginine-Rich Proteins. *Structure* **2013**, *21*, 2162–2174, DOI: 10.1016/j.str.2013.09.014.

(24) Rauscher, S.; Gapsys, V.; Gajda, M. J.; Zweckstetter, M.; de Groot, B. L.; Grubmüller, H. Structural Ensembles of Intrinsically Disordered Proteins Depend Strongly on Force Field: A Comparison to Experiment. *Journal of Chemical Theory and Computation* **2015**, *11*, 5513–5524, DOI: 10.1021/acs.jctc.5b00736.

(25) Huang, J.; Rauscher, S.; Nawrocki, G.; Ran, T.; Feig, M.; de Groot, B. L.; Grubmüller, H.; MacKerell, A. D. CHARMM36m: An Improved Force Field for Folded and Intrinsically Disordered Proteins. *Nature Methods* **2017**, *14*, 71–73, DOI: 10.1038/nmeth.4067.

(26) Huang, J.; MacKerell, A. D., Jr. Induction of Peptide Bond Dipoles Drives Cooperative Helix Formation in the (AAQAA)3 Peptide. *Biophysical Journal* **2014**, *107*, 991–997, DOI: 10.1016/j.bpj.2014.06.038.

(27) Chen, J.; Liu, X.; Chen, J. Atomistic Peptide Folding Simulations Reveal Interplay of Entropy and Long-Range Interactions in Folding Cooperativity. *Scientific Reports* **2018**, *8*, 13668, DOI: 10.1038/s41598-018-32028-7.

(28) Honda, S.; Akiba, T.; Kato, Y. S.; Sawada, Y.; Sekijima, M.; Ishimura, M.; Ooishi, A.; Watanabe, H.; Odahara, T.; Harata, K. Crystal Structure of a Ten-Amino Acid Protein. *Journal of the American Chemical Society* **2008**, *130*, 15327–15331, DOI: 10.1021/ja8030533.

(29) Ng, S. C.; Güttler, T.; Görlich, D. Recapitulation of Selective Nuclear Import and Export with a Perfectly Repeated 12mer GLFG Peptide. *Nature Communications* **2021**, *12*, 4047, DOI: 10.1038/s41467-021-24292-5.

(30) Beck, M.; Hurt, E. The Nuclear Pore Complex: Understanding Its Function through Structural Insight. *Nature Reviews Molecular Cell Biology* **2017**, *18*, 73–89, DOI: 10.1038/nrm.2016.147.

(31) Najbauer, E. E.; Ng, S. C.; Griesinger, C.; Görlich, D.; Andreas, L. B. Atomic Resolution Dynamics of Cohesive Interactions in Phase-Separated Nup98 FG Domains. *Nature Communications* **2022**, *13*, 1494, DOI: 10.1038/s41467-022-28821-8.

(32) Meng, E. C.; Goddard, T. D.; Pettersen, E. F.; Couch, G. S.; Pearson, Z. J.; Morris, J. H.; Ferrin, T. E. UCSF ChimeraX: Tools for Structure Building and Analysis. *Protein Science* **2023**, *32*, e4792, DOI: 10.1002/pro.4792.

(33) Berendsen, H. J. C.; van der Spoel, D.; van Drunen, R. GROMACS: A Message-Passing Parallel Molecular Dynamics Implementation. *Computer Physics Communications* **1995**, *91*, 43–56, DOI: 10.1016/0010-4655(95)00042-E.

(34) Abraham, M. J.; Murtola, T.; Schulz, R.; Páll, S.; Smith, J. C.; Hess, B.; Lindahl, E. GROMACS: High Performance Molecular Simulations through Multi-Level Parallelism from Laptops to Supercomputers. *SoftwareX* **2015**, *1–2*, 19–25, DOI: 10.1016/j.softx.2015.06.001.

(35) MacKerell, Jr., A. D.; Bashford, D.; Bellott, M.; Dunbrack, Jr., R. L.; Evanseck, J. D.; Field, M. J.; Fischer, S.; Gao, J.; Guo, H.; Ha, S.; Joseph-McCarthy, D.; Kuchnir, L.; Kuczera, K.; Lau, F. T. K.; Mattos, C.; Michnick, S.; Ngo, T.; Nguyen, D. T.; Prodhom, B.; Reiher, W. E.; Roux, B.; Schlenkrich, M.; Smith, J. C.; Stote, R.; Straub, J.; Watanabe, M.; Wiórkiewicz-Kuczera, J.; Yin, D.; Karplus, M. All-Atom Empirical Potential for Molecular Modeling and Dynamics Studies of Proteins. *The Journal of Physical Chemistry B* **1998**, *102*, 3586–3616, DOI: 10.1021/jp973084f.

(36) Jorgensen, W. L.; Chandrasekhar, J.; Madura, J. D.; Impey, R. W.; Klein, M. L. Comparison of Simple Potential Functions for Simulating Liquid Water. *The Journal of Chemical Physics* **1983**, *79*, 926–935, DOI: 10.1063/1.445869.

(37) Miyamoto, S.; Kollman, P. A. Settle: An Analytical Version of the SHAKE and RATTLE Algorithm for Rigid Water Models. *Journal of Computational Chemistry* **1992**, *13*, 952–962, DOI: 10.1002/jcc.540130805.

(38) Hess, B.; Bekker, H.; Berendsen, H. J. C.; Fraaije, J. G. E. M. LINCS: A Linear Constraint Solver for Molecular Simulations. *Journal of Computational Chemistry* **1997**, *18*, 1463–1472, DOI: 10.1002/(SICI)1096-987X(199709)18:12<1463::AID-JCC4>3.0.CO;2-H.

(39) Darden, T.; York, D.; Pedersen, L. Particle Mesh Ewald: An N·log(N) Method for Ewald Sums in Large Systems. *The Journal of Chemical Physics* **1993**, *98*, 10089–10092, DOI: 10.1063/1.464397.

(40) Bussi, G.; Donadio, D.; Parrinello, M. Canonical Sampling through Velocity-Rescaling. *The Journal of Chemical Physics* **2007**, *126*, 014101, DOI: 10.1063/1.2408420.

(41) Berendsen, H. J. C.; Postma, J. P. M.; van Gunsteren, W. F.; DiNola, A.; Haak, J. R. Molecular Dynamics with Coupling to an External Bath. *The Journal of Chemical Physics* **1984**, *81*, 3684–3690, DOI: 10.1063/1.448118.

(42) Parrinello, M.; Rahman, A. Polymorphic Transitions in Single Crystals: A New Molecular Dynamics Method. *Journal of Applied Physics* **1981**, *52*, 7182–7190, DOI: 10.1063/1.328693.

(43) Kabsch, W. A Solution for the Best Rotation to Relate Two Sets of Vectors. *Acta Crystallographica Section A: Crystal Physics, Diffraction, Theoretical and General Crystallography* **1976**, *32*, 922–923, DOI: 10.1107/S0567739476001873.

(44) Jumper, J.; Evans, R.; Pritzel, A.; Green, T.; Figurnov, M.; Ronneberger, O.; Tunyasuvunakool, K.; Bates, R.; Žídek, A.; Potapenko, A.; Bridgland, A.; Meyer, C.; Kohl, S. A. A.; Ballard, A. J.; Cowie, A.; Romera-Paredes, B.; Nikolov, S.; Jain, R.; Adler, J.; Back, T.; Petersen, S.; Reiman, D.; Clancy, E.; Zielinski, M.; Steinegger, M.; Pacholska, M.; Berghammer, T.; Bodenstein, S.; Silver, D.; Vinyals, O.; Senior, A. W.; Kavukcuoglu, K.; Kohli, P.; Hassabis, D. Highly Accurate Protein Structure Prediction with AlphaFold. *Nature* **2021**, *596*, 583–589, DOI: 10.1038/s41586-021-03819-2.

(45) Jo, S.; Kim, T.; Iyer, V. G.; Im, W. CHARMM-GUI: A Web-Based Graphical User Interface for CHARMM. *Journal of Computational Chemistry* **2008**, *29*, 1859–1865, DOI: 10.1002/jcc.20945.

(46) Jo, S.; Cheng, X.; Islam, S. M.; Huang, L.; Rui, H.; Zhu, A.; Lee, H. S.; Qi, Y.; Han, W.; Vanommeslaeghe, K.; MacKerell, A. D.; Roux, B.; Im, W. Chapter Eight - CHARMM-GUI PDB Manipulator for Advanced Modeling and Simulations of Proteins Containing

Nonstandard Residues. In *Biomolecular Modelling and Simulations*; Advances in Protein Chemistry and Structural Biology; Academic Press, 2014; Vol. 96; pp 235–265, DOI: 10.1016/bs.apcsb.2014.06.002.

(47) Park, S.-J.; Kern, N.; Brown, T.; Lee, J.; Im, W. CHARMM-GUI PDB Manipulator: Various PDB Structural Modifications for Biomolecular Modeling and Simulation. *Journal of Molecular Biology* **2023**, *435*, 167995, DOI: 10.1016/j.jmb.2023.167995.

(48) Kong, L.; Park, S.-J.; Im, W. CHARMM-GUI PDB Reader and Manipulator: Covalent Ligand Modeling and Simulation. *Journal of Molecular Biology* **2024**, *436*, 168554, DOI: 10.1016/j.jmb.2024.168554.

(49) Zagrovic, B.; Snow, C. D.; Khaliq, S.; Shirts, M. R.; Pande, V. S. Native-like Mean Structure in the Unfolded Ensemble of Small Proteins. *Journal of Molecular Biology* **2002**, *323*, 153–164, DOI: 10.1016/S0022-2836(02)00888-4.

(50) Meng, W.; Shan, B.; Tang, Y.; Raleigh, D. P. Native like Structure in the Unfolded State of the Villin Headpiece Helical Subdomain, an Ultrafast Folding Protein. *Protein Science* **2009**, *18*, 1692–1701, DOI: 10.1002/pro.152.
