## Supplementary Movie Legends for "Generating Structural Ensembles of Disordered Proteins with Diffusion Models"

**Description of Additional Supplementary Files**

**File Name: nup98_12-coarse-structure-denoising-trajectory.mp4**
**Description:** Example of a denoising trajectory for sampling the coarse structure of the Nup98-12 system. Black nodes identify the Cα atom of each residue, and red nodes correspond to the sidechain bead. Cα atoms of adjacent residues along the protein sequence are connected by edges. An edge between a Cα atom and a sidechain bead indicates they belong to the same residue. The plot title updates with each frame and shows the molecule ID, sampling frame number, and noise level in nanometers.

**File Name: nup98_12-fine-structure-denoising-trajectory.mp4**
**Description:** Example of a denoising trajectory for sampling the fine (heavy-atom) structure of the Nup98-12 system. Nodes represent heavy atoms, colored by element: grey for carbon, red for oxygen, and blue for nitrogen. Bonded atoms are connected by edges. The plot title updates with each frame and shows the molecule ID, sampling frame number, and noise level in nanometers.

**File Name: nup98_24-coarse-structure-denoising-trajectory.mp4**
**Description:** Example of a denoising trajectory for sampling the coarse structure of the Nup98-24 system. Black nodes identify the Cα atom of each residue, and red nodes correspond to the sidechain bead. Cα atoms of adjacent residues along the protein sequence are connected by edges. An edge between a Cα atom and a sidechain bead indicates they belong to the same residue. The plot title updates with each frame and shows the molecule ID, sampling frame number, and noise level in nanometers.

**File Name: nup98_24-fine-structure-denoising-trajectory.mp4**
**Description:** Example of a denoising trajectory for sampling the fine (heavy-atom) structure of the Nup98-24 system. Nodes represent heavy atoms, colored by element: grey for carbon, red for oxygen, and blue for nitrogen. Bonded atoms are connected by edges. The plot title updates with each frame and shows the molecule ID, sampling frame number, and noise level in nanometers.

**File Name: CLN025-coarse-structure-denoising-trajectory.mp4**
**Description:** Example of a denoising trajectory for sampling the coarse structure of the CLN025 system. Black nodes identify the Cα atom of each residue, and red nodes correspond to the sidechain bead. Cα atoms of adjacent residues along the protein sequence are connected by edges. An edge between a Cα atom and a sidechain bead indicates they belong to the same residue. The plot title updates with each frame and shows the molecule ID, sampling frame number, and noise level in nanometers.

**File Name: CLN025-fine-structure-denoising-trajectory.mp4**
**Description:** Example of a denoising trajectory for sampling the fine (heavy-atom) structure of the CLN025 system. Nodes represent heavy atoms, colored by element: grey for carbon, red for oxygen, and blue for nitrogen. Bonded atoms are connected by edges. The plot title updates with each frame and shows the molecule ID, sampling frame number, and noise level in nanometers.

**File Name: RS-coarse-structure-denoising-trajectory.mp4**
**Description:** Example of a denoising trajectory for sampling the coarse structure of the RS peptide system. Black nodes identify the Cα atom of each residue, and red nodes correspond to the sidechain bead. Cα atoms of adjacent residues along the protein sequence are connected by edges. An edge between a Cα atom and a sidechain bead indicates they belong to the same residue. The plot title updates with each frame and shows the molecule ID, sampling frame number, and noise level in nanometers.

**File Name: RS-fine-structure-denoising-trajectory.mp4**
**Description:** Example of a denoising trajectory for sampling the fine (heavy-atom) structure of the RS peptide system. Nodes represent heavy atoms, colored by element: grey for carbon, red for oxygen, and blue for nitrogen. Bonded atoms are connected by edges. The plot title updates with each frame and shows the molecule ID, sampling frame number, and noise level in nanometers.

**File Name: AAQAA3-coarse-structure-denoising-trajectory.mp4**
**Description:** Example of a denoising trajectory for sampling the coarse structure of the $(AAQAA)_3$ system. Black nodes identify the Cα atom of each residue, and red nodes correspond to the sidechain bead. Cα atoms of adjacent residues along the protein sequence are connected by edges. An edge between a Cα atom and a sidechain bead indicates they belong to the same residue. The plot title updates with each frame and shows the molecule ID, sampling frame number, and noise level in nanometers.

**File Name: AAQAA3-fine-structure-denoising-trajectory.mp4**
**Description:** Example of a denoising trajectory for sampling the fine (heavy-atom) structure of the $(AAQAA)_3$ system. Nodes represent heavy atoms, colored by element: grey for carbon, red for oxygen, and blue for nitrogen. Bonded atoms are connected by edges. The plot title updates with each frame and shows the molecule ID, sampling frame number, and noise level in nanometers.
